# Carbonara: a SAXS-guided seeding framework for exploring protein solution-state dynamics

**DOI:** 10.64898/2026.07.25.740665

**Authors:** Josh McKeown, Cameron Brown, Arron Bale, Hayden Fisher, Robert P. Rambo, Jonathan W. Essex, Matteo T. Degiacomi, Christopher Prior

## Abstract

Proteins in solution often populate conformational ensembles that differ from the static states captured by crystallography or AI-based structure prediction. Conventional molecular dynamics (MD) simulations often fail to cross the energy barriers separating these states on accessible timescales, and statistical reweighting cannot recover conformations never sampled. Here we present Carbonara, a framework that uses experimental small-angle X-ray scattering (SAXS) data to predict alternative physically plausible protein conformations. Carbonara builds on Wiggle, a standalone C*α*-based SAXS forward model validated against explicit-solvent calculations and experimental benchmarks. Using two case studies, an AI-predicted multi-domain helicase (SMAR-CAL1) and a crystallographic antibody fragment (ChiLob7/4 IgG2), we show how seeding MD simulations from Carbonara conformations enables efficient exploration of solution-state conformational landscapes. In both cases, MD ensembles initiated from available models either fail to match the SAXS data or do so only after discarding nearly all sampled conformations, whereas Carbonara-seeded ensembles reach agreement while retaining the majority of conformations. Our modelling framework provides a route from static structural models of flexible multi-domain proteins and multimeric assemblies to solution-state ensembles.

## 1. Introduction

Protein function arises from the interplay between molecular structure and dynamics [1]. Traditional techniques, such as X-ray crystallography [2], Nuclear Magnetic Resonance (NMR) spectroscopy [3], and cryo-electron microscopy (cryo-EM) [4], have long provided the atomistic snapshots underpinning modern structural biology. More recently, AI-based predictors such as AlphaFold [5] and RoseTTAfold [6], have transformed structure prediction on a proteome-wide scale.

Despite this progress, AI-based models remain constrained by biases in the Protein Data Bank (PDB), which predominantly contains lattice-stabilised crystallographic structures. As a result, these models provide limited sampling of the conformational diversity accessible in solution [7], and frequently struggle with disordered or weakly ordered regions that are central to function [8, 9]. Systematic analyses have revealed that AI models under-represent larger-scale structural rearrangements [10]. Ensemble-oriented extensions such as AlphaFlow [11], P2DFlow [12] and more recently BioEmu [13] substantially increase conformational diversity and in some cases capture large scale domain motions, yet remain constrained by their training distributions. These reflect the same crystallographic and MD-sampled conformational basins that static models occupy and more importantly do not incorporate experimental solution-state data during generation.

Small-angle X-ray scattering (SAXS) provides an experimentally accessible view of protein conformational space in solution [14]. It is applicable across a broad molecular-weight range [15, 16], accommodates flexible and partially disordered proteins [17], and has illuminated functional mechanisms in systems as diverse as dengue-virus replication proteins [18], apolipoproteins [19], and lipid-metabolism enzymes [20]. SAXS captures ensemble-averaged information from proteins in near-native solution conditions, making it particularly valuable for systems that undergo large-scale conformational changes or occupy multiple states. Crucially, SAXS and static crystallographic or AI-predicted structures [21, 22, 23] are often in disagreement, presenting a fundamental challenge for bridging high-resolution static models and low resolution solution-state ensemble data.

Atomic resolution approaches for incorporating SAXS data into structural modelling almost universally require a starting model, whether to evaluate agreement or generate candidate conformations. Methods that circumvent this dependency, such as *ab initio* envelope reconstruction [24, 25] generate low-resolution representations directly from scattering data but lack the atomic detail required for mechanistic interpretation or downstream simulation. Within atomic resolution approaches, validation tools [26, 27, 28, 29, 30, 31, 32] assess agreement between SAXS profiles and existing models but cannot propose new conformations. Rigid-body modelling tools such as CORAL [33] efficiently optimise domain orientations for systems with well-defined rigid units, yet cannot simultaneously sample backbone flexibility across multiple linker regions while enforcing biochemical restraints such as disulfide bonds or predicted contact maps. The recently proposed SAXS-A-FOLD [34] attempts to generate new backbone conformations guided by SAXS agreement through sequential perturbation of individual torsion angles, but this constrains its ability to achieve large collective rearrangements and currently supports only single chain systems without biochemical restraints. Molecular Dynamics (MD) simulations [35] can in principle access solution-state ensembles, but when the starting structure lies in a conformational basin within a deep energy well, simulations may fail to sample other important conformational states within accessible timescales. A natural response is to introduce SAXS restraints directly into the force field to drive such transitions; however, strong restraints risk prioritising agreement with SAXS at the expense of genuine conformational dynamics, while weaker restraints leave large conformational rearrangements slow and prone to trapping [36, 37].

To enable an expanded exploration of experimentally consistent solution-state conformational basins we require a sampling strategy capable of generating large scale backbone rearrangements that are physically plausible and consistent with known structural constraints, and a sufficiently accurate SAXS forward model which ensures this search is accurately guided by the experimental constraint provided by the SAXS data.

Here we introduce Carbonara, a SAXS-guided conformational seeding framework designed to address this problem directly. Starting from a candidate structure or set of structures, Carbonara adopts a coarse-grained Cα representation to identify physically and topologically consistent collective domain rearrangements inaccessible via local energy minimisation or microsecond-long MD simulations. To guide this exploration using experimental scattering data we developed Wiggle, a Cα based SAXS forward model trained against explicit solvent all-atom SAXS calculations. Rather than converging to a single best-fit structure, an ill-posed problem for solution scattering, Carbonara generates multiple conformational families, providing starting structures for downstream molecular dynamics or ensemble-reweighting analyses. The method supports multiple flexible regions, automated flexibility assignment based on AlphaFold confidence metrics, the inclusion of additional data such as disulfide bonding or FRET data, and is applicable to proteins exhibiting large-scale flexibility, partial disorder, or multiple coexisting conformational states.

We demonstrate the use of Carbonara on two archetypal systems where the initial structures disagree with SAXS measurements: the AI-predicted DNA-repair helicase SMAR-CAL1, where incorrect domain arrangement produces global disagreement with the solution SAXS profile, and the crystallographic ChiLob7/4 IgG2 antibody, whose disulfide-bond architecture constrains domain organisation while lattice packing enforces a compact geometry not representative of solution state conformation. In both cases, MD ensembles generated from the original models fail to reach agreement with the data without extreme reweighting that excludes nearly all sampled conformations, whereas Carbonara-seeded ensembles require only modest reweighting and retain the majority of the MD sampled conformations as solution consistent.

## 2 Theory

Carbonara performs SAXS-guided sampling within a geometrically constrained backbone representation. The resulting conformations provide starting points for downstream ensemble refinement methods such as molecular dynamics or reweighting. Conformational space is explored subject to geometric and topological constraints that restrict sampling to physically plausible backbone geometries. By operating on a coarse-grained Cα representation, in which each parameter controls multiple residues, Carbonara can efficiently generate collective domain-scale rearrangements that would require many coordinated torsional changes in an all-atom model. This reduced representation enables rapid exploration of large-scale conformational basins while maintaining realistic backbone geometry. As a result, Carbonara can bypass the energy barriers that limit conventional molecular dynamics while retaining sufficient structural realism for downstream all-atom refinement.

To achieve this, Carbonara integrates a validated coarse-grained backbone representation for sampling physically restrained conformations, together with a tertiary and quaternary configuration landscape search [38, 39]. This representation is coupled with Wiggle, a coarse-grained scattering model trained against explicit-solvent WAXSiS-derived scattering profiles. The pipeline further incorporates user-controlled q-range fitting, optional structural constraints, such as disulfide bonds, and back-mapping to all-atom models. Key elements of the method are summarised here; full details are provided in the Methods and Supplementary Information.

### 2.1 Identifying flexible segments

Carbonara takes protein structures as input, composed of one or more subunits together with an experimental SAXS profile. Each input structure is reduced to a coarse-grained representation defined by the positions of its Cα atoms, forming a backbone chain that is segmented into secondary-structure elements, α-helices, β-sheets, and linker regions (Fig. 1a). Conformational sampling is achieved by selectively modifying linker regions, which serve as the primary sources of structural flexibility during refinement.

**Figure 1:**
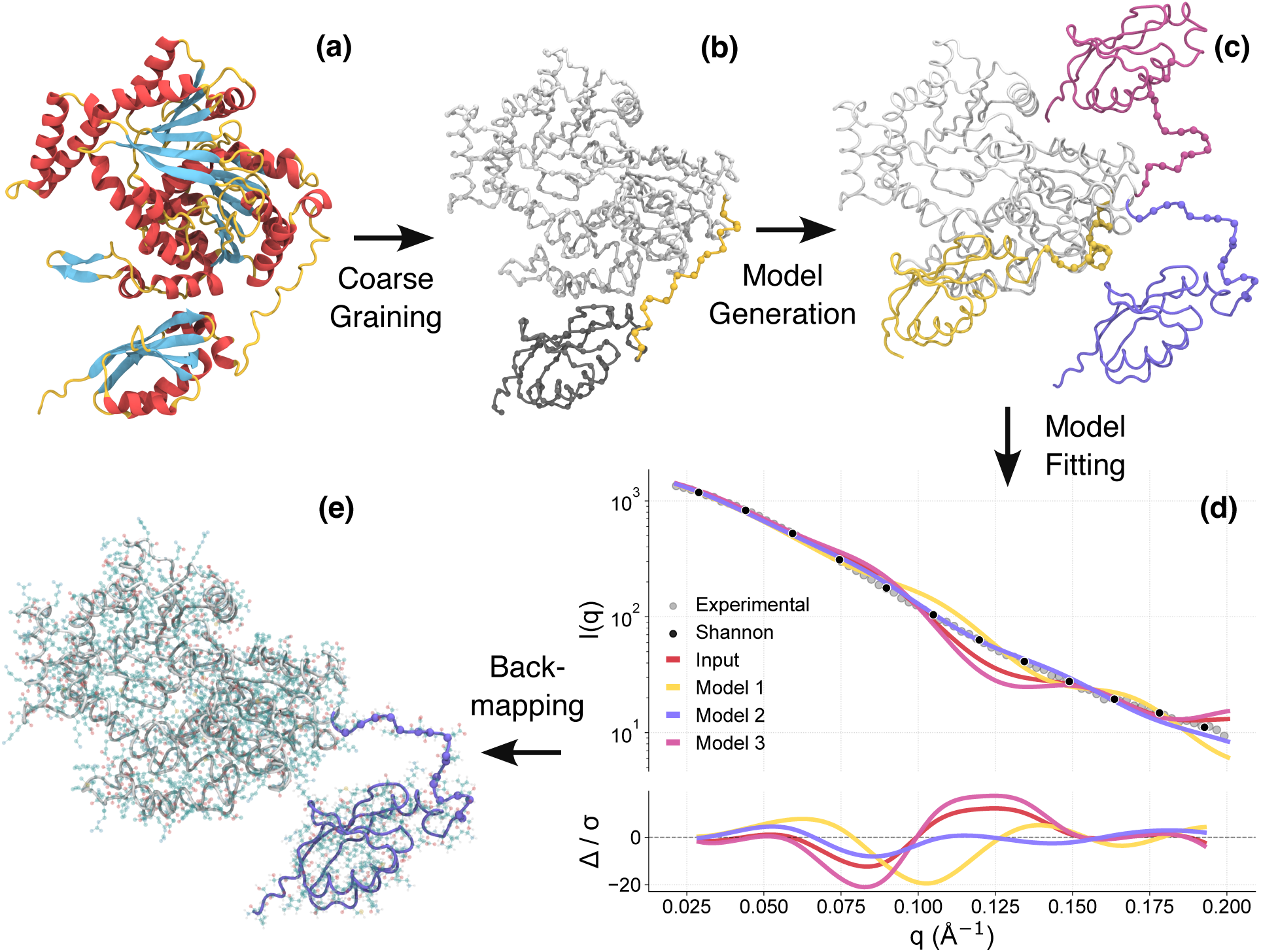
Illustration of the Carbonara modelling process. (a) the secondary structure of the input protein structure is calculated (sheets in blue, helices in red, coils in yellow). (b) the coordinates of Cα atoms are extracted, and linker sections (an example shown in yellow) identified. (c) new protein conformers are generated by altering the linker section so as to improve the fit with SAXS data, subject to statistical and user-defined constraints. (d) the models yielding the best fit, here “Model 2”, are selected (e) the back-mapped all-atomic coordinates are superimposed onto the protein Cα trace.

The set of linker segments permitted to vary can be determined automatically or specified by the user. Carbonara provides three complementary strategies for defining molecular flexibility: (i) assignment based on AlphaFold predicted aligned error (PAE) scores; (ii) an automated backbone-based procedure that identifies flexible regions while preserving existing inter–β-sheet contacts present in the initial AI-derived or crystallographic model; or (iii) explicit user-defined flexibility, which can be guided by interactive three-dimensional visualisation of the structure or by inspection of the secondary-structure sequence. The resulting flexibility assignment can be directly accepted or further edited according to user preference prior to refinement.

### 2.2 Coarse-grained conformational sampling

To generate new prospective configurations with a realistic protein architecture, the flexible segments identified in Section 2.1 are modified using a coarse-grained geometric representation defined by the curvature κ, describing the local bending of the backbone, and the torsion τ, describing its helical twist (Fig. S1). Carbonara constructs new Cα traces for these segments by sampling (κ, τ) values from empirically derived probability distributions obtained from a large database of experimentally determined structures [38, 39] (Fig. S2). A more detailed description of this model is given in Supplementary Methods 1.1. The three principal density peaks of these distributions correspond to the dominant Ramachandran regions, meaning that this coarse-grained representation preserves the local geometric constraints imposed by backbone dihedral preferences Fig. 1(a–c). For multimeric structures, rigid-body rotations of individual subunits may also be applied. The same conformational sampling procedure is used in option (ii) of the initial flexibility-selection process to test whether modifying a segment would disrupt existing β-sheet contacts. Additional structural information, such as disulfide bonds, NMR restraints, or FRET distances, can be incorporated by constraining the separation of user-defined Cα–Cα pairs.

Conformation plausibility of these generated structures is graded using a combination of an amino acid overlap metric, a distance constraint metric, and a fold complexity metric based on the empirically derived strict lower bound of the writhe of the curve’s backbone as a function of the size of the molecule [39]. The full mathematical form of each penalty is given in Methods Sections 8.2.2–8.2.4.

### 2.3 Coarse-grained SAXS modelling with Wiggle

Carbonara requires rapid SAXS profile evaluation across large numbers of candidate Cα-trace conformations. To enable this, it employs Wiggle, a standalone coarse-grained forward scattering model that predicts solution SAXS curves directly from backbone-only representations, prioritising the global size and domain organisation that dominate the low-q SAXS signal. Wiggle is trained against explicit-solvent WAXSiS scattering profiles, and validated on a held-out dataset (see methods) and experimental round-robin benchmark testing against established all-atom SAXS curve generation methods. This anchors the coarse-grained representation to an atomistic description of solution scattering. Details of the training procedure are given in Methods Section 8.1.

Wiggle incorporates three key elements. First, each amino acid (excluding glycine and alanine) is represented by two scattering centres: one at the Cα position and one representing the residue-specific side-chain centre of mass, placed geometrically, based on local backbone curvature. This two-centre representation substantially reduces phase errors relative to Cα-only models while remaining computationally efficient. Second, solvation effects are implicitly incorporated through residue-specific hydrated form factors, optimized subject to smoothness constraints. This captures the dominant solvent contributions to SAXS without explicit solvent modelling. Third, Wiggle is parameterised for q ≤ 0.2 Å*^−^*^1^, where SAXS primarily reports on domain-level and tertiary rearrangements. Within this range Wiggle achieves accuracy comparable to established all-atom scattering models (Supplementary Results). Higher-q features reflecting finer structural detail are intentionally not enforced during Carbonara’s coarse-grained search and are instead resolved during downstream all-atom refinement.

### 2.4 Ensemble-oriented SAXS fitting protocol

Carbonara is designed to determine whether structural heterogeneity is required by the SAXS data, rather than to converge toward a single nominally optimal structure. This distinction is critical for flexible or multi-domain proteins, for which solution scattering reflects an ensemble average where multiple conformational states may coexist. Refinement is performed through batches of independent stochastic runs (by default 20 parallel processes), each exploring physically admissible conformational space under identical experimental constraints. The resulting seed set refers to the collective outcome of these independent refinements, rather than to a time-evolved trajectory or a single structure optimised against an ensemble-averaged objective. When the SAXS data support structural heterogeneity, diversity emerges reproducibly across runs; when they do not, independent runs converge toward similar solutions despite permissive sampling.

By default each refinement optimises a single structure, with low-q SAXS tightly constraining overall size and shape such that independent solutions are expected to cluster in radius of gyration if a dominant solution-state exists. Optionally, Carbonara allows individual refinements to optimise mixtures of structures, with population weights refined alongside geometry, explicitly encoding the hypothesis that the protein populates multiple conformational states with differing compactness.

A central design choice underpinning this behaviour is Carbonara’s treatment of SAXS intensity scaling. Rather than re-optimising the intensity scale by least-squares fitting over the full data range for each trial structure, Carbonara fixes the scale using a restricted low-q window (q ≤ 0.1 Å^−1^). This prioritises agreement at the largest length scales, ensuring that overall size and domain organisation are corrected before finer structural detail is considered. Fixing the scale removes a degree of freedom from the optimisation, such that improvements in fit must arise from genuine structural rearrangements rather than from rescaling alone. This is particularly important when starting models lie far from the solution-state ensemble.

Within the validated operating range of the Wiggle scattering model (q ≤ 0.1 Å^−1^), users may further select the upper fitting limit 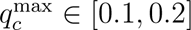 to control the spatial scales emphasised during refinement. Restricting the fit to lower q values promotes broader sampling of global shape and domain-level motions, while extending the range increasingly constrains refinement toward more localised features, at the cost of reduced conformational diversity.

### 2.5 All-atom reconstruction

Carbonara generates Cα-only conformations, which are converted to full all-atom models using established backbone-to-all-atom reconstruction methods for SAXS evaluation and molecular dynamics simulations. The reconstruction procedure and its fidelity are described in Methods and assessed explicitly in Results.

## 3 Results

### 3.1 Wiggle provides accurate coarse-grained SAXS evaluation

Carbonara’s conformational search relies on coarse-grained SAXS profile calculations that are sufficiently accurate for reproduction of experimental scattering profiles and are able to discriminate between distinct conformational states. We start by characterising the accuracy of our method before applying it within the overall Carbonara pipeline. On held-out structures from the validation set (Supplementary Methods 1.2), Wiggle achieved a mean absolute percentage error (MAPE) of 3.30% (compared to the WAXSiS scattering profiles). Applied to five proteins from a SAXS round-robin benchmark [40], comprising 171 measurements collected across 12 synchrotron facilities, none of which were included in training, Wiggle achieved MAPE values of 2.0–6.3% across q ≤ 0.2 Å^-1^, comparable to CRYSOL (1.0– 6.6%), WAXSiS (1.2–3.9%), Pepsi-SAXS (0.5–5.4%), and FoXS (0.5–3.6%) (Supplementary Table S2).

To assess conformational discrimination, we selected twenty pairs of crystal structures representing identical sequences in different conformations (Supplementary Methods 1.2). The paired structures had a mutual difference in radius of gyration ΔR*_g_* = 0.5–8.5 Å and comparative TM-scores 0.37–0.93, reflecting variations in global compactness ranging from nearly indistinguishable conformations to substantially more expanded versus compact states, and spanning structural similarity from close agreement to markedly different folds. Synthetic mixture profiles generated using WAXSiS at population ratios of 20:80, 50:50 and 70:30 were recovered with good accuracy across a structurally diverse test set (Supplementary Table S3, Fig. S7). Performance was consistently strong when the two states differ substantially in global size or shape, while larger deviations arise primarily for pairs with similar radii of gyration and high structural similarity, where the SAXS signal provides limited discriminatory power. Against experimental data, previous benchmarks identified calmodulin as a two-state ensemble comprising 70% open and 30% closed conformations (χ^2^ = 0.79) [41]. Wiggle recovered 75:25 (χ^2^ = 2.24, Fig. 2a). For bovine serum albumin (BSA), where a MultiFoXS analysis identified a three-conformer ensemble with population weights of 74%, 8%, and 18% [41] (χ^2^ = 0.82), Wiggle recovered 70%, 11%, and 19% respectively, using the same three conformer ensemble (χ^2^ = 1.03 Fig. 2b) in close agreement with the published values.

**Figure 2:**
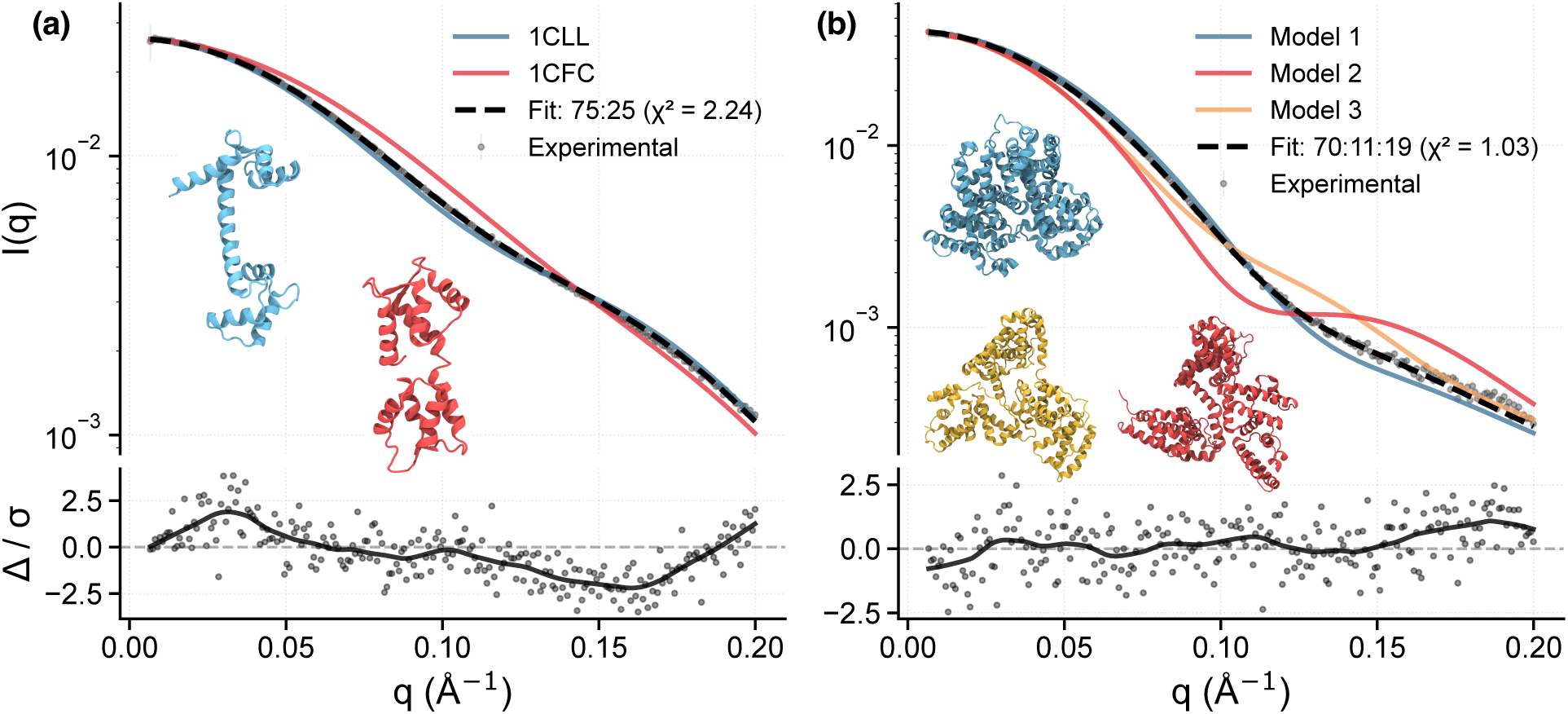
Deconvolution of experimental SAXS datasets. (a) Calmodulin two-state ensemble deconvolution. Experimental SAXS data (gray points) is compared with predictions from open (1CLL, red, R*_g_* = 21.9 Å) and closed (1CFC, blue, R*_g_* = 18.7 Å) crystal structures. Wiggle enables predicting an optimal 75:25 mixture (black dashed line, χ^2^ = 2.24), in close agreement with the experimentally determined 70:30 ratio from SEC-MALS measurements. (b) Bovine serum albumin (BSA) three-state ensemble. Experimental SAXS data (gray points) fitted with three MultiFoXS-optimised conformers, seeded from the PDB 4F5S (chain 1), of increasing radius of gyration (Model 1: R*_g_* = 26.4 Å, red; Model 2: R*_g_* = 29.8 Å, blue; Model 3: R*_g_* = 30.4 Å, orange). Our model predicts population weights of 70:11:19 (black dashed line, χ^2^ = 1.03), consistent with MultiFoXS predictions of 74:8:18. Both methods identify the most compact conformer as the dominant species.

Together, these results demonstrate that Wiggle’s computationally efficient coarse-grained representation captures the key structural determinants required for accurate SAXS prediction and robust discrimination of protein conformational states. Wiggle is designed to capture the SAXS features most sensitive to global conformation (q ≲ 0.2 Å*^−^*^1^). Higher-q features, where hydration-shell heterogeneity becomes important, are subsequently evaluated using all-atom scattering models during downstream refinement.

### 3.2 Carbonara reveals an experimentally consistent conformational ensemble of the SMARCAL1 ATPase

Human SMARCAL1 is a DNA-dependent ATPase involved in replication fork stabilisation. Its catalytic domain (SMARCAL1*^CD^*, residues 325–870) comprises an N-terminal HARP2 domain and a C-terminal bilobed ATPase motor domain. Previous SAXS studies have shown that SMARCAL1*^CD^* undergoes large-scale conformational rearrangements during its functional cycle [42], making it a stringent test for SAXS-guided conformational sampling. An AlphaFold3-predicted model of SMARCAL1*^CD^* showed poor agreement with experimental SAXS data (χ^2^ = 15.06, FoXS), with pronounced deviations at low q, indicating substantial large scale disagreement with the solution-state ensemble. We therefore applied Carbonara, assigning flexibility to the linker regions connecting the HARP2 and ATPase domains (see Methods).

### 3.2.1 Carbonara consistently produces high-quality atomistic predictions capturing protein flexibility

To evaluate robustness, efficiency, and structural diversity of the Carbonara seed set, we performed 100 independent Carbonara refinement runs in five batches of 20 parallel processes. All runs substantially improved SAXS agreement relative to the AlphaFold3 model, with all-atom back-mapped structures achieving mean χ^2^ values of 1.4–1.6 (FoXS) and 2.1–2.3 (CRYSOL) across all batches (Supplementary Data 2.1), confirming reliable convergence without manual selection. Median χ^2^ values fell below 2 within approximately 10–20 minutes of sampling and continued sampling produced additional acceptable conformations rather than collapsing onto a single structure (Figure S12).

Despite comparable SAXS agreement, the resulting seed set exhibited substantial structural diversity. Relative to the AlphaFold3 model, Carbonara predictions showed large deviations (RMSD ∼ 25 Å, TM-score ∼ 0.10), consistent with major domain rearrangements (Fig. S16). Pairwise comparisons within the seed set spanned RMSD values of approximately 3–30 Å with higher TM-scores up to ∼ 0.94, indicating distributed moderate-amplitude rearrangements rather than a single dominant conformational transition (Fig. S16).

To place the Carbonara ensemble in context, we compared its structural variability with predictions generated using SAXS-A-FOLD and CORAL, both run using their respective default workflows (see Methods Section 8.6). All three approaches produced models consistent with the experimental scattering data. CORAL achieved mean values of χ^2^ = 0.668 over the full data range and χ^2^ = 1.028 over q ∈ [0, 0.2], while the SAXS-A-FOLD ensemble fit gave χ^2^ = 1.09. Whereas SAXS-A-FOLD yielded structures closely clustered around the initial AlphaFold model and CORAL primarily generated alternative rigid-body arrangements, Carbonara sampled a substantially broader conformational space while maintaining backbone continuity. In addition, the CORAL model proposes a HARP2 domain that is extended away from the ATPase, a model inconsistent with limited proteolysis studies which demonstrated the HARP2-ATPase linker region is resistant to proteolytic digestion [43] (Fig. S18). This expanded structural dispersion, reflected in a significantly wider range of RMSD and TM-score distributions (see Supplementary Results 2.1.2 and Fig. S10), indicates that Carbonara captures a greater extent of conformational flexibility compatible with the SAXS data, rather than converging on a small number of discrete rearrangements.

#### 3.2.2 Carbonara-seeded MD better reflects SMARCAL1’s solution-state dynamics

To test whether these structures provide improved starting points for generating MD ensembles, we ran a new batch run of the Carbonara algorithm in default mode (20 predictive runs). This is distinct from the 5 batches analysed in the previous section (which were run to test consistency of the algorithm) and represents the suggested default use of the pipeline for the purpose of seeding MD simulations. We initiated 300 ns MD simulations from each of 20 Carbonara-derived conformations (6 µs total) and compared them to three independent 2 µs simulations seeded from the AlphaFold3 model. Principal component analysis (PCA) on aligned Cα coordinates revealed separation between the two MD ensembles (Fig. 3a), with structures exhibiting low SAXS χ^2^ values clustered exclusively within the Carbonara-seeded region (all χ^2^ values were evaluated on the full data range q ∈ [0, 0.25]). SAXS-A-FOLD predictions fell near the AlphaFold seeded region and CORAL predictions overlapped partially with Carbonara but spanned only a restricted subset of the conformational space.

**Figure 3:**
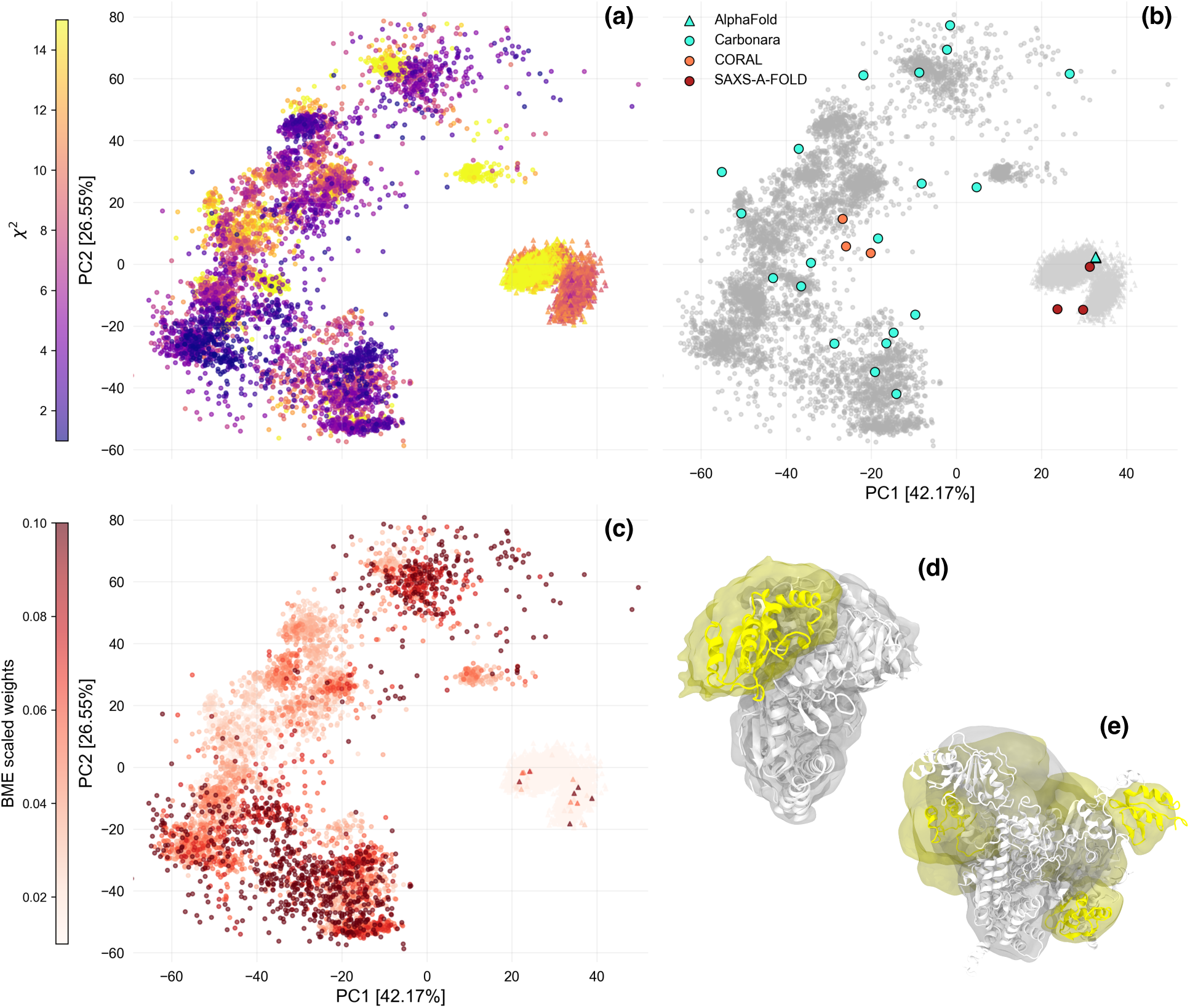
Conformational space exploration and subdomain positioning of Human SMARCAL1. Throughout, AlphaFold-seeded ensemble shown as triangles and Carbonara-seeded ensemble as circles. (a) Principal component analysis (PCA) projection of 12,000 MD snapshots (1 ns intervals) coloured by χ^2^ fit (FoXS, q_max_ = 0.2 Å*^−^*^1^), colouring capped at χ^2^ ∈ [1, 15]. PC1 and PC2 capture 42.17% and 26.55% of explained variance respectively; alignment performed on Cα traces over residues 400–480. (b) The same PCA projection, trajectories in light gray, teal markers denote the starting structures. SAXS-A-FOLD predictions shown in red circles and CORAL orange circles. (c) PCA projection coloured by BME weights (normalised to [0, 1] per ensemble) with colour capped at 0.1, darker shades indicate higher weighting. (d,e) BME-weighted spatial density of the AlphaFold-seeded (d) and Carbonara-seeded (e) ensembles, with the HARP2 domain shaded yellow and the ATPase domains in white. Sample structures are rendered inside density. The AlphaFold-seeded ensemble shows a tightly localised HARP2 density, while the Carbonara-seeded ensemble reveals a broad distribution of HARP2 positions consistent with the extensive domain rearrangements identified by PCA.

The AlphaFold-seeded MD ensemble showed a narrow R*_g_* distribution well below the experimental Guinier value, whereas the Carbonara-seeded MD ensemble sampled a substantially broader range approaching it (Table 1). Inter-domain geometry confirmed these differences, with AlphaFold-seeded simulations showing restricted ATPase-N to ATPase-C distances (31.7 ± 3.0 Å) and inter-domain angles (85.7*^◦^* ± 4.4*^◦^*) while Carbonara-seeded simulations explored substantially broader ranges (41.1 ± 6.9 Å and 66.8*^◦^* ± 25.2*^◦^*) reflecting extensive HARP2 repositioning and ATPase lobe flexibility (Fig. 3b,c).

**Table 1:** Comparison of MD ensembles seeded from original models versus Carbonara conformations. For each system, the table reports the SAXS fit quality before and after Bayesian Maximum Entropy (BME) reweighting, the BME regularisation parameter θ, the effective ensemble size N_eff_, and the unweighted and reweighted mean radius of gyration (R*_g_*). Experimental Guinier R*_g_* values are shown for reference. All χ^2^ values were evaluated using FoXS.

|  | SMARCAL1 |  | ChiLob7/4 IgG2 |  |
| --- | --- | --- | --- | --- |
|  | AlphaFold | Carbonara | Crystal | Carbonara |
| Unweighted $\chi^2$ | 9.22 | 3.07 | 3.44 | 1.56 |
| BME-reweighted $\chi^2$ | 4.96 | 0.54 | 1.78 | 1.50 |
| BME $\theta$ | 80 | 70 | 15 | 65 |
| $N_{\text{eff}}$ | 0.02 | 0.60 | 0.02 | 0.95 |
| Unweighted $R_g$ (Å) | 26.6 | 28.9 | 37.8 | 40.1 |
| Reweighted $R_g$ (Å) | 27.4 | 30.0 | 37.8 | 40.3 |
| Guinier $R_g$ (Å) | $31.9 \pm 0.3$ | | $40.6 \pm 0.5$ | |

Bayesian Maximum Entropy (BME) reweighting further highlighted these differences [44] (Table 1, Fig. 4, a,b). The AlphaFold-seeded ensemble showed poor agreement with uniform weighting, and even aggressive reweighting failed to achieve an acceptable fit, maintaining pronounced deviations at low q. In contrast, the Carbonara-seeded ensemble reached statistical agreement with experiment under modest reweighting, with the R*_g_* distribution spanning up to 35.2 Å encompassing the experimental value. The N_eff_ vs χ^2^ curves make the fundamental difference between the two ensembles clear (Fig. 4, d). For the AlphaFold-seeded MD ensemble, χ^2^ decreases continuously as structures are excluded, never reaching a plateau and still failing to achieve acceptable agreement even when 98% of the ensemble is disregarded. This behaviour indicates that the ensemble does not contain experimentally relevant conformations. Reweighting cannot recover states that were never sampled. The Carbonara-seeded ensemble shows a different pattern, with a plateau at χ^2^ ≈ 0.5 for N_eff_ < 0.6 indicating that relevant states are present rather than overfit, though we note χ^2^ < 1 may reflect conservative experimental uncertainty estimates. The BME weights, projected onto the PCA landscape (Fig. 3c), reveal that reweighting does not collapse the ensemble onto a single conformational region. Instead two preferentially weighted clusters emerge in PC space, with a connecting region modestly yet not entirely down-weighted.

**Figure 4:**
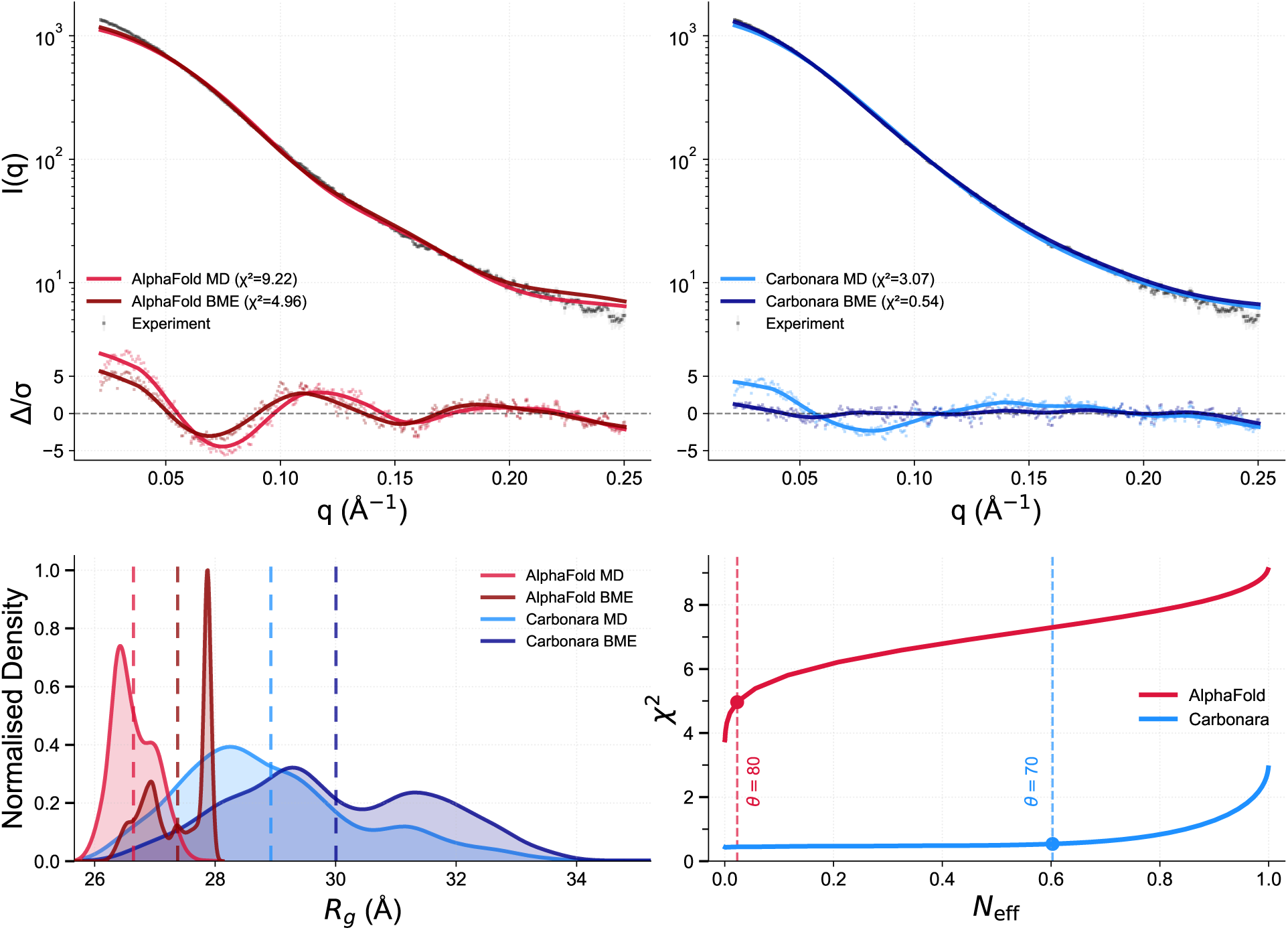
Bayesian/Maximum Entropy (BME) reweighting of SMARCAL1*^CD^* ensembles. (a, b) SAXS intensity profiles and error weighted residuals (Δ/σ) for AlphaFold-seeded (a, red) and Carbonara-seeded ensembles (b, blue). Light lines show unweighted MD ensemble averaged theoretical SAXS, darker lines show BME-reweighted profiles, gray points show experimental data with error bars. All theoretical SAXS profiles were calculated up to q = 0.25Å*^−^*^1^ with FoXS. Unweighted fits gave χ^2^ = 9.22 (AlphaFold) and χ^2^ = 3.07 (Carbonara), while reweighting achieved χ^2^ = 4.96 and χ^2^ = 0.54 for AlphaFold and Carbonara ensembles respectively. Regularisation θ = 80 for AlphaFold and θ = 70 for Carbonara, resulting ensemble sample sizes (N_eff_) of 2% (AlphaFold) and 60% (Carbonara). Despite extreme reweighting, AlphaFold residuals exceed ±5σ particularly at low-q, while Carbonara residuals remain within ±1.6σ across the entire q range. (c) Radius of gyration (R*_g_*) distributions, lighter shades show unweighted MD and darker shades show BME-reweighted. AlphaFold (red) shows narrow distribution centred at R*_g_* = 26.6 Å and is shifted to R*_g_* = 27.4 Å upon aggressive reweighting. Carbonara (blue) shows broader distribution centred at R*_g_* = 28.9 Å and is shifted to R*_g_* = 30.0 Å with minimal reweighting. (d) N_eff_ against χ^2^ as BME regularisation (θ) varies. AlphaFold curve (red) exhibits continuous deterioration without plateau. Carbonara (blue) demonstrates a rapid initial improvement followed by plateau at χ^2^ ≈ 0.5 below N_eff_ = 0.6.

### 3.3 Refinement of a crystal structure of ChiLob7/4 IgG2 constrained by disulfide bond distances

The C239S variant of ChiLob7/4 IgG2 F(ab)_2_ presents a complementary challenge to SMARCAL1*^CD^*, representing a high-resolution multimeric crystal structure that is locally accurate, but globally misrepresents the solution conformation. This variant is notable for its CD40 agonistic activity [45] and its crystal structure (PDB: 6TKD, 1.90 Å resolution) fails to reflect the solution scattering (χ^2^ = 7.96, FoXS), with radius of gyration R*_g_* = 36.4 Å substantially smaller than the experimental Guinier value R*_g_* = 40.6 ± 0.52 Å [33]. Previous MD studies initiated from the crystal structure failed to reach conformations consistent with experimental SAXS data [45]. Carbonara was applied with flexibility assigned to the inter-F(ab) hinge region, constraining the disulfide bonding pattern characteristic of the C239S variant while preserving intra-F(ab) domain structures as rigid bodies (see Methods Section 8.5.2).

#### 3.3.1 Carbonara generates a SAXS-consistent seed ensemble without artificial diversity

Across 100 independent refinement runs (five batches of 20) and multiple fitting ranges 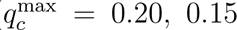, and 0.10), Carbonara consistently improved agreement with the experimental SAXS data relative to the starting crystal structure when evaluated using both FoXS and CRYSOL (Supplementary Data 2.2). The improvement is most pronounced at low q, where the FoXS χ^2^ value decreases from 9.57 to a mean of 1.86 over q ∈ [0, 0.1], indicating that the dominant discrepancy in the crystallographic model arises from global domain arrangement rather than local structural detail. Despite the larger size of the ChiLob7/4 assembly, high-quality SAXS-consistent predictions appear on similar timescales to the SMARCAL1*^CD^* case (Fig. S15), reflecting the reduced conformational search space associated with its largely rigid architecture.

In contrast to the flexible SMARCAL1*^CD^* system, pairwise comparisons between refined ChiLob7/4 models reveal only limited structural dispersion (RMSD ∼ 0.5–10 Å, TM-score ∼ 0.4–0.98, see Fig. S17). This behaviour demonstrates that Carbonara introduces structural diversity only when supported by the SAXS data, remaining conservative for systems whose solution-state architecture is largely rigid. Refined structures consistently adopted more open geometries than the crystal structure, with mean R*_g_* increasing from 36.4 to 40.3 ± 0.4 Å and hinge angles opening from 116.4*^◦^* to 129.4*^◦^* ± 3.75*^◦^*.

Direct comparison with alternative SAXS-guided modelling approaches was not feasible for this system. Existing methods either do not currently support multimeric assemblies (e.g., SAXS-A-FOLD) or cannot enforce structural constraints such as the disulfide-bond connectivity that defines the C239S IgG2 hinge geometry (e.g., CORAL and SAXS-A-FOLD).

#### 3.3.2 Carbonara-seeded MD captures the solution-state ensemble of ChiLob7/4 IgG2

MD simulations from each of the 20 Carbonara-refined structures (20 × 3 × 100 ns, 6 µs total) were compared with three crystal-seeded 2 µs simulations. Again this represents the suggested use of the Carbonara algorithm in default mode (20 predictive runs) and these structures are distinct from those analysed in the 5 batch runs of the previous section. Theoretical SAXS profiles were calculated for snapshots at 1 ns intervals using FoXS. Crystal-seeded simulations remained tightly clustered around the crystallographic geometry (hinge angle 122.4*^◦^* ± 3.4*^◦^*, dihedral angle 64.2*^◦^* ± 5.0*^◦^*) and failed to sample conformations with R*_g_* > 39.8 Å, despite microsecond-scale sampling. Carbonara-seeded simulations explored substantially broader conformational space (hinge angle 134.8*^◦^*±11.9*^◦^*, dihedral angle 82.3*^◦^*± 25.7*^◦^*) with an R*_g_* distribution in excellent agreement with the experimental Guinier value (Table 1). Individual χ^2^ values exhibit a characteristic U-shaped dependence on R*_g_* with the minimum near R*_g_* ≈ 40–41 Å, with the crystal-seeded ensemble confined to the left-hand arm and Carbonara-seeded ensemble spanning the full width (Fig. 5b).

**Figure 5:**
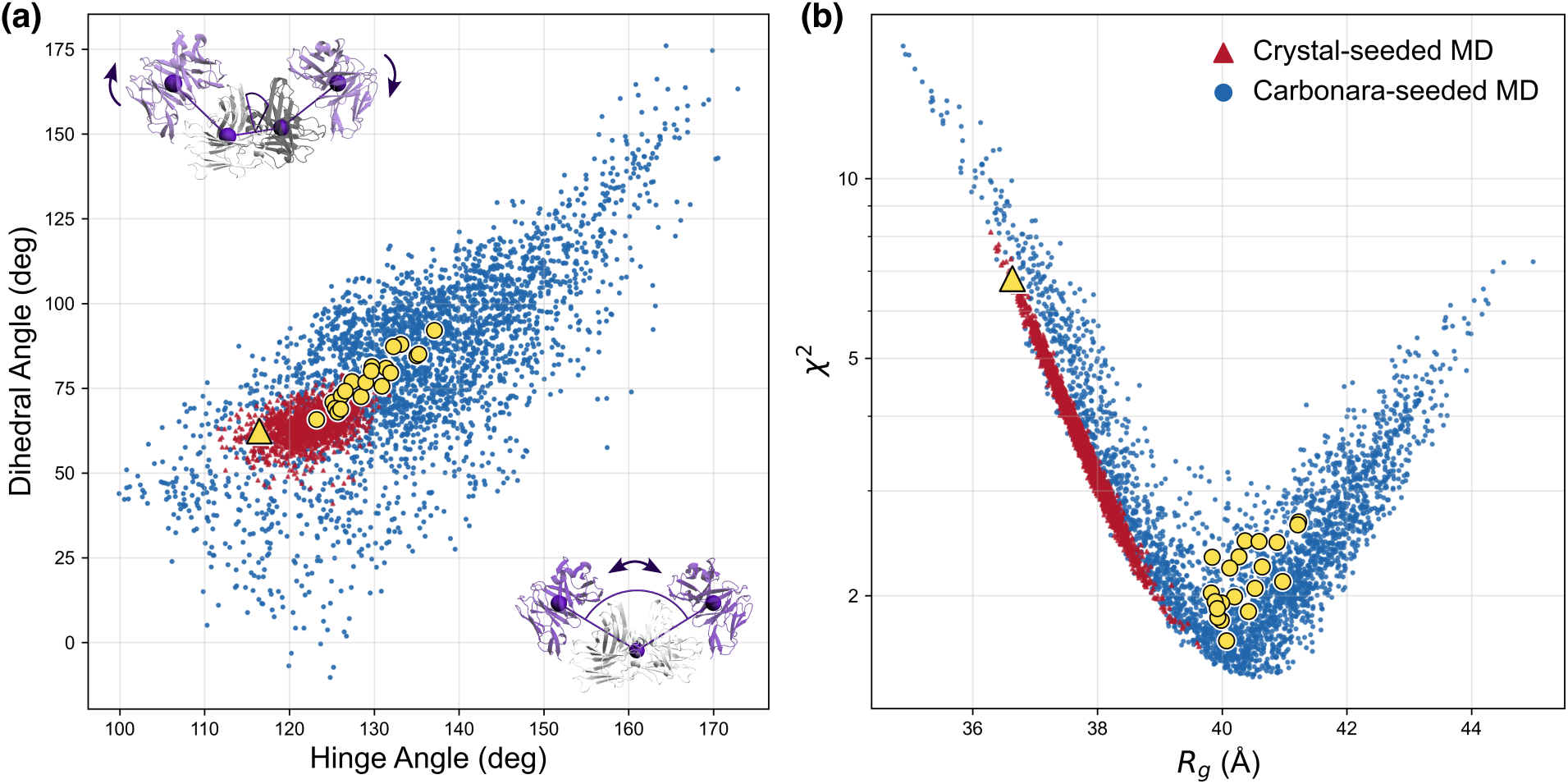
Space explored by crystal and Carbonara-seeded simulations. (a) Inter-domain hinge angle versus dihedral angle for crystal-seeded (red triangles) and Carbonara-seeded (blue circles) MD ensembles. Protein renders illustrate the hinge and dihedral angle definitions and centre-of-mass selections on the corresponding axes. Crystal-seeded simulations cluster tightly near the crystallographic geometry, while Carbonara-seeded simulations explore substantially broader conformational space. The crystal structure (yellow triangle) and 20 Carbonara starting structures (yellow circles) are overlaid. (b) Radius of gyration (R*_g_*) versus χ^2^ (FoXS q_max_ = 0.15 Å*^−^*^1^) for individual MD structures. Individual fits exhibit a characteristic U-shaped dependence on R*_g_*, with the minimum near the experimental Guinier R*_g_* of 40.6 ÅĊ rystal-seeded simulations (red) are confined to the left-hand arm of the profile, sampling only compact conformations that never reach the χ^2^ minimum. Carbonara-seeded simulations (blue) span the full width of the profile. Yellow markers as in (a).

BME reweighting further quantified these differences (Table 1, Fig. 6a,b). The crystal-seeded ensemble reached only moderate agreement by excluding approximately 98% of structures, with persistent systematic deviations at low q. The Carbonara-seeded ensemble requires no such exclusion, with uniform weighting already yielding good agreement and reweighting preserving 95% of structures while leaving residuals evenly distributed across q. The N_eff_ versus χ^2^ curves show continuous deterioration for the crystal ensemble and a plateau at χ^2^ ≈ 1.45 for the Carbonara ensemble (Fig. 6d), indicating that the majority of sampled conformations are SAXS-consistent.

**Figure 6:**
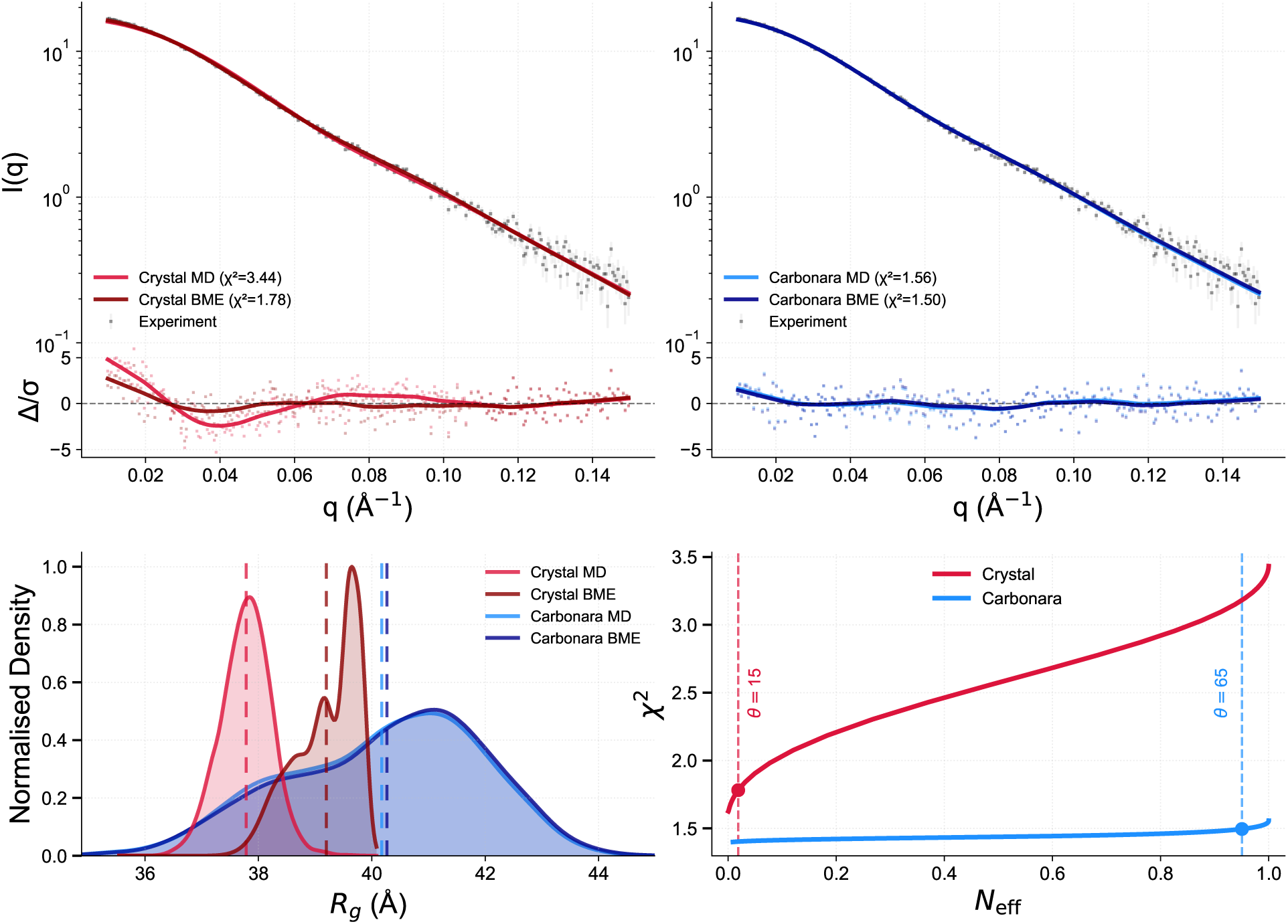
Bayesian Maximum Entropy analysis of C239S IgG2 MD ensembles. (a, b) SAXS intensity profiles and error weighted residuals (Δ/σ) for crystal-seeded (a, red) and Carbonara-seeded (b, blue) MD ensembles evaluated up to q = 0.15 Å*^−^*^1^ (FoXS). Light colours show unweighted MD ensemble averaged SAXS while dark colours show BME-reweighted profiles, gray points show experimental data and error bars. The crystal ensemble, unweighted χ^2^ = 3.44, BME reweighted χ^2^ = 1.78, θ = 15, N_eff_ = 0.02. Carbonara ensemble, unweighted χ^2^ = 1.56, BME reweighted χ^2^ = 1.50, θ = 65, N_eff_ = 0.95. (c) Radius of gyration (R*_g_*) distributions. Light colours show unweighted MD distribution and dark BME reweighted. Dashed vertical lines indicate the distribution means, crystal MD centred at 37.78 Å (light red) and BME reweighted centred at 37.82 Å (dark red). Carbonara MD centred at 40.17 Å (light blue) BME reweighting at 40.3 Å (dark blue). (d) N_eff_ versus χ^2^ as BME regularisation parameter (θ) varies. Crystal ensemble (red) shows elevated χ^2^ across all N_eff_ values that continues to decay with exclusion of structures. The Carbonara ensemble (blue) exhibits a plateau at χ^2^ ≈ 1.45, demonstrating that the majority of sampled conformations are SAXS-consistent.

#### 3.3.3 Carbonara’s mixture refinement can highlight large scale conformational variability

To assess whether the broader conformational range evident in the MD could be recovered directly at the refinement stage, we performed additional Carbonara runs optimising pairs of structures simultaneously with their weighted average required to reproduce the scattering profile. Rather than converging on a single intermediate, the two structures within each pair diverge across the conformational landscape, and the resulting pairs (40 structures) spanned R*_g_* = 34–42 Å (Fig. S19), comparable to the breadth of the MD-sampled distribution rather than a single structure at its mean.

## 4 Discussion and Conclusions

Our case studies illustrate how the choice of starting structure shapes downstream MD results. For SMARCAL1, conformational basins consistent with SAXS data were absent from the microsecond-scale MD simulations seeded via AlphaFold. In contrast, Carbonara generated seeds placed the system directly within these regions of the conformational space. BME reweighting retained 60% of the Carbonara-seeded MD conformations. The resulting ensemble clustered into two dominant conformational subsets that included the Carbonara seeds, but not those suggested by the alternative modelling approaches. These broader HARP2–ATPase domain rearrangements are consistent with the large-scale repositioning required during DNA remodelling and annealing [46], supporting the physical plausibility of the sampled conformational space.

For ChiLob7/4 the relevant solution-state conformational energy basin does exist, but crystal packing renders important sub-states effectively inaccessible to microsecond-scale MD started from the crystallographic structure. Retention of 95% of structures upon reweighting indicates that once Carbonara placed MD in the correct region, virtually the entire MD simulation sampled a solution-consistent distribution. The enforcement of a known disulfide bonding structure reduced the conformational search to physically accessible hinge configurations.

The Carbonara set of seed structures is not intended to represent the final solution-state ensemble. Instead, they position MD simulations within relevant regions of conformational space, where multiple short trajectories can explore local neighbourhoods, rather than relying on rare energy barrier-crossing events from a single starting structure. Triplicate 2 µs simulations were initiated from the AlphaFold and crystallographic models to test whether spontaneous barrier crossing could occur within conventional MD timescales. In contrast, Carbonara-generated seeds were used to initiate multiple shorter trajectories (300 ns each) distributed across the conformational landscape. The fact that the long trajectories from the original structures failed to reach SAXS-consistent states, whereas the shorter Carbonara-seeded simulations succeeded, shows how selecting appropriate starting coordinates can facilitate broader sampling of a protein’s conformational space.

For a flexible system like SMARCAL1, conformational heterogeneity is already apparent at the seed-generation stage. Independent Carbonara runs produce single structures with comparable χ^2^ values that span a broad region of conformation space, directly signalling that the SAXS data supports multiple structural states. The more rigid ChiLob7/4 system presented a potential challenge for the method, where convergence of independent refinements may falsely suggest the presence of a single dominant conformation.

Carbonara’s mixture-refinement strategy addresses this ambiguity directly. For ChiLob7/4, individual Carbonara seeds cluster near the minimum of the characteristic U-shaped R*_g_*–χ^2^ relationship, corresponding to the single structure that best satisfies the scattering pro-file. This minimum lies close to the average R*_g_* of the Carbonara-seeded MD ensemble. However, when pairs of structures were optimised simultaneously, the solutions diverged, spanning the same R*_g_* range sampled by the MD simulations rather than collapsing toward the population mean. This behaviour indicates that the presence and approximate extent of conformational heterogeneity can already be inferred at the seeding stage, before any MD simulations are performed. Full characterisation of the solution-state distribution and its physical basis nevertheless requires downstream MD refinement, which Carbonara is designed to enable by providing seed structures distributed across the relevant regions of conformational space.

We compared Carbonara with two existing SAXS-guided modelling tools, CORAL and SAXS-A-FOLD, both of which are designed to identify conformations that reproduce the scattering profile and can subsequently be used for further structural analysis. While these approaches achieved comparable χ^2^ agreement with the experimental SAXS data, Carbonara explored a substantially broader region of conformational space, as reflected in the wider distributions of RMSD and TM-scores among the resulting models. Importantly, MD simulations seeded from the Carbonara derived structures produced ensembles consistent with experimental data with only modest or no reweighting.

SAXS-driven MD methods address a different stage of the modelling workflow. Rather than searching for candidate conformations, these approaches refine conformational ensembles by applying SAXS-derived restraints during the simulation itself [36]. In this sense they correspond more closely to the second stage of the Carbonara pipeline, where MD simulations explore the conformational landscape starting from SAXS-consistent seed structures. While the classical MD simulations seeded by Carbonara yielded ensembles that already reproduced experimental data, adopting SAXS-driven MD starting from Carbonara-generated seeds may prove useful for systems where additional biasing is required to refine the resulting ensemble.

Carbonara is most naturally suited to proteins where solution-state disagreements arise from the relative positioning of structured elements, whether connected by flexible linker within a single chain or arranged as independent subunits in a multimeric assembly, while those elements themselves retain their folded architecture in solution. Although the framework can resample any designated backbone segment using empirical distributions associated with different secondary-structure types, its strengths lie in repositioning existing structural elements rather than generating entirely disordered conformations. Systems dominated by extensive intrinsically disordered regions may therefore require additional benchmarking, as the behaviour of the sampling framework in such regimes has not yet been systematically evaluated.

As with all SAXS-based modelling approaches, the low information content of solution scattering data means that multiple structurally distinct ensembles may achieve comparable agreement with experiment and uniqueness cannot be guaranteed. Complementary experimental techniques such as FRET spectroscopy [47] or X-ray footprinting [48] can provide orthogonal constraints that reduce solution ambiguity and extend applicability. Carbonara readily supports such information through arbitrary Cα–Cα distance restraints, allowing experimentally derived distance measurements to be incorporated directly during conformational sampling to generate ensembles that more faithfully reflect the conformational states populated in solution.

Wiggle, the coarse-grained SAXS forward model developed for Carbonara, addresses a broader methodological need arising from the increasing use of backbone-level representations in modern protein modelling. Recent AI-driven protein design and generative modelling approaches frequently operate directly on protein backbones rather than full atomic coordinates [49, 50, 51], while coarse-grained molecular simulations similarly represent proteins at the residue level. Wiggle provides a validated SAXS forward model operating directly on Cα traces, enabling scattering evaluation without requiring intermediate all-atom reconstruction.

Taken together, Carbonara provides a practical framework for generating diverse, physically viable seed structures. Its design reflects a deliberate response to the low information content of solution scattering. A conformational generator with sufficient degrees of freedom, such as a random chain model, can achieve good agreement with any experimental profile, rendering favourable χ^2^ largely uninformative about structural realism. Carbonara avoids this by preserving secondary structure elements throughout sampling, enforcing topologically derived constraints, and biasing sampling towards SAXS-consistent regions from the outset. Structural heterogeneity, therefore, emerges only when supported by both the experimental data and the structural constraints.

By rapidly producing SAXS-consistent conformations, Carbonara enables MD simulations to meaningfully explore solution relevant conformational spaces that would otherwise remain inaccessible on practical timescales. Understanding and predicting protein dynamics in solution remains a central challenge in structural biology and is increasingly important for interpreting the expanding set of structures produced by AI-based prediction methods [52]. In this context, Carbonara provides an experimentally grounded route from static structural models to solution-state ensembles.

## 5 Data and Software availability

Carbonara is available open-source at https://github.com/Prior-Lab-Durham-University/carbonara. Wiggle is available open-source at https://github.com/mckeownish/wiggle-SAXS. The data that support the findings of this study is available at http://doi.org/10.15128/r2dr26xx42n.

## 6 Acknowledgements

JM acknowledges the EPSRC and the SOFI2 CDT (grant EP/S023631/1) for financial support. AB acknowledges an EPSRC MoSMed CDT (EP/S022791/1) and a Diamond Light Source Ltd funded grant for financial support. CB acknowledges the EPSRC DTP and Diamond Light Source for financial support. HF acknowledges CRUK for financial support. This work used the ARCHER2 UK National Supercomputing Service (www.archer2.ac.uk) and the Durham HPC Hamilton and NCC. This project made use of time on HPC granted via the UK High-End Computing Consortium for Biomolecular Simulation, HECBioSim (http://hecbiosim.ac.uk), supported by EPSRC (grant no. EP/X035603/1).

## 7 Contributions

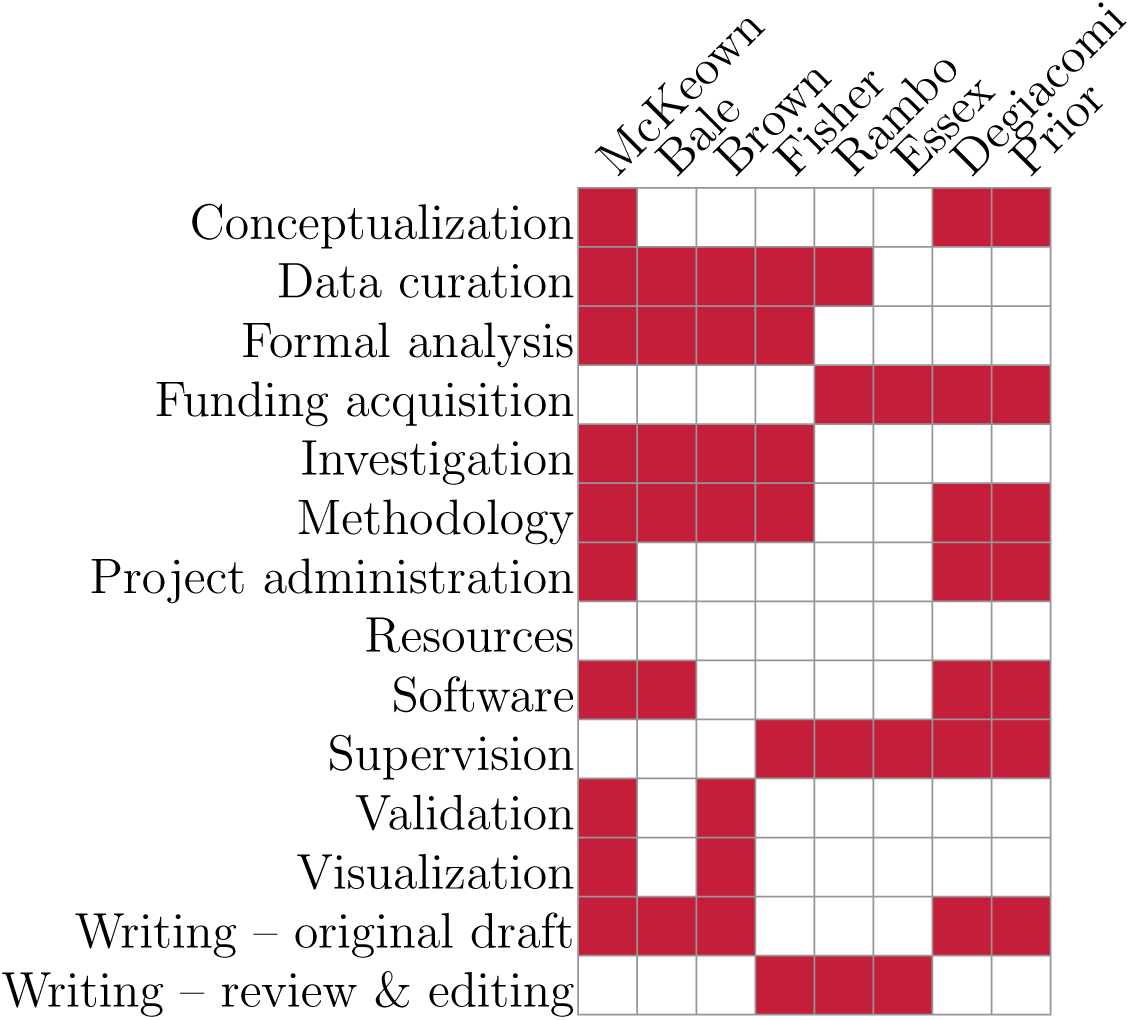

## 8 Methods

### 8.1 Wiggle implicit scattering model

#### 8.1.1 Coarse-grained representation and side-chain placement

Proteins are represented using a coarse-grained model defined on Cα coordinates, augmented with a side chain scattering centre for each non-glycine, non-alanine residue. The side chain centre of mass (COM) is placed along the local backbone normal, a unit vector derived from the curvature of the Cα curve at residue-specific distances d*_a_* determined empirically from heavy atom COMs across the training set, further details can be found in Supplementary Methods 1.2.2. This two centre representation, at Cα and approximate side chain COM, captures residue specific scattering without requiring side chain reconstructions. The accuracy and robustness of this geometric approximation were assessed by comparison to true sidechain centres of mass across the training dataset; detailed angular error statistics and validation analyses are reported in Supplementary Methods 1.2.

#### 8.1.2 The forward scattering model

SAXS intensities are computed using the Debye scattering formula [53]

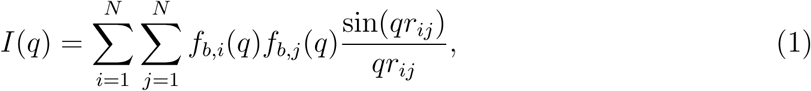

where f*_b,i_*, f*_b,j_* are the form factors for each coarse-grained bead type b and 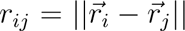 is the Euclidean distance between beads i and j.

The form factors of each of the 21 bead types were optimised such that the SAXS signal of our coarse-grained representation matches that of a reference all-atom dataset. To this end, we assembled a training dataset of 1570 protein structures selected to ensure Wiggle generalises across a broad range of folds, molecular weights, and conformational states (see Supplementary Methods 1.2.1). SAXS profiles for each of these reference structures were calculated using WAXSiS [31]. The form factors were optimised by minimising an objective function L_total_ that balances fit quality and with physically motivated smoothness constraints

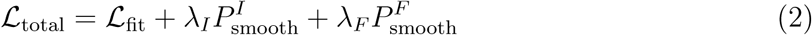

The fit term L_fit_, is defined as the average mean absolute percentage error (MAPE)

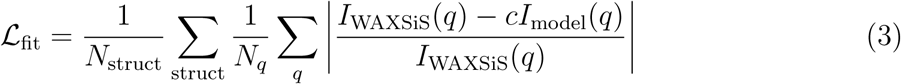

where I_WAXSiS_ is the scattering profile generated by WAXSiS, I_model_ the scattering generated by Wiggle as described above, and c is a scaling factor determined independently for each structure. MAPE was chosen rather than the more established χ^2^ metric because WAXSiS generates scattering profiles without experimental uncertainties.

The term 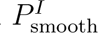 penalises oscillations in the calculated intensity curves

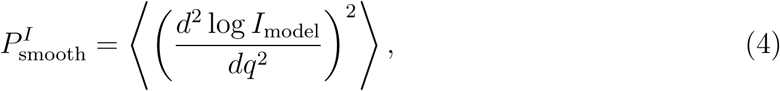

ensuring the form factors collectively produce physically realistic scattering profiles. The term 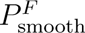 ensures the individual form factors F(q) are smooth:

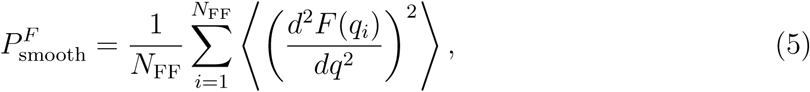

In both cases the averaging ⟨⟩ was taken over all 21 form factors and for the penalty 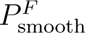, N*_ff_* is the number of points used to create an interpolatory function for the Form Factors (we use N*_ff_* = 20 on q ∈ [0, 0.2]). This combination of penalties helps prevent some of the unphysical oscillations in individual form factors, while allowing flexibility to capture residue-specific scattering and the impact of implicit hydration.

The hyperparameters λ*_I_* = 10*^−^*^2^ and λ*_F_* = 10*^−^*^1^ were selected to balance fit quality against smoothness, with form factor smoothness weighted more heavily. For further details, see Supplementary Methods 1.2.3.

### 8.2 Assessing model quality during a Carbonara fitting run

#### 8.2.1 SAXS intensity scaling and goodness-of-fit evaluation

For each candidate structure (or mixture of structures), Carbonara computes the predicted scattering intensity I_model_(q) using the Wiggle coarse-grained scattering model. Prior to evaluating the quality of fit, a single global intensity scale factor is determined independently of the optimisation objective.

Specifically, the logarithmic difference between the model and experimental intensities,

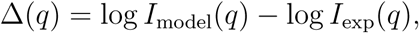

is computed and averaged over a restricted low-to-intermediate scattering range q ∈ [q_min_, q_min_+ a], with a = 0.1 Å^-1^by default. This mean logarithmic offset defines a multiplicative scale factor

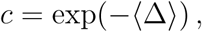

which is applied uniformly to the model intensity,

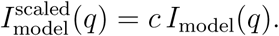

This procedure enforces agreement between the geometric mean intensities of the model and experimental curves over the scaling window, while explicitly avoiding least-squares reoptimisation of the scale factor for each trial structure. The scaling window width a = 0.1 Å^-1^ was selected to prioritise the large scale structure of the protein as discussed in the main text.

After scaling, the agreement between model and experiment is evaluated in linear intensity space over the user-selected fitting range q ∈ [q_min_, q_max_] using an error-weighted reduced chi-squared statistic,

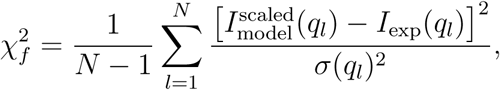

where *N* is the number of experimental data points within the fitting range and σ(q*_l_*) denotes the corresponding experimental uncertainty.

Rather than minimising 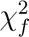 directly, Carbonara minimises the absolute deviation from unity,

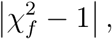

reflecting the expectation that a statistically consistent structural model should yield 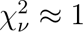. This choice discourages artificial overfitting of experimental noise while retaining sensitivity to systematic structural mismatch. The resulting score is combined with additional structural plausibility penalties to define the overall objective function guiding conformational exploration.

#### 8.2.2 Overlap prevention measure χ*_o_*

To prevent physically unreasonable conformations, a penalty χ*_o_* is applied when any pair of non-adjacent Cα atoms are separated by ≤ 4 Å. While a harmonic potential was used previously [38], here we adopt an exponential form,

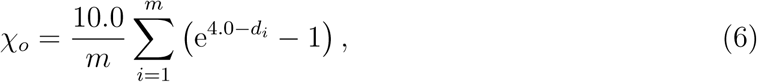

where d*_i_* are the m non-adjacent Cα–Cα distances below the threshold. This formulation produces a comparable penalty near 4 Å while enforcing a more strongly repulsive response at smaller separations.

#### 8.2.3 Custom distance restraints χ*_d_*

Carbonara supports optional distance restraints on selected pairs of Cα atoms to incorporate external structural information, such as disulfide bonds or predicted contacts. For n*_p_* restrained pairs with target distances 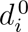, the penalty is defined as

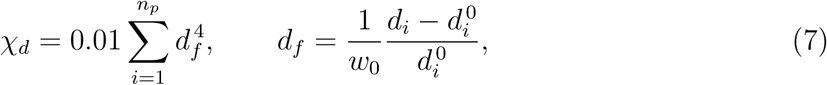

where d*_i_* is the instantaneous distance and w_0_ controls the tolerance of the restraint. With w_0_ = 1, d*_f_* corresponds to the fractional deviation from the ideal distance. The default value w_0_ = 0.5 sets χ*_d_* = 10*^−^*^4^ for deviations comparable to those tolerated by a high-quality scattering fit (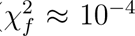), ensuring consistent weighting between geometric and scattering terms.

#### 8.2.4 Topological fold metric χ*_t_*

To prevent unrealistically unfolded conformations, Carbonara enforces a lower bound on backbone self-entanglement using the absolute crossing number (acn) of a smoothed Cα trace. Protein structures in the PDB exhibit a length-dependent minimum acn [39]; a brief summary of this measure is given in Supplementary methods 1.3. Violations of this bound are penalised using

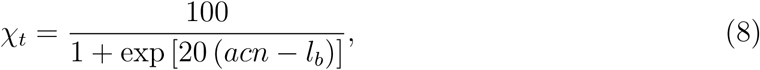

where l*_b_* is the empirical lower bound for a protein of the given length. Structures satisfying acn > l*_b_* incur negligible penalty, while overly extended conformations yield χ*_t_* ≈ 0.5, dominating the objective function and effectively excluding such geometries.

#### 8.2.5 Conformational optimisation

For a given fitting range q ∈ [q_min_, q_max_], Carbonara minimises the total objective

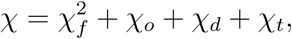

with χ*_d_* = 0 if no distance restraints are specified. Conformational exploration proceeds via a Monte Carlo scheme that iterates over each designated flexible region. At each step, a new backbone geometry is proposed (Supplementary Methods 1.1), and the corresponding χ value is evaluated. A proposed configuration is accepted only if it improves the total objective.

Because all penalty terms are included explicitly, improvements in steric or topological plausibility may occasionally coincide with small increases in the scattering term. The relative magnitudes of χ*_o_* and χ*_d_* are chosen such that such cases correspond only to minor, physically tolerable deviations. All accepted improvements are logged during execution and made available to the user in real time, enabling access to intermediate predictions without requiring completion of the full optimisation.

### 8.3 All-atom reconstruction of Carbonara predictions

All-atom protein structures were reconstructed from Carbonara-generated Cα traces using a customised MODELLER-based pipeline (Supplementary Methods, Section 1.4) [54]. For each predicted backbone, residue identities, chain identifiers, and residue numbering were recovered from the Cα trace and used to guide reconstruction of the full atomic model.

Secondary-structure information was incorporated through chain-aware restraints derived from DSSP [55] or STRIDE [56] assignments (or user-provided annotations). Contiguous helical and strand segments were restrained during model building, while linker regions were left unrestrained to preserve conformational flexibility and avoid artificial bias. No secondary structure restraint crossed chain boundaries in multimeric systems.

Model refinement was performed using MODELLER. For each input Cα trace, multiple trial models were generated and ranked using the DOPE statistical potential [57], with the top-scoring model selected. Final structures were completed to full heavy-atom representation and synchronised to preserve the original chain identifiers and residue numbering.

Analyses of the SMARCAL*^CD^* and ChiLob models (Supplementary Methods 1.6,1.7 and Supplementary Data 2.1 and 2.2) demonstrate that all-atom reconstruction introduces only minor backbone perturbations (Mean TM Scores > 0.96), confirming that the (κ, τ)-sampled conformations generated by Carbonara are stereochemically realistic.

### 8.4 SAXS evaluation of all-atom models

Theoretical SAXS profiles for all-atom structures were computed using FoXS [26] (via the pyFoXS interface) and CRYSOL (ATSAS version 3.3.0) [28, 33]. Where these exhibited systematic discrepancies, WAXSiS [31] was used to adjudicate, given its explicit solvent treatment of hydration. WAXSiS was not used for routine large scale evaluation due to its high computational cost. Unless stated otherwise, all methods were used in their standard or recommended operating modes, see [33] (CRYSOL), [26] (FoXS), and [31](WAXSiS).

### 8.5 Model generation

#### 8.5.1 SMARCAL1

We analysed a truncated structure of the Human SMARCAL1 protein, spanning residues 325 to 870, which contains both the HARP2 and ATPase domains [42]. We produced an atomistic model with AlphaFold3 [5], and identified its flexible regions using Carbonara’s automated procedure (see details in Supplementary Methods 1.5).

The SMARCAL1 data was previously reported in [42] and the figures from that paper are recreated in Fig. S24. The protein was purified by size exclusion chromatography with a Shodex KW-402 column in 20 mM HEPES (pH 7.5), 1% glycerol, 0.01% sodium azide, 2 mM TCEP, 200 mM NaCl and 2 mM MgCl_2_. During elution, 60 µL of peak fraction was taken for SAXS analysis as described in [42].

Secondary structure was identified with DSSP, from which flexible linkers were identified with our *automated* sheet preserving search. Fitting was performed for 20 parallel independent runs up to q = 0.2 with maximum steps set to 10,000. The final best fitting structures from each run were selected as the start point for MD simulations to generate dynamic ensembles.

#### 8.5.2 ChiLob 7/4 IgG2

The crystal structure of ChiLob7/4 IgG2 (PDB: 6TKD) and SAXS data (SASBDB: SAS-DLJ4) for its C239S variant were taken from [45]. Flexible linkers were assigned using the automated sheet-preservation option, which restricts flexibility primarily to the hinge region of the antibody (see Supplementary Methods 1.5).

Although the deposited structure contains four chains, the antibody forms a well-defined homodimer of two heavy-chain/light-chain units. To reflect this architecture during sampling, the two chains within each Fab unit were grouped and treated as rigid bodies. Carbonara was therefore restricted to sampling relative rotations and translations of the two dimer units about the hinge region.

To ensure the two F(ab) arms were held together, the Cα atoms of opposing disulfide bonding cysteine residues in the hinge region are constrained at 6 Å by Carbonara. The specific amino acid pairs subjected to disulfide bond restraints were 136−680, 149−205, 225−591, 231−686, 264 − 329, 375 − 435, 477 − 551, 604 − 660, 719 − 784, and 830 − 890. This distance constraint is particularly significant for disulfide bonds found within the hinge region, which confer the C239S variant its distinctive physiological properties [45].

The initial Carbonara seeding ensemble was generated using q*_max_* = 0.2 with a maximum of 1,000 sampling steps per run, executed as a batch of 20 parallel independent runs. The final structure from each run was used as a starting configuration for the subsequent MD simulations. The same parameters were used for the 5 batches of testing runs.

### 8.6 Comparison with alternative modelling methods

SAXS-A-FOLD modelling was performed using the public web server described in [34]. The AlphaFold model of SMARCAL1*^CD^* was uploaded together with the corresponding AlphaFold JSON file containing the predicted aligned error (PAE) scores. The server was run in the standard mode with default parameters. In this mode SAXS-A-FOLD automatically identifies flexible regions based on the PAE matrix and performs ensemble refinement to obtain models consistent with the experimental SAXS profile. The flexibility assignments suggested by the server were accepted without modification. All algorithmic details of the SAXS-A-FOLD optimisation procedure are described in [34].

Rigid-body modelling was performed using CORAL from the ATSAS package [33]. The AlphaFold model of SMARCAL1*^CD^* was manually divided into two rigid domains corresponding to residues 1-112 and 123-547. The intervening 10-residue segment (residues 113-122), corresponding to the linker connecting the HARP2 domain to the remainder of the ATPase core was treated as a flexible linker during the optimisation. CORAL therefore sampled the relative reorientation of these two structural units while representing the connecting region by dummy residues. To assess the influence of the SAXS fitting range, two sets of CORAL refinements were performed: one using the full experimental scattering range and one restricted to q ≤ 0.2 Å*^−^*^1^, consistent with the Carbonara sampling range used elsewhere in this study. For each fitting range, five independent CORAL runs were carried out using the default simulated-annealing parameters.

### 8.7 Molecular Dynamics simulations

MD simulations were performed using GROMACS (v2021.2 for ChiLob7/4 IgG2, v2023.4 for SMARCAL1*^CD^*). The Amber ff14SB force field [58] was employed for protein atoms and Joung-Cheatham parameters for ions [59]. Protein structures were prepared using Protein Preparation Wizard [60], capping chain termini with ACE and NMA residues and protonating at pH 7.0, followed by a restrained minimisation.

Systems were solvated in a TIP3P water box with a 2.5 nm buffer between the protein and box edges. Neutralisation was achieved by addition of Na^+^ and Cl*^−^* ions to reach a physiological concentration of 150 mM. Energy minimisation proceeded in two phases: 1000 steps of steepest descent followed by 1500 steps of conjugate gradient protocol. Equilibration was performed in two phases: first, a 50 ps NVT equilibration heating to 300 K, followed by a 100 ps NPT equilibration at 1 bar. Temperature was maintained using a Nose-Hoover thermostat [61] with a 2 ps*^−^*^1^ collision frequency, while pressure was controlled with a Parrinello-Rahman barostat [62] during equilibration using a 2.0 ps relaxation time. Independent production repeats in the NPT ensemble were performed using distinct random seeds for the thermostat. Simulations were all run with periodic boundary conditions with an 8 Å direct space cut-off for non-bonded interactions. Long-range electrostatics were handled by Particle Mesh Ewald summation [63], while long-range van der Waals interactions used an analytical dispersion correction. LINCS constraints [64] were applied to all bonds involving hydrogen, allowing a 2 fs timestep. From each production run, we extracted snapshots at 1 ns intervals.

For SMARCAL1, 3 x 2µs production runs were performed for the AlphaFold prediction structure and 20 x 300 ns for each of the Carbonara generated seed structures. For ChiLob7/4 IgG2, 3 x 2 µs production runs were performed for the Crystal structure and 20 x 3 x 100 ns for Carbonara generated seed structures.

### 8.8 Ensemble MD analysis and Bayesian reweighting

For SMARCAL1*^CD^*, we concatenated and aligned all the production runs (12,000 structures) on their Cα coordinates, residues 400-480, and then performed a Principal Component Analysis (PCA) on the Cα Cartesian coordinates. For ChiLob7/4 IgG2, terminal caps were removed and titratable residues renamed to match standard naming conventions. For each simulation snapshot (12,000 structures) χ^2^ values were calculated using FoXS [26] and radii of gyration (R*_g_*) calculated with Python library MDtraj [65].

Molecular dynamics (MD) simulations were analysed in conjunction with experimental SAXS data using an ensemble-based forward-modelling and reweighting framework. This protocol was applied identically across all systems studied to assess whether MD-generated ensembles adequately sample solution-state conformations consistent with experimental scattering data.

#### 8.8.1 Forward calculation of SAXS profiles

Theoretical SAXS intensities were computed independently for each MD simulation snapshot using both CRYSOL and FoXS. For SMARCAL1*^CD^*, systematic differences were observed between FoXS and CRYSOL (see Supplementary Data 2.1). Independent evaluation using the more physically detailed but computationally expensive WAXSiS model showed closer agreement with FoXS across the tested structures. On this basis, FoXS outputs were selected for subsequent analyses of both SMARCAL1*^CD^* and ChiLob7/4 IgG2 systems.

#### 8.8.2 Bayesian Maximum Entropy reweighting

MD ensembles were reweighted against experimental SAXS data using Bayesian Maximum Entropy [44]. For each ensemble, optimised weights were determined by minimising the negative log posterior, balancing the χ^2^ agreement with experiment against the Kullback-Leibler divergence from uniform weighting and controlled by a regularisation parameter θ. The optimal θ was selected by L-curve analysis [66] of N_eff_ –χ^2^ plots, identifying the elbow beyond which further reweighting provides diminishing improvements in fit quality at disproportionate cost to ensemble diversity [67, 68]. While a reduced χ^2^ = 1 is often adopted as a target, this criterion assumes perfectly calibrated experimental uncertainties and is only one approach (Gull-Daniell) within the general Bayesian ensemble refinement framework [67].

### 8.9 Visualisation

All plots were generated in Python (3.11) with the matplotlib (3.8.2) library. Molecular visualisations were produced with VMD 1.9.4 [69].

