## Supplemental Information for "Carbonara: a SAXS-guided seeding framework for exploring protein solution-state dynamics"

‖*ESRF - The European Synchrotron, 71 Avenue des Martyrs, 38043, Grenoble, France*

⊥*School of Informatics, University of Edinburgh, Edinburgh, UK*

### 1 Supplementary Methods

#### 1.1 The tertiary backbone model used by Carbonara

In the following we summarise the essential components of the tertiary backbone model introduced in[1] to interpret small angle scattering BioSAXS data.

##### Constraints on local backbone geometry: curvature and torsion

The C$\alpha$ backbone is modelled as a discrete set of three dimensional coordinates $\{\mathbf{x}_i\}$ expressed in a Cartesian coordinate system. These backbone curves are generated using Monte-Carlo sampling from a pair of distributions of two discrete geometric quantities which enforce realistic secondary geometry of this curve. The first quantity is the curvature $\kappa$, which describes the tightness of coiling of the curve (it is larger for an $\alpha$-helix than a $\beta$ strand) and is represented by the inverse of the sphere circumscribed on a section of four C$\alpha$ coordinates, as shown in fig S1(a) (one can see a less tightly coiled curve section fig S1(b) has a larger inscribed sphere). The second quantity is the torsion $\tau$ which represents the local chiral nature of the backbone. It is positive for right handed coiling and negative for left handed coiling. One can see in fig S1(c) it has a very similar definition to the Ramachandran torsion angles, except here it represents the angle $\theta_n$ of the two plane normals constructed from four C$\alpha$ coordinates $(\mathbf{x}_1, \mathbf{x}_2, \mathbf{x}_3, \mathbf{x}_4)$. An example from a real protein structure is shown in Fig S1(d). To formally define these quantities for the set $(\mathbf{x}_1, \mathbf{x}_2, \mathbf{x}_3, \mathbf{x}_4)$ we denote the midpoints between each pair as $\mathbf{c}_i = (\mathbf{x}_i + \mathbf{x}_{i+1})/2$ and calculate $(\kappa, \tau)$ for this quadruplet using the following formulae.

$$\kappa(\mathbf{x}_1, \mathbf{x}_2, \mathbf{x}_3, \mathbf{x}_4) = \frac{2|\sin(\theta_{123})|}{||\mathbf{c}_1 - \mathbf{c}_2||}, \tag{S1}$$

$$\tau(\mathbf{x}_1, \mathbf{x}_2, \mathbf{x}_3, \mathbf{x}_4) = \sigma\frac{2}{l}\sin(\theta_n/2), \tag{S2}$$

$$l = (||\mathbf{x}_2 - \mathbf{x}_1|| + ||\mathbf{x}_3 - \mathbf{x}_2|| + ||\mathbf{x}_4 - \mathbf{x}_3||)/3.$$

where $\theta_{123}$ is the angle between the vectors $\mathbf{c}_1 - \mathbf{c}_3$ and $\mathbf{c}_2 - \mathbf{c}_3$, $\theta_n$ is the angle made by the normal vectors $\mathbf{n}_1$ and $\mathbf{n}_2$ of the planes formed by the sets $(\mathbf{x}_1, \mathbf{x}_2, \mathbf{x}_3)$ and $(\mathbf{x}_2, \mathbf{x}_3, \mathbf{x}_4)$, and $\sigma$ is 1 for right handed rotation $-1$ for left handed rotation.

In[1] an algorithm was defined, the discrete backbone algorithm, which inverts these formulae (Eqs.S1 and S2) to generate a new C$\alpha$ coordinate $\{\mathbf{x}_i\}$ from a pair of $(\kappa_i, \tau_i)$ values and the prior three coordinates $(\mathbf{x}_{i-1}, \mathbf{x}_{i-2}, \mathbf{x}_{i-3})$. The C$\alpha$-C$\alpha$ distances are constrained to be 3.80 Å. We now describe how the values of $(\kappa, \tau)$ are constrained to be representative of local protein geometry.

**Imposing Ramachandran-like constraints on the local backbone geometry**

Values of $(\kappa, \tau)$ pairs for all subsections of over 10000 protein structures were calculated (these proteins were chosen to have less than 70% sequence similarity). These values were then separated if they had been classed as belonging to an $\alpha$-helix or a $\beta$ strand. The remaining values were classed as linker sections. Probability distributions of these data sets are shown in Figure S2. For the linker distribution there are three distinct isolated peaks which correspond to the peaks in the respective $\alpha$ helical and $\beta$-strand distributions. These peaks were shown in the supplement of[1] to correspond to the three dominant Ramachandran angles hence we refer to them as “Ramachandran-like”. One can find similar low resolution flexible C$\alpha$ backbone models in a number of published studies *e.g.*[2–7] To the best of our knowledge ours is the only model that explicitly couples to a solution scattering model for the purpose of interpreting BioSAXS data.

##### Refining the C$\alpha$ chain

The C$\alpha$ trace is split into segments by assigning subsets of atoms to three possible secondary structure types: helices, strands, or linkers (see Supplementary Methods 1.5). In any protein sequence, there are $n$ distinct secondary structure groupings in this assignment, $m$ of them linker sections. The geometry of an individual linker section is then altered as follows: say a given secondary grouping covers C$\alpha$ 's $l$ to $m$ (1-3 for the first one for example), with coordinates $\{\mathbf{x}_i\}_{i=l}^{m}$, then new values $\mathbf{x_i}$ are determined as described in the previous section by choosing values $\{(\kappa_i, \tau_i)\}_{i=l}^{m}$ for each coordinate from the linker distribution, and generating new coordinates $\{\mathbf{x}'_i\}_{i=l}^{m}$ which ensure the chain locally has these curvature and torsion values. As illustrated in Figure 1 of the main text this alters the geometry of that sub-section of the curve and also then applies a translation of the coordinates $\mathbf{x}_j$ $j > m$ (as the vector separating $\mathbf{x}_{m+1}$ and $\mathbf{x}_m$ remains fixed). Thus, the new configurations sampled are representative of plausible geometries and allow the side-chains to be reinserted later.

##### Specifying the flexibility of the molecule

This model requires that the secondary structure of the protein under study is defined. Carbonara can do this in several ways:

- Parsing PDB-defined assignment from the input structure
- Applying DSSP to the initial structure[8]
- Applying STRIDE to the initial structure[9]
- Accepting a user-defined assignment.

In this work we used DSSP. This yields a set of linker sections imparting flexibility to the model. While initially every linker is defined as flexible, the user can manually lock a subset of them. For example, in the case of ChiLob 7/4 IgG2, the linkers connecting constant and variable domains were not marked for variation, while those connecting the inter-F(ab) domains (hinge region) were.

This selection can potentially significantly restrict the search space. To ease this choice, in the Carbonara Python front end, we provide a routine which tests which sections of the structure can be varied without altering any $\beta$-sheet structures (by testing if any sheet bonds are broken). This provides (if requested) an initial suggestion for which linker sections to be rendered flexible and which to keep fixed.

#### 1.2 *Wiggle* Development and Validation

##### 1.2.1 Constructing the Wiggle Training Dataset

To achieve sufficient coverage of the protein sequence space while minimising redundancy, we embedded 35,000 protein sequences from the Protein Data Bank[10] using the ESM-2 language model (650M parameters),[11] a protein language model pre-trained on 65 million protein sequences. ESM-2 embeddings capture evolutionary relationships and structural properties, providing a biologically informed basis for clustering. We clustered these embeddings with k-means, and selected the sequences closest to each cluster. The final dataset contained 1570 protein structures.

To evaluate the diversity of our selected structures, we performed independent clustering analysis using MMseq2[12] at a 30% sequence identity threshold. This revealed 92.04% diversity with 1445 clusters identified from 1570 sequences. This minimal redundancy was confirmed by CD-HIT,[13] reporting a 90.68% diversity at a higher 40% identity threshold. These results demonstrate that our ESM-2-based selection strategy successfully captured a broad and unbiased coverage of the protein sequence space suitable for model training. The resulting data set not only featured proteins with a broad range of sequence lengths, but also with different levels of compaction and secondary structure content.

##### 1.2.2 Geometric Prediction of Sidechain Centres of Mass

Side-chain centres of mass are placed geometrically from the C$\alpha$ trace alone. Given a sequence of C$\alpha$ coordinates $\{\mathbf{r}_i\}_{i=1}^{N}$, backbone vectors are defined as

$$\mathbf{v}_i = \mathbf{r}_{i+1} - \mathbf{r}_i. \tag{S3}$$

For interior residues, a local geometric normal vector is computed from backbone curvature,

$$\mathbf{n}_i = -\frac{\mathbf{v}_{i+1} - \mathbf{v}_i}{\|\mathbf{v}_{i+1} - \mathbf{v}_i\|}, \tag{S4}$$

which points approximately outward from the local backbone trajectory. For terminal residues, where this quantity is ill-defined, directions are assigned as

$$\mathbf{n}_1 = -\frac{\mathbf{v}_1}{\|\mathbf{v}_1\|}, \qquad \mathbf{n}_N = \frac{\mathbf{v}_{N-1}}{\|\mathbf{v}_{N-1}\|}. \tag{S5}$$

The side-chain COM for a residue of type $a$ with C$\alpha$ position $\mathbf{r}_i$ is then placed at

$$\tilde{\mathbf{r}}_{\mathrm{COM},a} = \mathbf{r}_i + d_a\,\mathbf{n}_i, \tag{S6}$$

where $d_a$ is a residue-specific scalar distance between the C$\alpha$ atom and the side-chain COM (except Gly and Ala). The values of $d_a$ were determined empirically from the training data set by averaging the mass-weighted COMs of all non-backbone heavy atoms for each residue type:

$$\mathbf{r}_{\mathrm{COM}} = \frac{\sum_i m_i \mathbf{r}_i}{\sum_i m_i} \tag{S7}$$

Values of $d_a$ for each residue type are tabulated in Table S1.

We assessed the deviation between our predicted geometric normals and true COM directions across all residue types for all proteins in the training dataset, described in spherical coordinates (Figure S4). We found only a small systematic bias (mean $\Delta\theta = -0.022$, $\Delta\phi = -0.013$), with an approximately radially symmetric distribution of errors ($\sigma_{\Delta\theta} = 58.2°$, $\sigma_{\Delta\phi} = 27.8°$). The total angular error, $\omega_{err} = \sqrt{\Delta\theta^2 + \Delta\phi^2}$, has a mean of 52.8° and median 42.2°, and 90% of predictions fall within 110.1° of the true direction. This level of precision is acceptable for our coarse-grained framework, where exact atomic positions are less critical than capturing the overall spatial distribution of sidechain COM.

##### 1.2.3 Form Factor Optimisation

For each of the 1570 structures in the protein dataset, we generated reference SAXS profiles using the WAXSiS server,[14] which employs explicit MD simulations to model the hydration layer. The WAXSiS model avoids assumptions about hydration shell structure and provides high quality reference data for training our coarse-grained model. The resulting dataset was randomly partitioned into training (70%), test (15%) and validation (15%) sets, using fixed random seeds for reproducibility. The validation set monitored convergence model generalisation during training, while the test set remained held out until final evaluation.

Optimisation used the Adam algorithm[15] with an initial learning rate of $2 \times 10^{-1}$, using the loss function described in Main Text (see Methods). To avoid local minima we implemented a cosine annealing warm restart scheduler[16] with an initial cycle length of 100 epochs, doubling at every restart, with a minimum rate of $2 \times 10^{-3}$ (1% of the initial value). Training involved mini-batches of 4 randomly chosen structures per iteration. For each batch, we calculated the Debye intensity using the current form factor values along with the smoothness penalties. The form factors were then updated using back-propagation. They were constrained to non-negative values, as negative form factors are physically meaningless. Training was performed on an NVIDIA GPU (3080) using PyTorch. Early stopping terminated training if no test loss improvement occurred for 1,000 epochs.

##### 1.2.4 Performance against test and validation set

The optimisation of Wiggle form factors converged after approximately 1500 epochs, with a best test set performance of MAPE 3.21%. Final evaluation on the completely held out validation set gave a MAPE score of 3.30%. This minimal difference between test and validation suggests a very good model generalisation to unseen protein structures. Figure S5 illustrates the optimised form factors for all 21 scattering centre types. As expected for implicit hydration models,[1,17,18] the form factors exhibit non-monotonic behaviour as a function of $q$, reflecting the interplay between amino acid scattering, excluded volume effects, and surrounding solvent contributions. Figure S6 presents the distribution of relative errors across training, test, and validation splits, demonstrating consistent performance across all dataset partitions.

##### 1.2.5 Performance against experimental benchmark set

Table S2 summarises the agreement between experimental SAXS data and theoretical scattering profiles computed using Wiggle and established all-atom methods. Because the experimental profiles were generated by averaging a large number of high-precision measurements, the associated experimental uncertainties are exceptionally small. As a consequence, conventional error-weighted $\chi^2$ values are uniformly large across all methods, limiting their usefulness as an absolute measure of model quality.

To enable a more interpretable comparison, we therefore focus primarily on the mean absolute percentage error (MAPE), which quantifies the average relative deviation between predicted and experimental intensities and is independent of the experimental error model. MAPE provides a direct, scale-free measure of predictive accuracy and is particularly well suited to this consensus benchmark, where inflated $\chi^2$ values would otherwise obscure meaningful differences in performance.

Across the benchmark proteins, Wiggle achieves MAPE values that are comparable to those obtained with established all-atom scattering models, consistently falling within the same order of magnitude. For completeness, the corresponding $\chi^2$ values are also reported in Table S2 (in parentheses), allowing direct comparison with prior SAXS studies while recognising the limitations of $\chi^2$ in this specific low-noise regime.

The MAPE analysis reveals that Wiggle achieves average errors of 1-6% across the five proteins. While Wiggle does not consistently match the accuracy of all-atom methods with explicit hydration for smaller proteins, it performs within 1 percentage average error of the next ranked method. Notably, Wiggle outperforms CRYSOL for both urate oxidase (6.28% vs. 6.61%) and xylose isomerase (4.39% vs. 5.06%), the two multimers. This performance pattern is consistent with the expected physics, where hydration shell contributions, which Wiggle models implicitly, constitute a larger fraction of the total scattering signal for smaller proteins, while for larger proteins the protein-protein scattering dominates.

Visual inspection of the fits (Figure S25) reveals that Wiggle captures the overall profile shapes and characteristic features across all five proteins. The error-weighted residuals demonstrate that the elevated $\chi^2$ values primarily reflect small systematic deviations rather than fundamental failures in prediction. All methods, including Wiggle, show similar patterns of oscillatory residuals arising from the exceptionally small experimental uncertainties in the consensus profiles.

For the smaller proteins (xylanase, lysozyme), Wiggle shows a slight underestimate for larger q compared to explicit hydration methods, consistent with the expected limitations of implicit hydration modelling for small proteins. However, the overall profile shapes remain correct, indicating that the coarse-grained representation captures the dominant scattering contributions. For larger proteins (urate oxidase, xylose isomerase), Wiggle's residual patterns become comparable to all-atom methods.

For the intended application of large-scale conformational sampling requiring millions of SAXS calculations, Wiggle's performance represents a favourable trade-off between accuracy and computational efficiency. Wiggle achieves MAPE of 2.0-6.3% across the benchmark proteins, performing within a similar range to CRYSOL (1.0-6.6%, uniform hydration shell model), WAXSiS (1.2-3.9%, explicit MD-simulated hydration), and Pepsi-SAXS (0.5-5.4%, explicit grid solvent model), while FoXS (0.5-3.6%, fitted hydration parameters) shows the best overall performance. Remarkably, Wiggle achieves this accuracy despite fundamental differences from all other methods; it requires only C$\alpha$ coordinates as input and employs no explicit hydration model or fitted implicit hydration parameters. The comparable performance suggests that the implicit absorption of hydration effects into residue-specific form factors that were optimised against high-quality WAXSiS reference data, successfully captures some of the physics governing protein scattering in the $q \leq 0.20$ Å$^{-1}$ regime. Importantly, none of the proteins in this benchmark set were included in Wiggle's training dataset, providing an independent test of generalization on gold-standard experimental data.

###### 1.2.6 Synthetic mixture recovery

To assess Wiggle's ability to discriminate between distinct conformational states, twenty pairs of crystal structures representing identical sequences in different conformations were selected from the PDB databank (Table S3). For each pair, WAXSiS scattering profiles were generated for both conformations and combined to produce synthetic mixture profiles

$$I_{\text{mix}}(q) = p \cdot I_1(q) + (1 - p) \cdot I_2(q), \tag{S8}$$

where $p$ represents the fractional population of state 1, evaluated at $p = 0.2$, 0.5, and 0.7. Profiles were aligned at $q = 0$ before averaging. Wiggle was then used to generate scattering profiles for both structures and the optimal mixing ratio $\hat{p}$ was determined by minimising the mean absolute error between the synthetic WAXSiS mixture and the weighted Wiggle prediction:

$$\hat{p} = \underset{p \in [0,1]}{\text{argmin}} \frac{1}{N} \sum_q \left| I_{\text{WAXSiS,mix}}(q) - c \left[ p \cdot I_{\text{model},1}(q) + (1 - p) \cdot I_{\text{model},2}(q) \right] \right|, \tag{S9}$$

where $c$ is a scaling constant determined independently for each structure. Optimisation used the Adam algorithm.[15] Across all 60 tests, Wiggle recovered the correct ratios with mean absolute errors of 0.079–0.086 and negligible systematic bias (Fig. S7).

###### 1.2.7 Experimental mixture recovery

The same procedure was applied to recover population mixtures from experimental SAXS data for calmodulin and bovine serum albumin, using crystal structures of the known conformational states as described in the main text. For calmodulin, the open (PDB: 1CLL, $R_g$ = 21.9 Å) and closed (PDB: 1CFC, $R_g$ = 18.7 Å) states were used. For bovine serum albumin, the three MultiFoXS-optimised conformers ($R_g$ = 26.4, 29.8, and 30.4 Å) identified in[19] were used, with the mixture fraction extended to three components accordingly.

##### 1.3 A description of the topological fold metric $\chi_t$

In order to produce realistically entangled structures, a writhe-based constraint was introduced to the model. The absolute crossing number (ACN) is a positive definite measure of global self-entanglement of a curve. Intuitively, if we project a curve onto a plane, there will be points of self intersection. We can count these so-called crossings. The number of crossings will necessarily be dependent on the angle of projection chosen, so the ACN can be thought of as an average of the sum of crossings over all directions. The ACN of a smooth curve $\boldsymbol{\gamma}(t)$ is given by the Gauss linking integral[20]

$$ACN = \frac{1}{4\pi}\int_{\gamma}\int_{\gamma} \boldsymbol{T}(s) \times \boldsymbol{T}(t) \cdot \frac{|\boldsymbol{\gamma}(s) - \boldsymbol{\gamma}(t)|}{\|\boldsymbol{\gamma}(s) - \boldsymbol{\gamma}(t)\|^3}\,\mathrm{d}s\,\mathrm{d}t, \tag{S10}$$

where $T(t)$ is the tangent vector to the curve $\boldsymbol{\gamma}(t)$. Since we are considering the discrete protein backbone curve, we use the discrete analogue of the ACN given by

$$ACN(\mathcal{C}) = 2\sum_{i=2}^{n-1}\sum_{j<i}\frac{|\Omega_{ij}|}{4\pi}, \tag{S11}$$

where $\Omega_{ij}$ represents the contribution to Eq. S10 from the crossing of edges connecting $\boldsymbol{x}_i$ to $\boldsymbol{x}_{i+1}$ and $\boldsymbol{x}_j$ to $\boldsymbol{x}_{j+1}$. There are numerous equivalent methods for computing $\Omega$, and we will follow Method 1a given in.[21] For this, we denote by $\boldsymbol{r}_{i,j}$ the edge between points $\boldsymbol{x}_i$ and $\boldsymbol{x}_j$. We then define the unit normal vectors:

$$\boldsymbol{n}_1 = \frac{\boldsymbol{r}_{i,j} \times \boldsymbol{r}_{i,j+1}}{\|\boldsymbol{r}_{i,j} \times \boldsymbol{r}_{i,j+1}\|}, \tag{S12}$$

$$\boldsymbol{n}_2 = \frac{\boldsymbol{r}_{i,j+1} \times \boldsymbol{r}_{i+1,j+1}}{\|\boldsymbol{r}_{i,j+1} \times \boldsymbol{r}_{i+1,j+1}\|}, \tag{S13}$$

$$\boldsymbol{n}_3 = \frac{\boldsymbol{r}_{i+1,j+1} \times \boldsymbol{r}_{i+1,j}}{\|\boldsymbol{r}_{i+1,j+1} \times \boldsymbol{r}_{i+1,j}\|}, \tag{S14}$$

$$\boldsymbol{n}_4 = \frac{\boldsymbol{r}_{i+1,j} \times \boldsymbol{r}_{i,j}}{\|\boldsymbol{r}_{i+1,j} \times \boldsymbol{r}_{i,j}\|} \tag{S15}$$

We consider the sum of the angles between these vectors

$$\Omega^* = \sin^{-1}(\boldsymbol{n}_1 \cdot \boldsymbol{n}_2) + \sin^{-1}(\boldsymbol{n}_2 \cdot \boldsymbol{n}_3) + \sin^{-1}(\boldsymbol{n}_3 \cdot \boldsymbol{n}_4) + \sin^{-1}(\boldsymbol{n}_4 \cdot \boldsymbol{n}_1). \tag{S16}$$

Then the evaluation of the Gauss integral from the crossing of $\boldsymbol{r}_{i,i+1}$ and $\boldsymbol{r}_{j,j+1}$ is given by

$$\frac{\Omega_{ij}}{4\pi} = \frac{\Omega^*}{4\pi} \operatorname{sgn}\left((\boldsymbol{r}_{j,j+1} \times \boldsymbol{r}_{i,i+1})\boldsymbol{r}_{i,j}\right) \tag{S17}$$

To compute the ACN of a protein, we define the backbone curve which is the discrete 3-dimensional curve connecting the central C$\alpha$ atom of each amino acid residue. To reduce the impact of the helical nature of secondary structures on the ACN calculation, we must smooth the backbone in an appropriate manner. To do this, we apply the SKMT algorithm[22] to reduce it to a minimal representation that preserves any essential non-local entanglement of the backbone curve. Adapting the KMT algorithm[23,24] to act solely within secondary structure elements (SSEs), we effectively replace each SSE with a straight edge with the number of edges for the linkers determined by checking if it reduces mutual entanglement.

As a result, the length of the SKMT smoothed backbone curve is proportional to the number of distinct SSEs.

#### 1.4 C$\alpha$-to-All-Atom Model Reconstruction with Secondary Structure Restraints

This section provides implementation-level details of the all-atom reconstruction pipeline summarised in the main Methods.

All-atom protein models were reconstructed from C$\alpha$-only backbone traces using a custom pipeline built around the `MODELLER` `automodel` framework.[25] The procedure first parses the input PDB file and extracts the residue identity, chain identifier, and residue index for each C$\alpha$ atom using a regular-expression filter applied to `ATOM` records. These data are used both to assemble the corresponding one-letter amino-acid sequence (with "/" denoting chain breaks) and to record the original residue numbering for later reinstatement. A temporary PIR alignment file is then generated, in which the C$\alpha$-only structure serves as the template (`model_ca`) for reconstruction of the full all-atom model.

Model building is performed using a subclassed `automodel` object that implements two types of custom restraints. First, original chain identifiers are restored via `rename_segments` within the `special_patches` method. Second, residue-level secondary-structure restraints are applied using a user-provided annotation list (`ss_list`). Contiguous segments annotated as helix (H) or strand (S) are automatically detected and converted into `MODELLER` $\alpha$-helix or $\beta$-strand restraints, respectively, using the `secondary_structure` module. Regions annotated as coil receive no additional restraints. Structural refinement is carried out using either `MODELLER`'s `refine.slow` or `refine.fast` molecular-dynamics protocols, and $N$ models are generated for each input C$\alpha$ trace. Both methods are tested in Supplementary Data 2.1 for their effect on the scattering model.

All generated models are evaluated using the DOPE statistical potential. The top-scoring model is processed with `complete_pdb` to ensure full completeness. To ensure that the output preserves the coordinate conventions of the input structure, the chain identifiers and residue indices from the original C$\alpha$ trace are re-mapped onto the reconstructed full-atom model. This is achieved by iterating synchronously through the final `ATOM` records and the original C$\alpha$ entries, and rewriting the relevant columns of each line to match the input numbering. This procedure guarantees that the final PDB file contains a fully rebuilt heavy-atom structure while maintaining the original chain topology and residue numbering.

All temporary files generated during the procedure (including intermediate alignment files, template PDBs, individual trial models, and log files) are removed automatically upon completion, leaving only the final full-atom model written to the user-specified output file.

##### 1.4.1 Adjustments for Multimeric Protein Reconstruction

For multimeric targets, the C$\alpha$-to-all-atom reconstruction procedure was extended to correctly handle multiple chains, chain-specific secondary-structure assignments, and optional inter- or intra-chain disulfide bond definitions. The multimeric version of the pipeline processes the input PDB in the same manner as for monomers, but retains chain identifiers for every C$\alpha$ atom and inserts "/" separators in the PIR sequence to denote chain boundaries. This ensures that `MODELLER` treats each polypeptide chain as an independent modelling segment during alignment and refinement.

A customised subclass of `automodel` was used to impose chain-level structural constraints. Within the `special_patches` method, the chain identifiers extracted from the input C$\alpha$ trace were reapplied to the model using `rename_segments`, enabling consistent chain naming throughout refinement and in the final output PDB. If a list of disulfide bonds was provided by the user, each pair of residues specified in the form `resnum:chain` was located within the model, and the corresponding cysteine residues were linked using the `DISU` patch. Missing or invalid residue identifiers were reported but did not interrupt model generation.

Secondary-structure restraints were adapted to the multimeric case by ensuring that restraints do not cross chain boundaries. The residue-wise secondary-structure annotation list (`ss_list`) was scanned in parallel with the chain identifier list extracted from the input structure. Contiguous stretches of residues with identical secondary-structure labels were converted into helical or strand restraints only if they belonged to the same polypeptide chain; transitions between chains automatically terminated the current secondary-structure segment. This procedure yields chain-specific helix and strand definitions that respect domain boundaries and prevent artificial inter-chain restraints.

Model refinement was performed using the `refine.fast` protocol to reduce computational cost while maintaining high-quality stereochemistry. Multiple models ($N$ iterations) were generated and evaluated using the DOPE statistical potential. The top-scoring model was completed using `complete_pdb`, written to disk, and subsequently renumbered so that each chain starts from residue 1 while preserving chain ordering. All temporary files produced during the procedure were removed upon completion.

#### 1.5 Secondary structure assignment

##### SMARCAL1$^{CD}$

The secondary structure assignment for the catalytic domain of Human SMARCAL1$^{CD}$, obtained from DSSP, was:

```
------SSSSSSS----SSSSSS---HHHHHHHH-----SSS----SSSSSHHHHHHHHHHH-----SSSS---
HHHHHH---HHH------------------HHHHHH--HHHHHHHHHHHH---SSSS-------HHHHHHHHHH
H-HHH--SSSSS----HHHHHHHHHHH-----HHHSSS------------SSSSSHHHH------------SSS
S--HHHH-----HHHHHHHHHHHH--SSSSS--------HHHHHHHHHHH-------HHHHHHHH---SSS--SS
S------HHHHHHHHHHH------HHHH------SSSSSSS--HHH--HHHHHHHHHHHHHHH-----HHHHHHHH
HHHHHHHHHHHHHHHHHHHHHHHH-----SSSSS--HHHHHHHHHHHHH----SSSS-------HHHHHHHHHHH-
----SSSSS--------------SSSS------HHHHHHHH-----------SSSSSSS-----HHHHHHHHHHHH
HHHHH---------HHHH--
```

with, S representing a $\beta$ strand, H an $\alpha$-helix and - a linker section. The linker sections highlighted in red were those permitted to be changed during the fitting procedure.

##### ChiLob 7/4 IgG2

Using the same notation as for SMARCAL1$^{CD}$, the four initial chains of the ChiLob 7/4 IgG2 molecule were set into two chains representing the homodimer units of the molecule (this restricts the applied rigid body motions to the two dimer units with respect to each other). The secondary structure assignment, using DSSP, was as follows:

***Chain 1***

```
--SSSS---SSS-----SSSSSSSS---HHH-SSSSSSS-----SSSSSSS------SSS-HHH---SSSSSS--
--SSSSSS----HHH-SSSSSSSSSS-----SSSSSS---SSSSS--------SSSSS----------SSSSSSS
SSSSS----SSSSHHH-----SSS----------SSSSSSSSSS---HHH--SSSSSSS----SSSSSSS-----
-------------------------SSSSS----SSSSSSS-------SSSSSS-----SSSSSS----------
--SSSSSS--SSSSSS----HHH--SSSSSS-----------SSSSSS------SSSSS---HHHH---SSSSSS
SSSSSS----SSSSSS--------SSSSS---------SSSSSSSSSSHHHHHH--SSSSSSS-------SSSSS
S----
```

***Chain 2***:

```
--SSSS---SSS-----SSSSSSSS---HHH--SSSSSS-----SSSSSSS------SSS-HHH---SSSSSS-
---SSSSSS----HHH-SSSSSSSSSS-----SSSSSS--SSSSS--------SSSSS----------SSSSSSS
SSSSS----SSSSHHH-----SSS----------SSSSSSSSSS---HHH---SSSSSS----SSSSSS------
-------------------------SSSSS----SSSSSSS-------SSSSSS-----SSSSSS----------
--SSSSSS--SSSSSS----HHH--SSSSSS-----------SSSSSS------SSSSS---HHHH---SSSSSS
SSSSSS----SSSSSS--------SSSSS---------SSSSSSSSSSHHHHHH--SSSSSSS-------SSSSS
S----
```

The highlighted sections form parts of the hinge region.

#### 1.6 Assessing secondary structure preservation in the Carbonara prediction pipeline

Secondary structure conservation between the reference model and predicted structures was assessed using DSSP assignments. For each structure, secondary structure was computed using the DSSP algorithm as implemented in `MDTraj`. The standard eight-state DSSP classification was reduced to a three-state representation to facilitate comparison. Helical states (H, G, I) were grouped as **H**, $\beta$-strand states (E, B) as **S**, and all remaining states were grouped as $-$ (coil/loop).

For each chain in the reference and model structures, the amino-acid sequence and the corresponding three-state secondary structure string were extracted. Chains between reference and model structures were matched using global sequence alignment. Only chain pairs exceeding a minimum sequence identity threshold of 0.3 were retained for comparison. The aligned sequences were then used to map secondary structure assignments residue-by-residue while excluding positions corresponding to alignment gaps.

Secondary structure preservation was quantified using three complementary metrics. First, a residue-level agreement score, $Q(\mathrm{H}, \mathrm{S}, -)$, was calculated as the fraction of aligned residues with identical three-state secondary structure assignments between the reference and model structures,

$$Q(\mathrm{H}, \mathrm{S}, -) = \frac{1}{N} \sum_{i=1}^{N} \delta(s_i^{\mathrm{ref}}, s_i^{\mathrm{model}}), \quad \text{(S18)}$$

where $s_i^{\mathrm{ref}}$ and $s_i^{\mathrm{model}}$ denote the secondary structure assignments at residue $i$ in the reference and model respectively, $N$ is the number of aligned residues, and $\delta$ is the Kronecker delta. Second, helix segment preservation was assessed by identifying contiguous helical segments in the reference structure and computing their overlap with corresponding segments in the model structure using an intersection-over-union measure,

$$\mathrm{IoU}(A, B) = \frac{|A \cap B|}{|A \cup B|}, \quad \text{(S19)}$$

where $A$ and $B$ denote the residue ranges of reference and model segments. For each reference helix segment, the maximum overlap with any model helix segment was calculated and the average across all reference helices was reported. Third, an analogous segment-overlap metric was computed for $\beta$-strand segments. For ensembles of predicted structures, these metrics were computed for each model relative to the reference structure. Summary statistics (mean, standard deviation, median, and interquartile range) were then calculated across the ensemble to quantify the overall level of secondary structure conservation.

#### 1.7 Comparison of per-residue structural deviations between backmapping methods

To assess whether the two all-atom backmapping procedures introduced systematic structural differences, we compared the local deviations between corresponding models generated by each method. For each run, the structures produced by Method A and Method B were aligned and the root-mean-square deviation (RMSD) was calculated independently for every residue using all heavy atoms. This yielded a per-residue deviation profile describing the local structural difference between the two backmapped models for that run.

Let $N$ denote the number of runs and $R$ the number of residues considered. For each run $i$, a deviation vector

$$\mathbf{d}_i = (d_{i1}, d_{i2}, \ldots, d_{iR}) \tag{S20}$$

was constructed, where $d_{ir}$ is the RMSD for residue $r$ between the two methods in run $i$. Collecting these vectors forms a deviation matrix

$$\mathbf{M} \in \mathbb{R}^{N \times R}, \tag{S21}$$

where rows correspond to runs and columns correspond to residues.

To determine whether particular residues systematically exhibited larger deviations between the two backmapping methods, correlations were computed between the deviation profiles of all runs. Specifically, the Pearson correlation coefficient was calculated between each pair of rows of $\mathbf{M}$,

$$C_{ij} = \mathrm{corr}(\mathbf{d}_i, \mathbf{d}_j), \tag{S22}$$

yielding an $N \times N$ correlation matrix $\mathbf{C}$. If the two methods consistently differed in specific regions of the protein, the corresponding residues would produce similar deviation patterns across runs, resulting in strong off-diagonal correlations in $\mathbf{C}$.

To further investigate common patterns in the deviation matrix, principal component analysis (PCA) was applied to $\mathbf{M}$. Prior to analysis, missing entries were replaced with zeros to maintain a fixed dimensionality across runs. PCA produced orthogonal components that describe the dominant modes of variation in the per-residue deviation patterns. The explained variance ratio of each principal component was examined to determine whether the deviations were dominated by a small number of common structural modes or distributed across many independent contributions.

Together, the correlation analysis and PCA provide complementary views of whether structural differences between the two backmapping methods arise from systematic residue-specific effects or from run-to-run variability associated with the underlying conformational ensemble.

### 2 Supplementary Data

#### 2.1 Testing the Carbonara fitting pipeline

In what follows all calculations were performed on a Linux workstation running Fedora 42. The machine was equipped with an Intel Xeon Silver 4108 CPU (16 physical cores, 32 threads, base 1.8GHz) and 30GB RAM.

##### 2.1.1 Test on human SMARCAL1$^{CD}$

To characterise the robustness, practical behaviour, and computational efficiency of Carbonara on a large, flexible system, we performed extensive testing on human SMARCAL1$^{CD}$. We carried out 100 independent refinement runs (five batches of 20 parallel processes), each with a maximum of 10,000 optimisation steps using the default fitting range $q \leq 0.2$ Å$^{-1}$. For each logged Carbonara C$\alpha$ prediction, we generated all-atom models using both the fast and slow backmapping protocols (Section 1.4) and evaluated SAXS agreement using FoXS and CRYSOL.

**All-atom fit quality and agreement between SAXS forward models.** Across the ensemble, Carbonara produced a large improvement in SAXS agreement relative to the original AlphaFold structure. Over the full experimental range $q \in [0, 0.25]$, mean $\chi^2$ values were 1.578 (fast) and 1.533 (slow) using FoXS, compared with 15.670 for the AlphaFold model; the corresponding CRYSOL means were 2.209 (fast) and 2.147 (slow) compared with 12.350 for AlphaFold. When restricting evaluation to the Carbonara fitting range $q \in [0, 0.2]$, FoXS means improved slightly (1.423 fast, 1.387 slow) while CRYSOL values increased modestly (2.237 fast, 2.193 slow), consistent with a systematic offset between the two packages. Despite this offset, FoXS and CRYSOL broadly agreed in their ranking of structures (Pearson/Spearman correlations 0.712/0.722 over $q \in [0, 0.25]$ and 0.775/0.740 over $q \in [0, 0.2]$; full distributions in Fig. S8 and summary statistics in Table S4).

**Fast versus slow backmapping.** Differences between fast and slow backmapping were small relative to run-to-run variability. For FoXS, the mean fast–slow $\chi^2$ difference was $0.045 \pm 0.386$ over $q \in [0, 0.25]$ and $0.036 \pm 0.392$ over $q \in [0, 0.2]$. For CRYSOL, the corresponding differences were $0.062 \pm 0.649$ (full range) and $0.043 \pm 0.741$ ($q \in [0, 0.2]$). These results indicate that, at the resolution captured by SAXS scoring, fast and slow all-atom reconstruction yield effectively equivalent assessments, motivating the use of fast backmapping for high-throughput screening (with slow refinement reserved for shortlisted models).

**Comparison to explicit-solvent WAXSiS.** FoXS and CRYSOL use different approximations for excluded volume and hydration, which can lead to systematic differences in $\chi^2$ values for flexible or surface-heterogeneous proteins. To adjudicate between the two rapid evaluators on SMARCAL1$^{CD}$, we compared both against WAXSiS, which estimates scattering from explicit-solvent molecular dynamics snapshots and therefore avoids an implicit hydration-shell parametrisation.[14] We first selected the 10 structures exhibiting the largest FoXS–CRYSOL discrepancies (full-range evaluation) and re-scored them with WAXSiS. For these outliers, WAXSiS was substantially closer to FoXS than to CRYSOL: the mean difference (FoXS $\chi^2$ – WAXSiS $\chi^2$) was $-0.1165$, while the mean difference (CRYSOL $\chi^2$ – WAXSiS $\chi^2$) was 1.8665 (full values in Table S5). Because discrepancy-selected structures are not representative by construction, we repeated the comparison on an additional set of 20 predictions generated using the same Carbonara settings and fast backmapping. Treating each method's $\chi^2$ values as a vector over these 20 models, the RMS distance FoXS–WAXSiS was 0.160, compared with 0.277 for CRYSOL–WAXSiS and 0.356 for FoXS–CRYSOL (Supplementary Table S6). Taken together, these results indicate that for SMARCAL1$^{CD}$, FoXS is typically closer to explicit-solvent WAXSiS than CRYSOL, with the difference most pronounced in the largest-disagreement cases.

**The last logged structure is not necessarily the best all-atom fit.** Each 10,000-step run has a typical wall time of $\sim$ 9 h, but Carbonara logs a new prediction whenever the internal objective improves, so models become available continuously throughout sampling. Time-resolved analysis shows that all-atom fit quality (as assessed by FoXS and CRYSOL) often improves rapidly at early times and then fluctuates non-monotonically thereafter. Across the 100 runs, the median best-so-far $\chi^2$ dropped below 2 within $\sim$ 10–20 min for both FoXS and CRYSOL ( Fig. S12). Continued sampling increases the number of acceptable structures recovered, but later logged models are not guaranteed to yield better all-atom fits, reflecting differences between Carbonara's coarse-grained objective and all-atom forward models, as well as the influence of structural plausibility penalties. Consistent with this, selecting (post hoc) the prediction with $\chi^2$ closest to unity rather than the final logged prediction improves mean fit quality: over $q \in [0, 0.25]$, the mean closest-to-one scores are 1.173 (FoXS) and 1.329 (CRYSOL), compared with the final-prediction means reported above; over $q \in [0, 0.2]$, the corresponding values are 1.145 (FoXS) and 1.334 (CRYSOL) ( Table S7).

**Structural variation introduced by all-atom backmapping.** Because the backmapping protocols include an energy-minimisation step, we quantified whether reconstruction introduces systematic perturbations relative to the Carbonara C$\alpha$ traces. Comparing each C$\alpha$ prediction to the C$\alpha$ trace extracted from its reconstructed all-atom model yields small deviations for both protocols: for fast backmapping, RMSD $1.216 \pm 0.370$ Å and TM-score $0.980 \pm 0.011$; for slow backmapping, RMSD $1.062 \pm 0.617$ Å and TM-score $0.982 \pm 0.013$ ( Table S8). Secondary structure is also strongly preserved relative to the AlphaFold reference, this was assessed using the metrics described in supplementary Methods 1.6. Using a three-state DSSP reduction (H/S/-), the agreement metric $Q(\mathrm{H}, \mathrm{S}, --)$ is $0.903 \pm 0.012$ (fast) and $0.903 \pm 0.010$ (slow), with helix segment overlap $0.762 \pm 0.021$ (fast) and $0.761 \pm 0.023$ (slow), and strand segment overlap $0.711 \pm 0.059$ (fast) and $0.733 \pm 0.053$ (slow) ( Table S9). These values indicate that backmapping preserves global fold and secondary-structure topology, with differences limited to modest local adjustments.

Systematic residue-level differences between the two backmapping procedures were assessed using the per-residue RMSD correlation and principal component analyses described in the Supplementary Methods 1.7. The resulting correlation matrix showed no consistent off-diagonal structure ( Fig. S11)(a) and PCA revealed that variance was distributed across multiple components ( Fig. S11)(b), indicating that deviations between methods were small and largely configuration-dependent rather than associated with specific structural regions.

**Structural diversity of the Carbonara ensemble.** To quantify the conformational diversity produced by Carbonara on SMARCAL1$^{CD}$, we compared (i) each prediction to the AlphaFold starting structure and (ii) predictions to one another using RMSD and TM-score ( Fig. S16). Relative to AlphaFold, Carbonara predictions show a significant range of AlphaFold–prediction RMSD values ($\approx 5 - 30$ Å) and TM-scores ($\approx 0.1 - 0.75$), indicating a mixture of minor and major domain-level rearrangements away from the starting model. Similarly the prediction–prediction comparisons span a broad range: RMSD values extend from 2 to 30 Å with an approximately even spread across $\sim$ 5–25 Å, while TM-scores range from 0.1 to 0.95 with modal peak around 0.28. Thus, Carbonara does not simply generate a narrow cluster of nearby conformers, but produces a structurally diverse ensemble that remains broadly physically plausible at the fold level.

**Expected time for high-quality predictions** The previous analyses demonstrate that Carbonara consistently produces structures that agree well with the experimental SAXS data. A practical question for users is therefore how long a refinement must run before high-quality predictions begin to appear.

In practice Carbonara is normally executed in batches of **20 parallel refinement runs**, which together generate a pool of candidate conformations. From the user's perspective it is therefore useful to consider how many acceptable predictions accumulate over time during such a batch run. Figures S13 and S14 show the cumulative number of predictions with

$\chi^2 < 2$ and $\chi^2 < 1.5$ respectively, obtained across the five batches of twenty runs used in this study. These plots demonstrate that multiple SAXS-consistent models typically appear rapidly once sampling begins. Orders of 10's of structures are obtained within 20–40 minutes of runtime, and the number of qualifying predictions continues to increase steadily as the runs progress. Consequently, even relatively short Carbonara runs generally produce a useful ensemble of SAXS-compatible starting conformations.

Taken together, these results indicate that Carbonara routinely identifies high-quality SAXS-consistent conformations on timescales of minutes rather than hours, and that running multiple refinements in parallel rapidly yields a diverse set of candidate structures suitable for downstream analysis.

**Alternative fitting ranges ($q^c_{\max}$).** To examine how the chosen Carbonara fitting window controls downstream all-atom fit quality across different spatial scales, we repeated the SMARCAL1$^{CD}$ experiments using reduced fitting ranges $q^c_{\max} = 0.10$ and 0.15 (each with 100 runs), and re-evaluated the resulting final structures using FoXS and CRYSOL across multiple windows ($q \in [0, 0.10]$, $[0, 0.15]$, $[0, 0.20]$, and the full experimental range). Two robust patterns emerge (full distributions in Table S10). First, models generated with the default $q^c_{\max} = 0.20$ already achieve strong performance when evaluated on restricted low-$q$ windows: for example, when scored on $q \in [0, 0.10]$ and $q \in [0, 0.15]$, the central tendency of $\chi^2$ is comparable to that obtained when Carbonara is run directly on those reduced ranges, indicating that low-$q$ agreement is not degraded by fitting a broader window. Second, there is a clear deterioration in all-atom $\chi^2$ when models are evaluated on $q$-ranges extending substantially beyond those enforced during the Carbonara stage, consistent with the fact that higher-$q$ regions increasingly reflect finer structural detail and hydration modelling choices not explicitly targeted by the coarse-grained search. Finally, the FoXS–CRYSOL discrepancy is smallest on the reduced low-$q$ windows and becomes more pronounced as the evaluation range extends to higher $q$, consistent with the increasing influence of hydration and excluded-volume approximations at intermediate-to-high $q$.

**Practical guidance for SMARCAL1$^{CD}$.** Overall, these analyses show that Carbonara reliably produces SAXS-consistent SMARCAL1$^{CD}$ conformers across independent runs, with rapid early discovery of acceptable fits and continued accumulation of diverse candidates throughout sampling. Fast and slow backmapping yield near-identical conclusions at the level resolved by SAXS scoring, motivating fast reconstruction for high-throughput screening. For SMARCAL1$^{CD}$ in particular, explicit-solvent WAXSiS comparisons indicate that FoXS is typically closer than CRYSOL, especially for the largest-disagreement outliers. Together, these results support the use of Carbonara as an ensemble-generating front-end that rapidly identifies SAXS-consistent conformational basins, providing suitable starting pools for downstream all-atom refinement and ensemble-reweighting workflows.

##### 2.1.2 Comparison of Carbonara to alternative methods for Human SMARCAL1$^{CD}$

Structural prediction sets were generated from the AlphaFold3 starting model using CORAL and SAXS-A-Fold, and compared with a Carbonara ensemble comprising 20 independent batch runs. The number of models analysed reflects the standard output of each workflow rather than an attempt to equalise sampling depth: Carbonara batch processing naturally produces multiple structures, CORAL refinements were performed as ten independent optimisations, and SAXS-A-Fold returns a small parsimonious ensemble by design. Comparisons should therefore be interpreted as reflecting the characteristic structural behaviour of each method under routine usage conditions.

RMSD distributions relative to the starting AlphaFold structure (Fig. S10(a)) show that Carbonara models span approximately 7-26 Å, with values extending across this range. In contrast, the three SAXS-A-Fold predictions exhibit substantially smaller deviations (0, 9.04, and 9.13 Å), consistent with limited modification of the initial fold. CORAL predictions display larger structural deviations of approximately 27-33 Åbut a much more limited range of variations than the Carbonara predictions.

A similar pattern is observed in TM-scores (Fig. S10(b)). Carbonara structures span approximately 0.15–0.65, indicating substantial rearrangement while retaining partial similarity to the starting topology. SAXS-A-Fold predictions yield TM-scores of 1.0, 0.71, and 0.67, reflecting preservation of the AlphaFold-predicted domain organisation. In contrast, CORAL predictions exhibit very low TM-scores ( 0–0.07), consistent with extensive alteration of the initial domain arrangement.

These differences arise from the distinct modelling formalisms employed. SAXS-A-Fold primarily applies constrained domain motions and linker sampling around the initial fold. Carbonara reconstructs backbone geometry under SAXS restraints while preserving continuous chain connectivity. CORAL performs rigid-body optimisation of structured domains connected by flexible dummy-residue linkers, permitting substantial domain repositioning during SAXS fitting.

As illustrated in Fig. S18, the largest CORAL deviations correspond to displacement of the HARP domain from the structural core via extended linker conformations, whereas Carbonara models remain comparatively compact due to backbone continuity constraints.

To quantify intra-method variability, pairwise RMSD and TM-score comparisons were computed between predictions generated by each modelling approach. SAXS-A-Fold structures exhibit limited dispersion, with pairwise RMSD values of 4–10 Å and TM-scores of 0.65–0.83. The two refined models distinct from the original AlphaFold structure show the smallest mutual RMSD, indicating convergence toward a consistent alternative conformation.

Carbonara predictions display broader variability, with pairwise RMSD values spanning approximately 2–39 Å and TM-scores ranging from 0.1 to 0.9, consistent with extensive conformational sampling.

In contrast, CORAL predictions segregate into two TM-score clusters (0.1-0.4 and 0.75-0.98), indicating two internally coherent but mutually divergent structural solution sets. Repeated optimisations under a fixed scattering range produced limited additional dispersion, suggesting that stochastic variability alone does not account for this bifurcation. Instead, the two clusters correspond to optimisation performed over different fitting ranges ($q \in [0, 0.2]$ and $q \in [0, 0.27]$), demonstrating sensitivity of the resulting structures to the selected scattering interval.

#### 2.2 Test on ChiLob7/4

ChiLob7/4 provides a complementary test case to SMARCAL1$^{CD}$, representing a larger, oligomeric assembly with limited intrinsic flexibility. In this system, most secondary and tertiary structural features are already well specified by the starting homology model, and large-scale conformational rearrangements are neither expected nor supported by prior experimental evidence. The following analyses assess whether Carbonara preserves this intrinsic rigidity while still improving agreement with SAXS data, and whether the method refrains from introducing artificial structural diversity when the data do not require it.

**Carbonara reliably improves the fit across all data ranges.** Carbonara was applied using five batches of twenty independent refinement runs for each fitting range $q_c^{\max} = 0.20$, 0.15, and 0.10, with each run limited to 1,000 optimisation steps. All-atom models were generated using the fast backmapping protocol and evaluated using both FoXS and CRYSOL across multiple $q$ ranges. Mean best-fit $\chi^2$ values across the 100 runs are summarised in Table S11. Across all fitting ranges, Carbonara consistently improves agreement with experimental SAXS data relative to the starting homology model. This improvement becomes progressively more pronounced as the fitting range is restricted to lower $q$. For $q \in [0, 0.1]$ the fit improves from $\chi^2 = 9.57$ down to a mean value of 1.859 for FoXs, and from $\chi^2 = 9.306$ down to 1.559 for CRYSOL. On the full range $q \in [0, 0.37]$ the $\chi^2$ value reduces from 3.626 down to 1.731 (FoXS) and 3.739 down to 1.734 (CRYSOL). The improvements for the intermediate ranges are between these values. This indicates that the dominant discrepancy in the homology model lies in its global domain arrangement rather than in local structural detail. Focusing optimisation to low-$q$ therefore enables Carbonara to focus on correcting large-scale rigid-body misalignments without perturbing well-defined secondary structure.

**FoXS and CRYSOL assessments are broadly aligned** In contrast to SMARCAL1$^{CD}$, FoXS and CRYSOL assessments for ChiLob7/4 remain closely aligned across the full and intermediate $q$ ranges. As indicated in Figure S9, Pearson and Spearman correlation coefficients exceed 0.9 for $q \in [0, 0.37]$ and $q \in [0, 0.2]$, indicating strong agreement in how the two methods rank structures. This consistency reflects the relative rigidity and compactness of the assembly, for which hydration and excluded-volume effects play a less dominant role in shaping the scattering profile. At smaller fitting ranges ($q \leq 0.15$ Å$^{-1}$), CRYSOL yields systematically lower $\chi^2$ values than FoXS, a trend also visible in the mean statistics. This divergence is consistent with methodological differences between the two packages. CRYSOL's use of a uniform Gaussian hydration shell and spherical-harmonics representation effectively smooths low-$q$ scattering contributions, which can partially absorb modest residual discrepancies in global size or shape. In contrast, FoXS employs a surface-based Debye formulation with solvent-accessibility-weighted hydration terms, leading to increased sensitivity to subtle global mismatches. The strong correlations observed across all ranges indicate that this difference reflects a systematic offset rather than inconsistent structural ranking.

**Structural variation introduced by all-atom backmapping.** Across all predictions the comparison of the original C$\alpha$ trace CARBONARA backbone prediction and the subsequent all atom predictions are summarised in table S12 for each of the three fitting ranges. As with the same tests for Human SMARCAL1$^{CD}$, the results show the structures are essentially the same thus the backmapping procedure is not fundamentally affecting the nature of the prediction. Assessment of secondary-structure preservation relative to the original homology model was performed for each fitting range. As detailed in Table S13, the overall agreement metric $Q(\mathrm{H}, \mathrm{S}, --)$ remained stable at $\approx 0.86$ across all three ranges, with strand-segment overlap of $\approx$ 0.73–0.75. The helical agreement values are lower ($\approx$ 0.35–0.39); however, $\alpha$-helical content represents only $\sim 6\%$ of the sequence and is predominantly composed of very short segments (three residues). Such short helices are inherently sensitive to small local structural variations and therefore contribute disproportionately to reductions in the helix-specific agreement score. Overall, these results indicate that the backmapping procedure preserves the global fold and secondary-structure topology, with deviations largely limited to modest local adjustments.

**Relative absence of structural diversity of the Carbonara ensemble.** Pairwise comparisons between refined structures are summarised via Histograms in Figure S17. By comparison to the Human SMARCAL1$^{CD}$ results reveal limited structural variation between the predictions (majority values between $0.5-10$Å and $0.4-0.98$ TM score). The same is true of the variations from the original homology model (majority values between $3-10$Å and $0.65-0.9$ TM score), Figure S17. This demonstrates Carbonara's ensemble-first strategy is permissive rather than prescriptive: substantial diversity emerges when supported by the data, but refinement remains conservative for structurally constrained systems.

**Expected time for high-quality predictions** Fig. S15 shows the cumulative number of predictions with $\chi^2 < 2$ obtained across the five batches of 20 runs used in this study, assessed on the domains $q \in [0, 0.37]$ and $q \in [0, 0.1]$. All results presented are for $q_{max} = 0.2$.

As in the case of the SMARCAL1$^{CD}$ molecule, these plots demonstrate that multiple SAXS-consistent models typically appear rapidly once sampling begins. Orders of 10's of structures are obtained within 20 minutes of runtime, and the number of qualifying predictions continues to increase steadily as the runs progress. Once again we see even relatively short Carbonara runs generally produce a useful ensemble of SAXS-compatible starting conformations. The fact that the number of structures is broadly similar for both domains indicates that the structures are improving due to large-scale rearrangements of the quaternary structure of this Tetramer.

**Performing mixture refinements substantially increases the range of structural variability of the predictions** Refinements were performed with mixtures of two states (for which the relative mixture percentage was varied for values between (10%,90%) to (90%,10%) for states 1 and 2 respectively. The range of Rg values obtained from these prediction runs compared to single fit run sets with $q^c_{max} = 0.2$ and $q^c_{max} = 0.1$ are shown in Fig S19. The mixture state predictions have a much wider $R_g$ variation $34 - 42$Å compared to the single state predictions $38 - 41$Å for $q^c_{max} = 0.2$ and $38.5 - 40.5$Å for $q^c_{max} = 0.1$. As discussed in the main text, the mixture state prediction $R_g$ range is more commensurate with those of the Carbonara seeded MD simulations (Fig. 6(c) of the main text).

### 3 Supplementary Tables

Table S1: Characteristic distances from C$\alpha$ to side chain centre of mass for each residue type, determined empirically from our protein test set. Glycine and alanine are represented by single scattering centres at their C$\alpha$ positions, hence have no side chain distances.

| **Res** | **Dist (Å)** | **Res** | **Dist (Å)** | **Res** | **Dist (Å)** | **Res** | **Dist (Å)** |
|---|---|---|---|---|---|---|---|
| ALA | — | GLN | 3.18 | LEU | 2.62 | SER | 1.96 |
| ARG | 4.24 | GLU | 3.23 | LYS | 3.60 | THR | 1.95 |
| ASN | 2.53 | GLY | — | MET | 3.19 | TRP | 3.88 |
| ASP | 2.55 | HIS | 3.18 | PHE | 3.40 | TYR | 3.88 |
| CYS | 2.38 | ILE | 2.31 | PRO | 1.88 | VAL | 1.95 |

Table S2: Comparison of SAXS prediction methods on experimental benchmark data. Values show mean absolute percentage error (MAPE) with $\chi^2$ in parentheses, calculated over $q \in [0, 0.20]$ Å$^{-1}$ for predictions from crystal structures against consensus experimental profiles.[26] MAPE $= \frac{1}{N}\sum_i \left|\frac{I_{\text{data}}(q_i)-I_{\text{model}}(q_i)}{I_{\text{data}}(q_i)}\right| \times 100$ provides a scale-independent measure of relative accuracy. $\chi^2 = \frac{1}{N}\sum_i \frac{[I_{\text{data}}(q_i)-I_{\text{model}}(q_i)]^2}{\sigma^2(q_i)}$ where $\sigma(q_i)$ is the experimental uncertainty. The very small experimental errors in consensus profiles lead to large absolute $\chi^2$ values despite good agreement.

| **Method** | **RNaseA** | **Urate Oxidase** | **Xylose Isomerase** | **Xylanase** | **Lysozyme** |
|---|---|---|---|---|---|
| CRYSOL | 1.25 (154.9) | 6.61 (152.5) | 5.06 (160.4) | 0.97 (26.54) | 0.93 (25.42) |
| Pepsi-SAXS | 0.69 (44.5) | 5.39 (99.0) | 2.48 (40.5) | 0.50 (9.7) | 0.49 (12.9) |
| FoXS | 1.43 (207.3) | 3.55 (55.5) | 2.07 (25.3) | 0.52 (5.51) | 0.45 (12.9) |
| WAXSiS | 2.23 (707.9) | 3.48 (47.4) | 3.89 (50.4) | 1.19 (38.7) | 1.62 (92.3) |
| **Wiggle** | **2.95 (677.4)** | **6.28 (166.8)** | **4.39 (59.0)** | **1.97 (68.7)** | **2.63 (200.6)** |

Values represent MAPE (%) with $\chi^2$ in parentheses. Lower MAPE indicates better agreement with experimental data; $\chi^2$ is dominated by experimental uncertainties. PDB entries: 7rsa (RNaseA), 3l8w (urate oxidase, with C-terminal residues modeled), 1mnz (xylose isomerase, with N-terminal Met added), 2dfc (xylanase), 2vb1 (lysozyme).

Table S3: Structural characterisation of protein conformational state pairs used for mixture recovery validation. All pairs represent identical (or near-identical) sequences in different conformational states. TM-scores and RMSD values quantify structural similarity and magnitude of conformational change after optimal superposition of all C$\alpha$ atoms. Synthetic mixture profiles were generated from WAXSiS calculations at ratios of 20:80, 50:50, and 70:30 (state 1:state 2, notated 0.20, 0.50 and 0.70, respectively). Values show the predicted population of state 1 from our coarse-grained model. See Figure S7

| **Protein (Abbrev.)** | **State 1 ($R_g$)** | **State 2 ($R_g$)** | **$\Delta R_g$** | **TM-score** | **RMSD (Å)** | **0.20** | **0.50** | **0.7** |
|---|---|---|---|---|---|---|---|---|
| Adenylate Kinase (AK) | 4AKE (19.6) | 1AKE (16.6) | 3.0 | 0.68 | 7.1 | 0.20 | 0.49 | 0.68 |
| Calmodulin (CaM) | 1CLL (21.9) | 1CFC (18.7) | 3.2 | 0.37 | 14.0 | 0.25 | 0.51 | 0.69 |
| Citrate Synthase (CS) | 2CTS (22.3) | 1CTS (22.8) | 0.5 | 0.93 | 2.4 | 0.35 | 0.63 | 0.81 |
| Vibrio cholerae LapD GGDEF-EAL (GGDEF) | 6PWK (23.6) | 6PWJ (22.2) | 1.4 | 0.63 | 16.5 | 0.21 | 0.47 | 0.63 |
| Glutamine Binding Protein (GlnBP) | 1WDN (17.6) | 1GGG (19.1) | 1.5 | 0.66 | 5.3 | 0.18 | 0.47 | 0.65 |
| Hexokinase (HK) | 1DGK (39.6) | 1HKC (41.0) | 1.4 | 0.86 | 4.4 | 0.22 | 0.50 | 0.69 |
| Heat Shock Protein 70 (Hsp70) | 4JNE (29.1) | 2KHO (37.6) | 8.5 | 0.57 | 33.7 | 0.22 | 0.51 | 0.70 |
| Lactoferrin (LF) | 1CB6 (28.4) | 1LFG (29.5) | 1.1 | 0.83 | 6.4 | 0.23 | 0.50 | 0.67 |
| Maltose Binding Protein (MBP) | 1ANF (20.7) | 1OMP (21.6) | 0.9 | 0.80 | 3.8 | 0.07 | 0.44 | 0.69 |
| Periplasmic binding protein (PBP) | 6MLP (17.8) | 6ML0 (19.9) | 2.1 | 0.69 | 5.1 | 0.42 | 0.72 | 0.92 |
| Phosphoglycerate Kinase (PGK) | 16PK (23.5) | 3PGK (24.1) | 0.6 | 0.41 | 12.8 | 0.22 | 0.73 | 1.00 |
| Polymerase Beta (Pol-$\beta$) | 1BPX (22.8) | 1BPY (21.9) | 0.9 | 0.89 | 2.8 | 0.34 | 0.68 | 0.91 |
| PRPP synthetase monomer (PRPP) | 1ECC (23.1) | 1ECJ (23.9) | 0.8 | 0.95 | 3.3 | 0.23 | 0.52 | 0.71 |
| Ribose Binding Protein (RBP) | 1URP (20.3) | 2DRI (19.4) | 0.9 | 0.73 | 4.1 | 0.00 | 0.25 | 0.49 |
| D-allose binding protein (ABP) | 1RPJ (19.3) | 1GUD (21.0) | 1.7 | 0.72 | 4.45 | 0.00 | 0.29 | 0.51 |
| T4 phage beta-glucosyltransferase (BGT) | 1JG6 (20.9) | 1JEJ (21.9) | 1.0 | 0.93 | 2.1 | 0.01 | 0.46 | 0.74 |
| EPSP Synthase (EPSPS) | 1RF4 (21.1) | 1RF5 (22.6) | 1.5 | 0.83 | 3.69 | 0.22 | 0.52 | 0.72 |
| Guanylate Kinase (GK) | 1EX6 (17.1) | 1EX7 (16.3) | 0.8 | 0.82 | 3.64 | 0.21 | 0.52 | 0.72 |
| GluR2 Ligand Binding Core (GluR2) | 1FTM (18.2) | 1FTO (19.1) | 0.9 | 0.89 | 2.26 | 0.27 | 0.62 | 0.85 |
| Insulin Hexamer (I-6-6) | 1TRZ-a4 (21.2) | 1TRZ-a3 (18.2) | 3.0 | – | – | 0.25 | 0.55 | 0.75 |

$R_g$ values shown in parentheses (Å). $\Delta R_g = |R_{g,1}$ - $R_{g,2}|$. TM-score and RMSD calculated using USalign with residue-based alignment.[27]

Table S4: mean $\chi^2$ values for FoXS and CRYSOL, comparing fast and slow backmapping modes and the original AlphaFold structure.

| q range | Method | Fast mode mean | slow mode mean | AlphaFold (original) |
|---|---|---|---|---|
| [0,0.25] | FoXS | 1.578 | 1.533 | 15.670 |
| | CRYSOL | 2.209 | 2.147 | 12.350 |
| [0,0.2] | FoXS | 1.423 | 1.387 | 15.060 |
| | CRYSOL | 2.237 | 2.193 | 14.640 |

Table S5: Fit to the full experimental range $q \in [0, 0.25]$: comparison of FoXS, CRYSOL, and WAXSiS for the 10 largest FoXS–CRYSOL discrepancy cases.

| Fit Number | Modeller mode | FoXS $\chi^2$ | CRYSOL $\chi^2$ | WAXSiS $\chi^2$ |
|---|---|---|---|---|
| 58 | fast | 2.446 | 5.015 | 2.221 |
| 80 | slow | 2.523 | 4.834 | 2.091 |
| 72 | slow | 2.292 | 4.788 | 2.675 |
| 70 | fast | 3.022 | 5.173 | 3.429 |
| 27 | slow | 1.697 | 3.717 | 1.694 |
| 69 | slow | 1.624 | 3.499 | 2.356 |
| 95 | slow | 1.809 | 3.515 | 1.919 |
| 55 | fast | 1.493 | 3.161 | 1.245 |
| 66 | fast | 1.831 | 3.411 | 1.653 |
| 11 | slow | 1.182 | 2.636 | 1.801 |

Table S6: Fit to the full experimental range $q \in [0, 0.25]$: comparison of FoXS, CRYSOL, and WAXSiS for 20 additional (representative) predictions (fast backmapping).

| Fit Number | FoXS $\chi^2$ | CRYSOL $\chi^2$ | WAXSiS $\chi^2$ |
|---|---|---|---|
| 1 | 1.180 | 2.744 | 1.52873 |
| 2 | 0.960 | 2.407 | 2.22085 |
| 3 | 1.513 | 2.724 | 2.27778 |
| 4 | 1.521 | 2.877 | 1.75465 |
| 5 | 1.679 | 2.235 | 1.49804 |
| 6 | 2.198 | 3.543 | 2.35855 |
| 7 | 1.291 | 3.539 | 2.22018 |
| 8 | 3.275 | 5.500 | 2.28408 |
| 9 | 1.566 | 2.374 | 2.11994 |
| 10 | 1.589 | 2.092 | 1.11772 |
| 11 | 2.026 | 2.869 | 2.28302 |
| 12 | 0.947 | 1.432 | 1.03187 |
| 13 | 1.725 | 2.824 | 2.44496 |
| 14 | 2.216 | 2.686 | 2.35047 |
| 15 | 2.891 | 3.943 | 2.68160 |
| 16 | 1.235 | 1.709 | 3.08930 |
| 17 | 1.400 | 4.040 | 1.35225 |
| 18 | 1.882 | 3.615 | 2.74110 |
| 19 | 1.180 | 2.308 | 1.99601 |
| 20 | 1.987 | 2.597 | 2.02477 |

Table S7: Mean values of the best fitting structure for the SMARCAL1$^{CD}$ structure (the structure whose $\chi^2$ value is closest to 1) from all 100 test runs.

| Method | $q \in [0, 0.25]$ | $q \in [0, 0.2]$ | AlphaFold (original) |
|---|---|---|---|
| FoXS | 1.173 | 1.145 | 15.670 / 15.060 |
| CRYSOL | 1.329 | 1.334 | 12.350 / 14.640 |

Table S8: RMSD and TM scores comparing the C$\alpha$ backbone of the Carbonara prediction and all atom structures backmapped using both the fast and slow modeller refinement protocols.

| Refinement Mode | RMSD | TM Score |
|---|---|---|
| Fast | $1.216 \pm 0.370$ | $0.980 \pm 0.011$ |
| Slow | $1.062 \pm 0.617$ | $0.982 \pm 0.013$ |

Table S9: Summary of the secondary structure preservation analysis for the comparisons of the refinement generated Human SMARCAL1$^{CD}$ structures and the original AlphaFold prediction.

| Refinement Mode | Metric | Mean ± SD | Median | IQR |
|---|---|---|---|---|
| fast | $Q(\mathrm{H}, \mathrm{S}, -)$ | 0.903 ± 0.012 | 0.903 | 0.897–0.912 |
| | Helix segment overlap | 0.762 ± 0.021 | 0.761 | 0.751–0.773 |
| | Strand segment overlap | 0.711 ± 0.059 | 0.711 | 0.678–0.759 |
| slow | $Q(\mathrm{H}, \mathrm{S}, -)$ | 0.903 ± 0.010 | 0.903 | 0.897–0.910 |
| | Helix segment overlap | 0.761 ± 0.023 | 0.759 | 0.751–0.774 |
| | Strand segment overlap | 0.733 ± 0.053 | 0.742 | 0.699–0.777 |

Table S10: (a) FoXS and CRYSOL mean (Standard deviation) qualities for Human SMARCAL1$^{CD}$ predictions obtained from Carbonara fits over varying $q$ ranges.

| Method | $q_{max}$ | $q \in [0, 0.1]$ | $q \in [0, 0.15]$ | $q \in [0, 0.2]$ | full range |
|---|---|---|---|---|---|
| | 0.10 | 0.997 (0.034) | 1.612 (1.327) | 3.070 (2.798) | 3.234 (2.545) |
| FoXS | 0.15 | 1.007 (0.084) | 0.999 (0.098) | 1.983 (1.068) | 2.078 (1.009) |
| | 0.20 | 1.111 (0.317) | 1.087 (0.289) | 1.145 (0.322) | 1.173 (0.335) |
| | 0.10 | 0.996 (0.034) | 1.612 (1.327) | 2.431 (1.407) | 4.142 (2.843) |
| CRYSOL | 0.15 | 0.988 (0.079) | 1.043 (0.156) | 2.431 (1.407) | 2.379 (1.240) |
| | 0.20 | 1.097 (0.310) | 1.121 (0.386) | 1.333 (0.454) | 1.330 (0.456) |

Table S11: fit quality for the Carbonara ChiLob7/4 predictions assessed using both the FoXS and CRYSOL models on various $q$ ranges. The first four lines represent the values of the initial homology model, the second four the lowest value obtained from the 100 Carbonara predictions.

| Model | q range | FoXS | CRYSOL |
|---|---|---|---|
| initial | $q \in [0, 0.37]$ | 3.626 | 3.739 |
| | $q \in [0, 0.2]$ | 5.489 | 5.563 |
| | $q \in [0, 0.15]$ | 6.792 | 6.949 |
| | $q \in [0, 0.1]$ | 9.567 | 9.306 |
| Carbonara | $q \in [0, 0.37]$ | 1.731 | 1.734 |
| | $q \in [0, 0.2]$ | 1.873 | 1.818 |
| | $q \in [0, 0.15]$ | 1.857 | 1.615 |
| | $q \in [0, 0.1]$ | 1.838 | 1.546 |

Table S12: Comparison of RMSD and TM scores comparing the C$\alpha$ backbone of the Carbonara ChiLob7/4 predictions and all atom structures backmapped using both the fast modeller refinement protocol.

| $q_c^{max}$ | Mean RMSD | SD RMSD | Mean TM Score | SD TM score |
|---|---|---|---|---|
| 0.20 | 1.927 | 0.808 | 0.973 | 0.025 |
| 0.15 | 1.988 | 0.813 | 0.971 | 0.027 |
| 0.10 | 2.117 | 1.011 | 0.965 | 0.039 |

Table S13: A summary of the secondary structure preservation analysis for the comparisons of the Carbonara derived ChiLob7/4 predictions obtained with a fitting ranges including data from $q \in [0, q_c^{max}]$.

| $q_c^{max}$ | Metric | Mean ± SD | Median | IQR |
|---|---|---|---|---|
| 0.2 | $Q(\mathrm{H}, \mathrm{S}, -)$ | 0.860 ± 0.014 | 0.860 | 0.851–0.870 |
| | Helix segment overlap | 0.391 ± 0.066 | 0.391 | 0.351–0.431 |
| | Strand segment overlap | 0.726 ± 0.033 | 0.724 | 0.704–0.746 |
| 0.15 | $Q(\mathrm{H}, \mathrm{S}, -)$ | 0.862 ± 0.013 | 0.860 | 0.854–0.872 |
| | Helix segment overlap | 0.405 ± 0.083 | 0.399 | 0.354–0.458 |
| | Strand segment overlap | 0.729 ± 0.027 | 0.728 | 0.713–0.747 |
| 0.1 | $Q(\mathrm{H}, \mathrm{S}, -)$ | 0.861 ± 0.014 | 0.862 | 0.853–0.873 |
| | Helix segment overlap | 0.394 ± 0.081 | 0.389 | 0.342–0.452 |
| | Strand segment overlap | 0.729 ± 0.031 | 0.731 | 0.707–0.748 |

### 4 Supplementary Figures

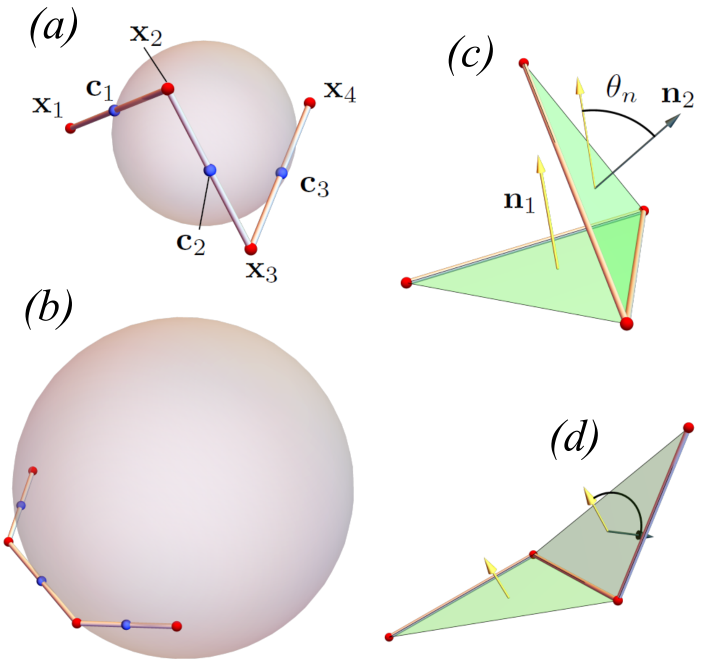


Figure S1: **Illustrations of the geometrical interpretations of the curvature $\kappa$ and torsion $\tau$ used to constrain the C$\alpha$ backbone model**. (a) four C$\alpha$ coordinates $\mathbf{x}_i$ and their midpoints $\mathbf{c}_i$. There is a circumscribed sphere touching these midpoints whose inverse radius defines the curvature. (b) demonstration that a less tightly coiled section of curve than in (a) sees a larger sphere and hence smaller curvature. (c) a section of a set of 4 coordinates and the two planes they define. The normal vectors to these planes $\mathbf{n}_1$ and $\mathbf{n}_2$ are shown. The angle $\theta_n$ they make is used to define the torsion and is a measure of the non planarity of the set. (d) demonstration that $\theta_n$ can be large.

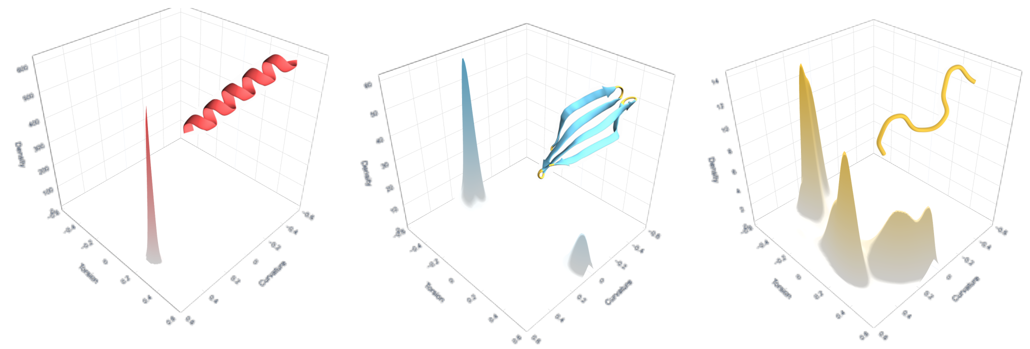


Figure S2: **Empirical probability densities for the curvature and torsion of each secondary structure type**. These densities are used to generate new secondary structure in the internal Carbonara C$\alpha$ trace model. Left to right the $\alpha$-helix, $\beta$-strand, and linker distributions.

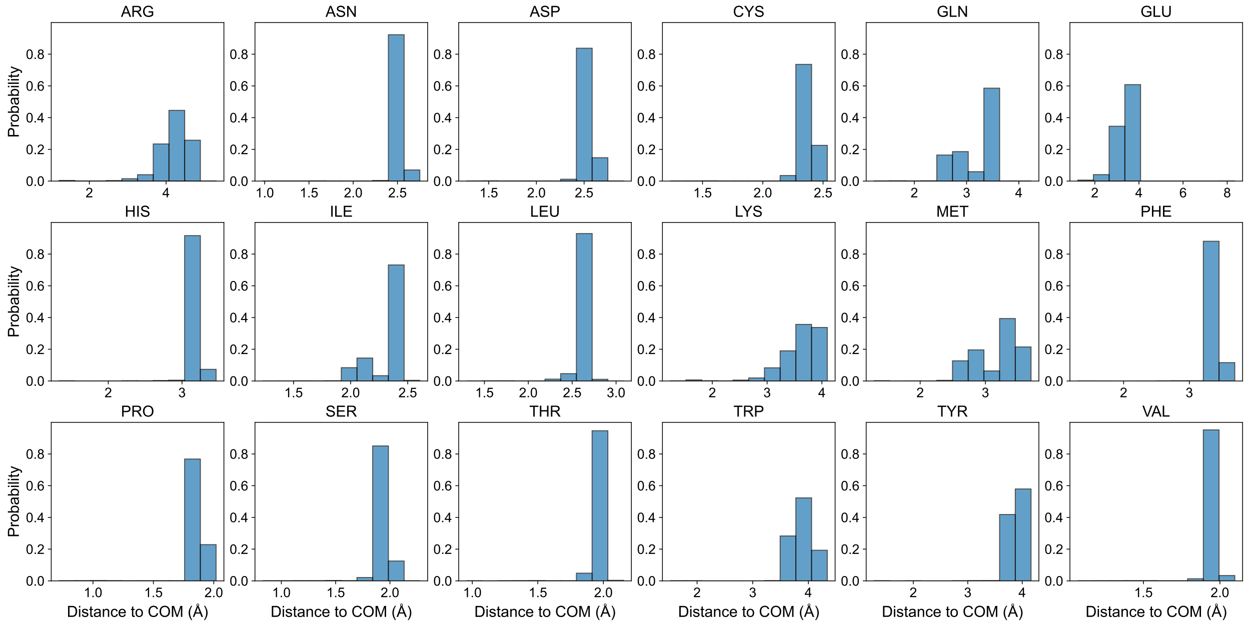


Figure S3: **Side chain distance distributions.** Distribution of C$\alpha$ to side chain centre of mass distances for each residue type, determined empirically from our diverse set of proteins. Glycine and alanine (not shown) are represented by single scattering centres at their C$\alpha$ positions. Average distances given in Table S1.

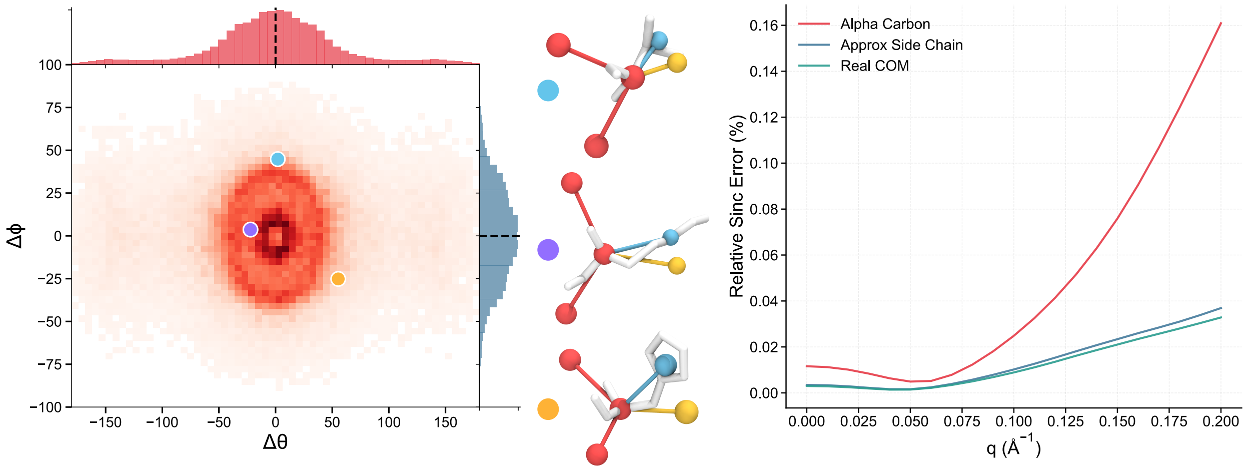


Figure S4: **Geometric side chain placement validation.** (Left) Heatmap showing angular deviations between geometric normal vectors and true centre-of-mass directions across all residue types in the training dataset. (Centre) Representative examples showing C$\alpha$ positions (COM, red), true atomic centres of mass (blue), and geometric predictions (yellow) for three diverse residue types. (Right) Relative error in the sinc function $\sin(qr_{ij})/(qr_{ij})$ comparing C$\alpha$-only, true COM, and geometric COM models against all-atom structures across the validation dataset.

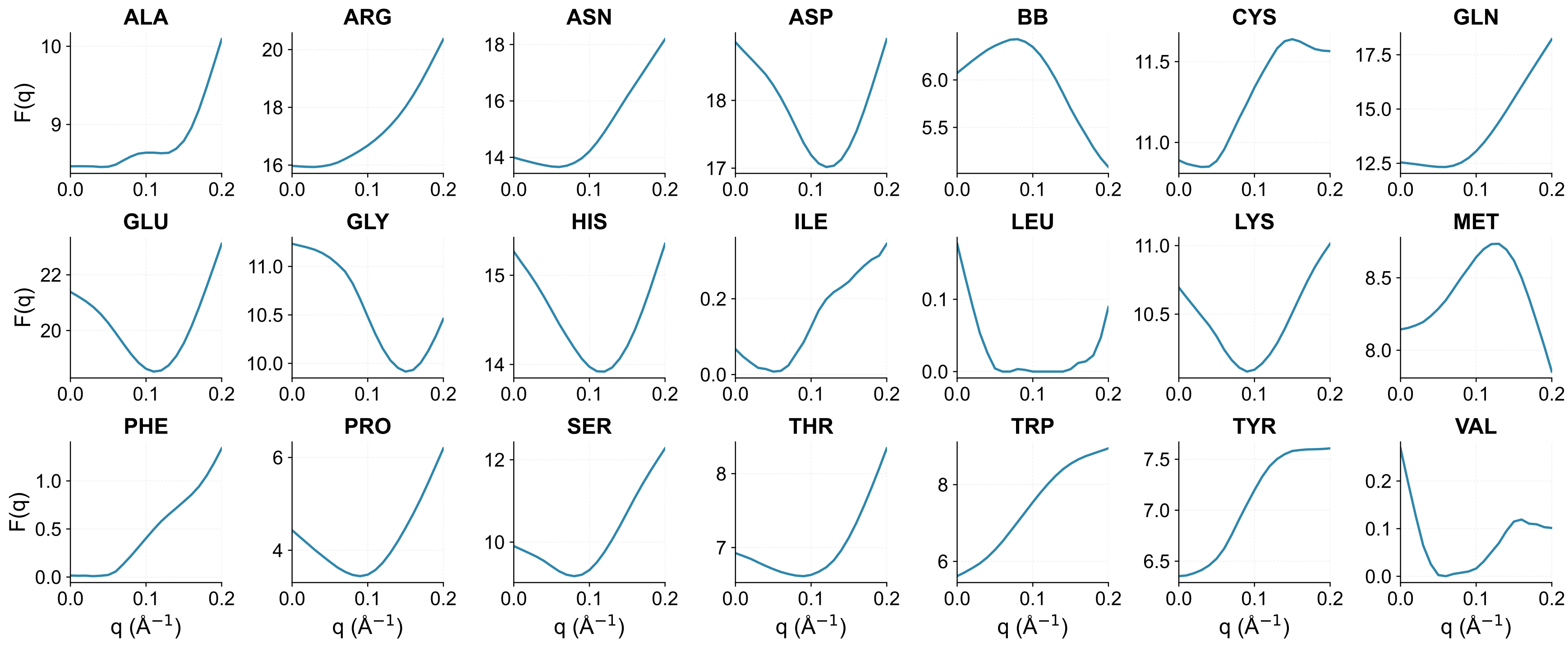


Figure S5: **Optimised form factors for the two-body scattering model.** Form factor profiles $F(q)$ for all 21 scattering centre types as a function of the scattering angle $q$.

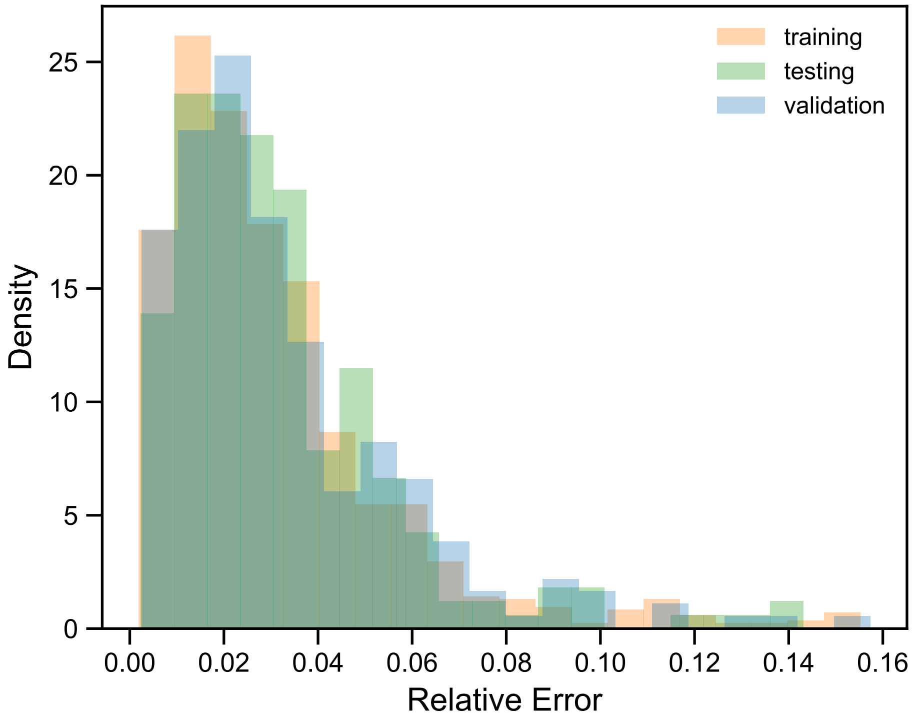


Figure S6: **Relative error distributions in model fitting.** Distribution of relative errors of scattering prediction compared to WAXSiS ground truths in training, testing and validation dataset splits. We observe a similar distribution in the validation dataset, indicating a robust generalisable SAXS prediction model.

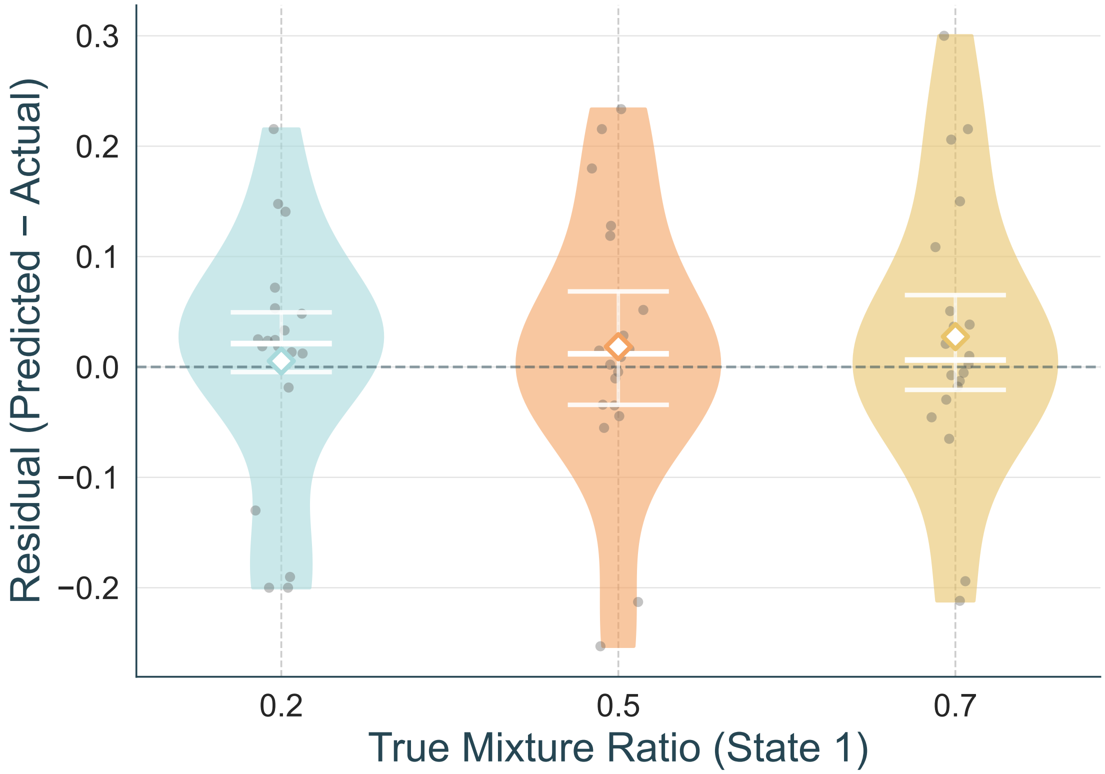


Figure S7: **Residuals of conformation mixture prediction via Wiggle.** Distribution of residuals (predicted - actual) for mixture recovery validation across synthetic two-state protein mixtures at ratios 0.2:0.8, 0.5:0.5, and 0.7:0.3 (state 1 : state 2). Violin plots show the residual distributions, with white horizontal lines indicating quartiles and white diamonds showing mean values. Individual protein systems are shown as gray points. The coarse-grained model achieves consistent accuracy across all mixture ratios (MAE = 0.079, 0.083, 0.086 for ratios 0.2, 0.5, and 0.7 respectively), with no systematic bias toward over- or under-prediction ($\mu = 0.005, 0.018, 0.027$; $\sigma = 0.11, 0.12, 0.12$)

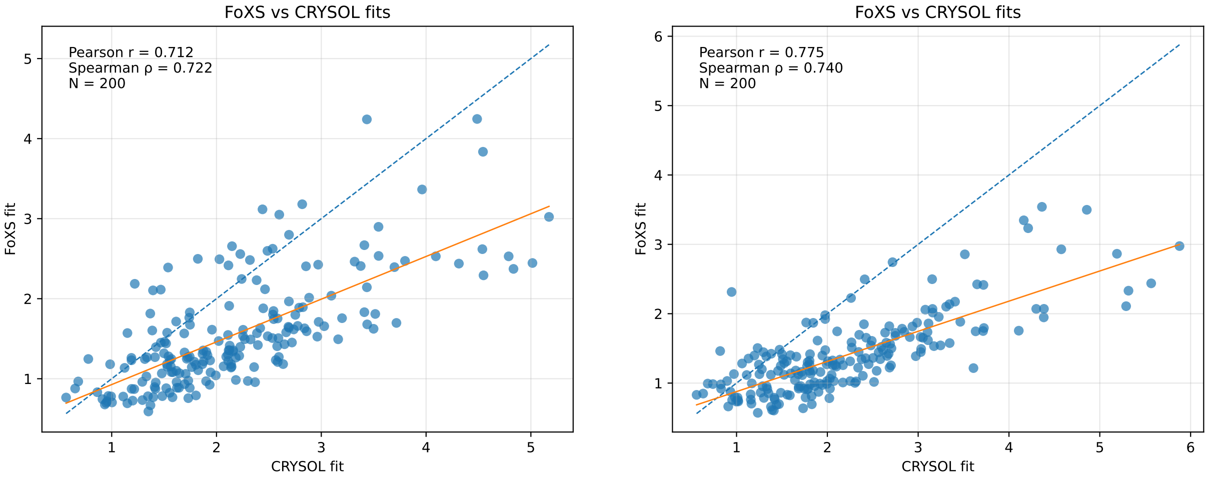


(a) $q \in [0, 0.25]$ (b) $q \in [0, 0.2]$

Figure S8: **Comparison of model fitting to SMARCAL1$^{CD}$ experimental SAXS data using FoXS and CRYSOL**. The two packages are used to predict the SAXS signal of all-atom SMARCAL1$^{CD}$ models based on Carbonara predictions, with the default setting $q_{\max} = 0.2$. Panel (a) shows the $\chi^2$ values for fits to the full data range $q \in [0, 0.25]$ using both packages. The least-squares best-fit line is shown in orange (dashed blue indicates equality). Panel (b) shows the same comparison for fits restricted to $q \in [0, 0.2]$.

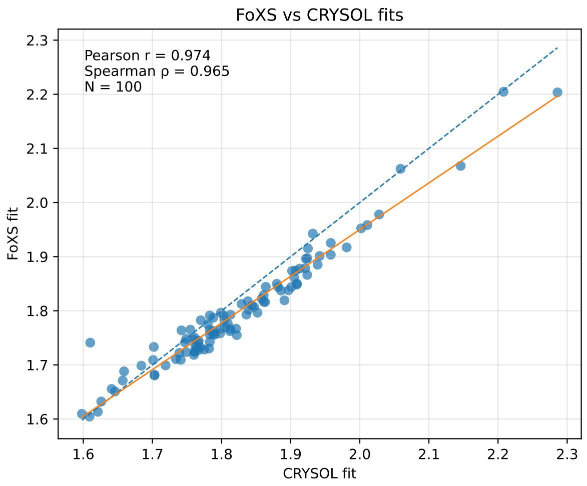


(a) $q \in [0, 0.37]$

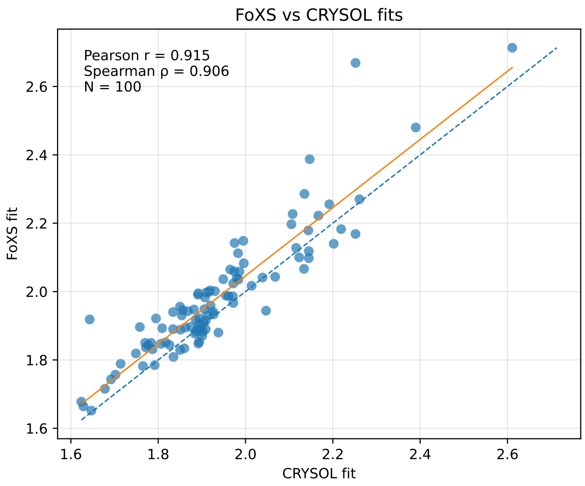


(b) $q \in [0, 0.2]$

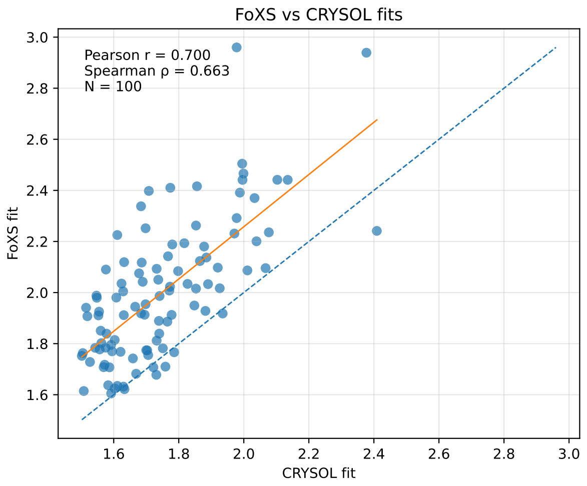


(c) $q \in [0, 0.15]$

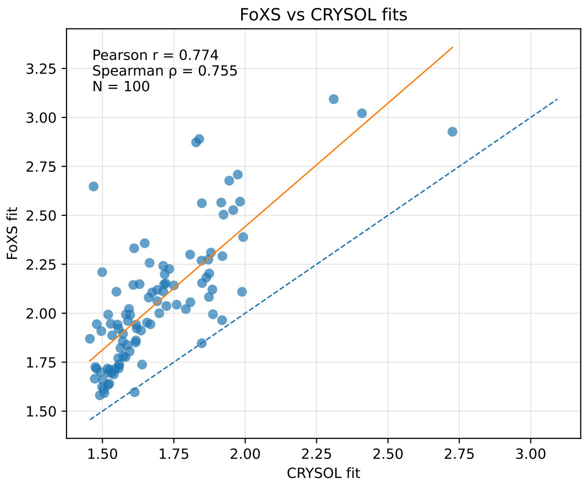


(d) $q \in [0, 0.1]$

Figure S9: **Comparison of model fitting to ChiLob7/4 experimental SAXS data using FoXS and CRYSOL**. The two packages are used to predict the SAXS signal of all-atom ChiLob7/4 based on Carbonara predictions. Panel (a) shows the $\chi^2$ values for fits to the full data range $q \in [0, 0.37]$ using both packages, based on Carbonara predictions with the default setting $q_{\max} = 0.2$. The least-squares best-fit line is shown in orange (dashed blue indicates equality). Panel (b) shows the $\chi^2$ values for fits to the full data range $q \in [0, 0.37]$ using both packages, based on Carbonara predictions with the default setting $q_{\max} = 0.2$. Panel (c) shows the $\chi^2$ values for fits to the full data range $q \in [0, 0.15]$ using both packages, based on Carbonara predictions with the default setting $q_{\max} = 0.15$. Panel (d) shows the $\chi^2$ values for fits to the full data range $q \in [0, 0.1]$ using both packages, based on Carbonara predictions with the default setting $q_{\max} = 0.1$.

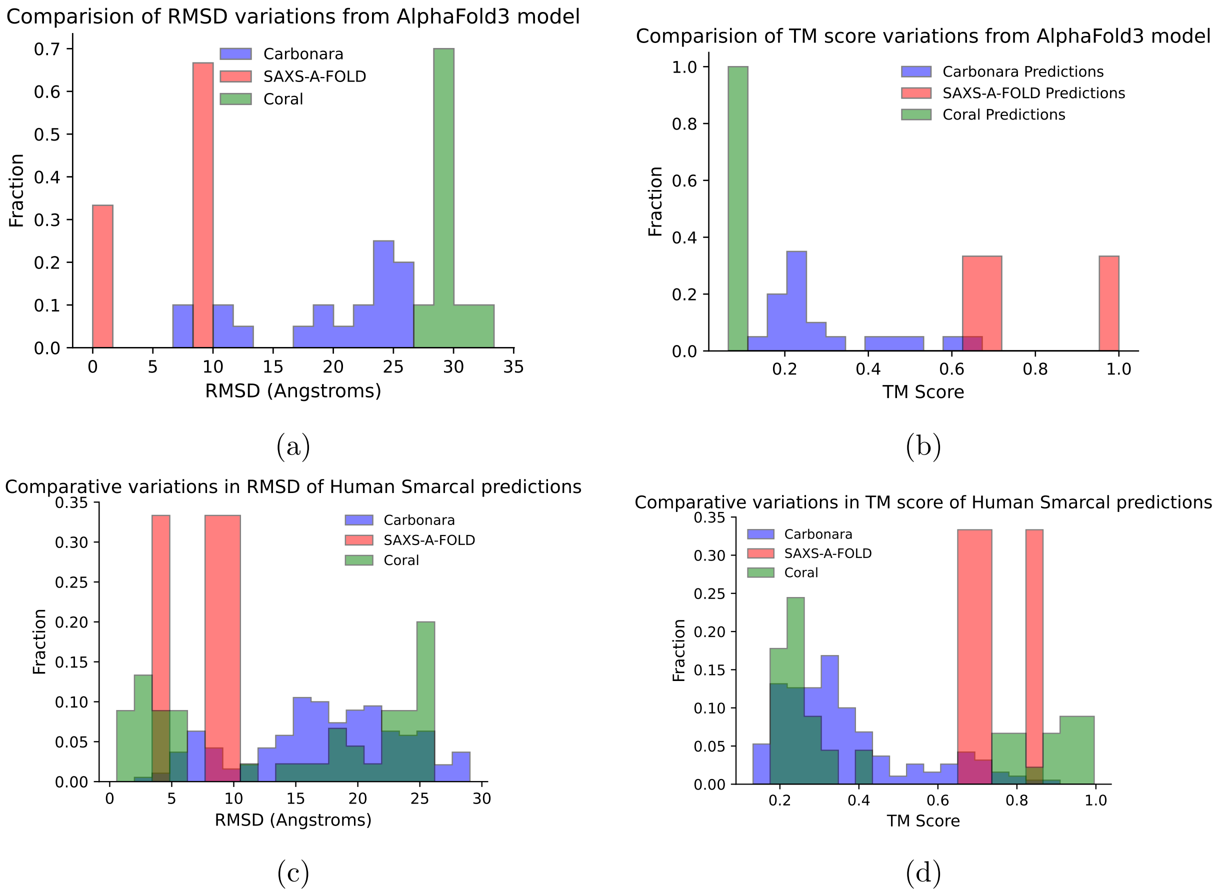


Figure S10: **Comparisons of TM and RMSD scores for Human SMARCAL1$^{CD}$ predictions made using Carbonara, SAXS-A-FOLD and CORAL**. Panels (a)(RMSD) and (b)(TM) illustrate the variation of predictions with respect to the original AlphaFold3 model. Panels (c) and (d) illustrate the variation between predictions in terms of RMSD and TM-score, respectively.

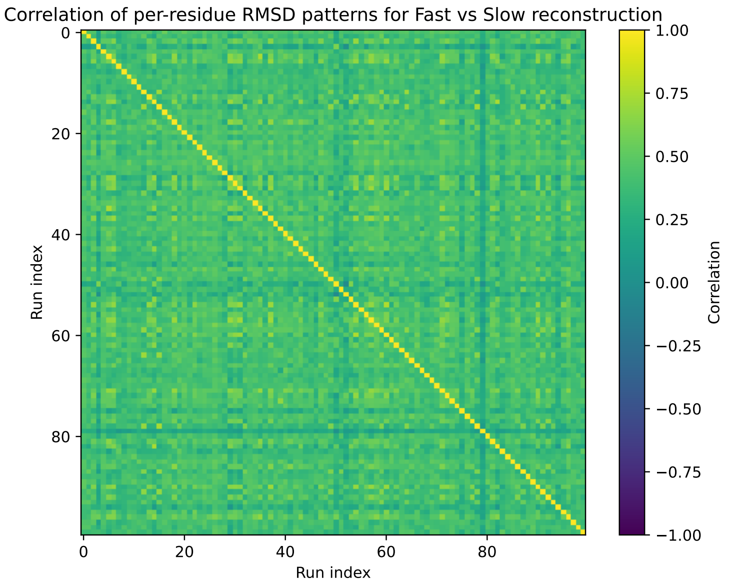


(a)

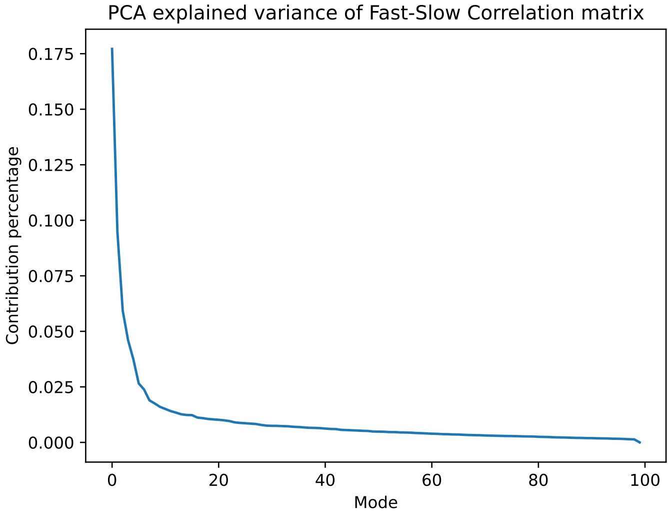


(b)

Figure S11: **Correlation of Fast vs Slow all-atom backmapping protocols**. Panel (a) shows the per-residue RMSD correlation matrix for both methods a cross the 100 predictions for Human SMARCAL1$^{CD}$(final state). Panel (b) shows the PCA contribution as a function of mode for the matrix in (a). See Supplementary Methods 1.7.

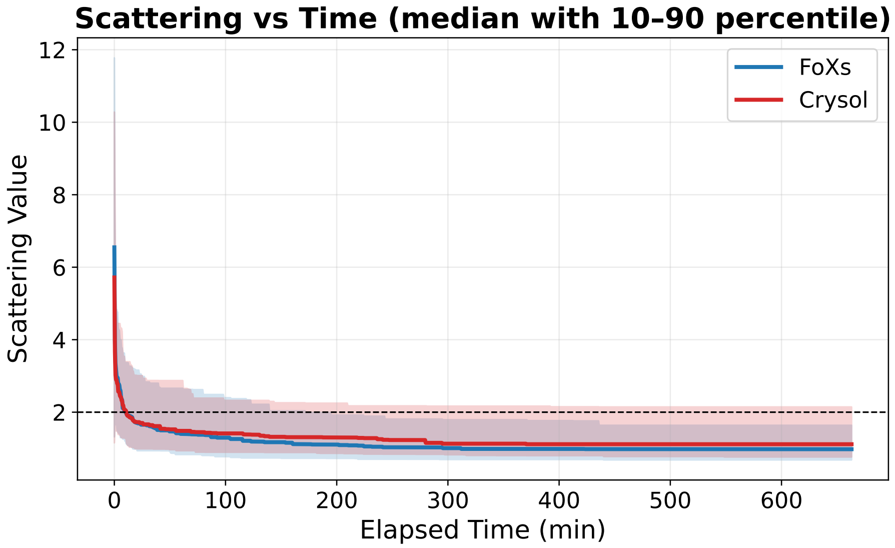


(a)

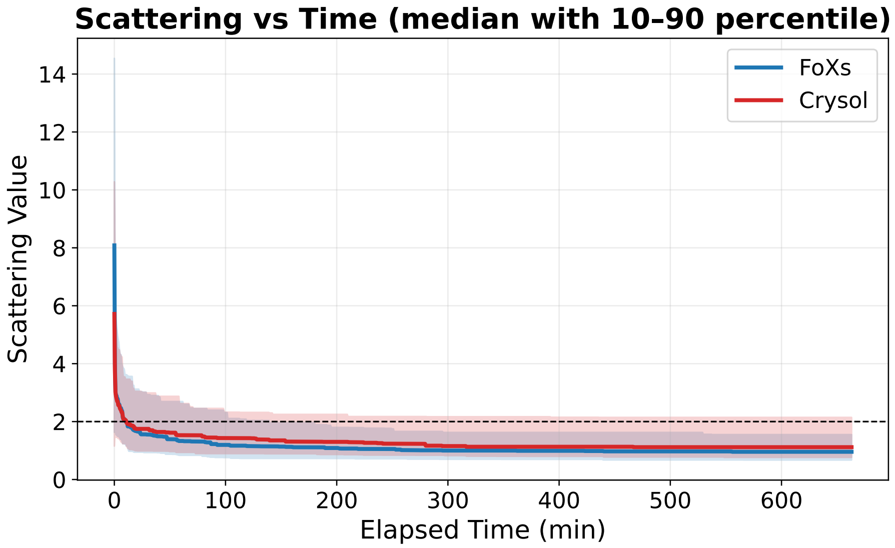


(b)

Figure S12: **Minimum $\chi^2$ values for FoXS and CRYSOL as a function of runtime**. The solid lines show mean values, and the coloured band the 10-90 percentile window over 100 runs. The horizontal dashed line shows where the curves fall below 2. In panel (a), results over the full scattering range, in panel (b) in the 0-0.2 Å$^{-1}$ region.

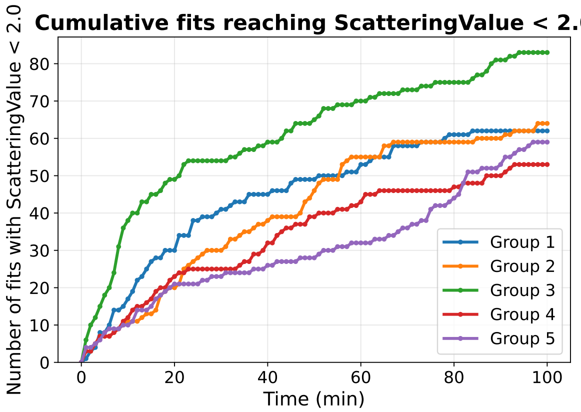


(a) FoXS $\chi^2 < 2$

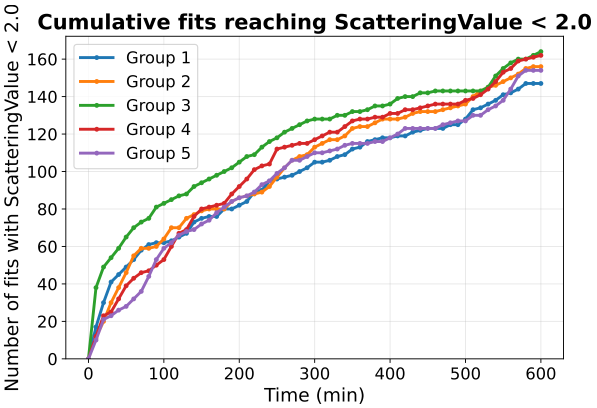


(b) FoXS $\chi^2 < 2$, full runtime

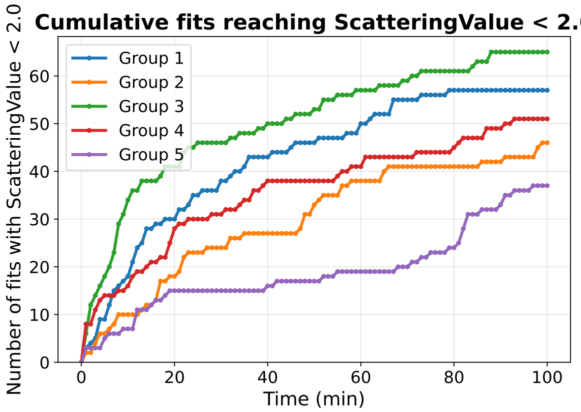


(c) CRYSOL $\chi^2 < 2$

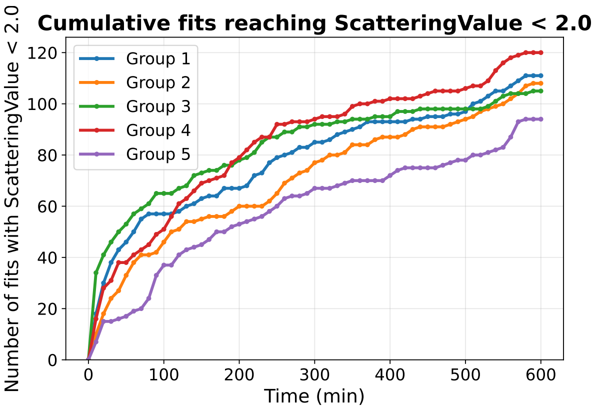


(d) CRYSOL $\chi^2 < 2$, full runtime

Figure S13: **Quantification of successful ($\chi^2 < 2$) SMARCAL1$^{CD}$ models produced at runtime**. Number of all-atom Human SMARCAL1$^{CD}$ predictions obtained, up to a given time, with a fit quality $\chi^2 < 2$ as assessed by FoXS and CRYSOL on the full data range $q \in [0, 0.25]$. The five plots in each panel correspond to the 5 batch runs of twenty refinements. Panels (a) and (b) are for the FoXS assessment over time periods $[0, 100]$ and $[0, 600]$ minutes respectively. Panels (c) and (d) are for CRYSOL over the same time periods.

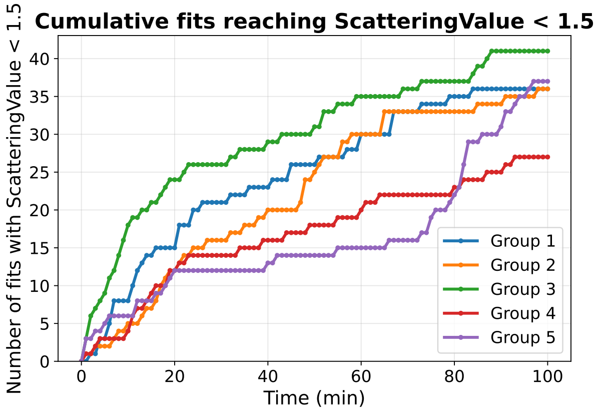


(a) FoXs $\chi^2 < 1.5$

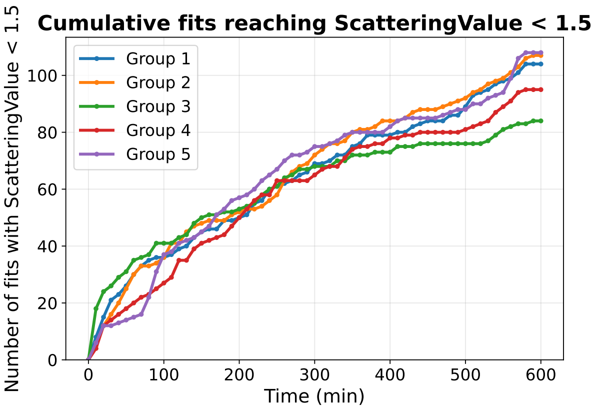


(b) FoXs $\chi^2 < 1.5$, full runtime

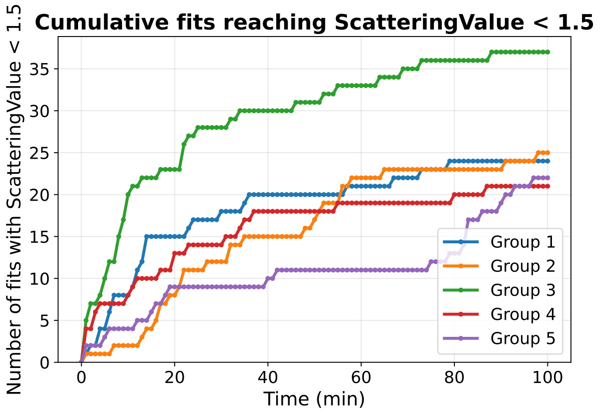


(c) CRYSOL $\chi^2 < 1.5$

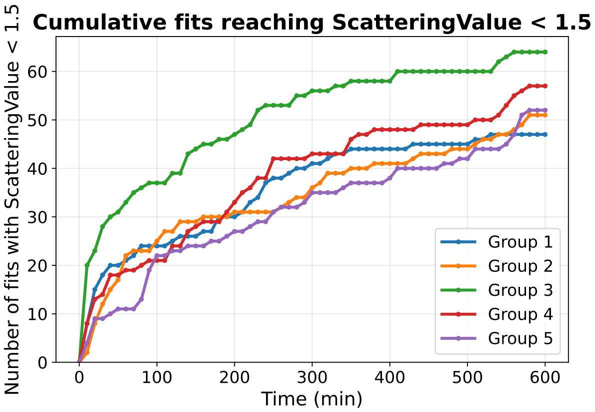


(d) CRYSOL $\chi^2 < 1.5$, full runtime

Figure S14: **Quantification of successful ($\chi^2 < 1.5$) SMARCAL1$^{CD}$ models produced at runtime**. Number of all-atom Human SMARCAL1$^{CD}$ predictions obtained, up to a given time, with a fit quality $\chi^2 < 1.5$ as assessed by FoXS and CRYSOL on the full data range $q \in [0, 0.25]$. The five plots in each panel correspond to the 5 batch runs of twenty refinements. Panels (a) and (b) are for the FoXS assessment over time periods $[0, 100]$ and $[0, 600]$ minutes respectively. Panels (c) and (d) are for CRYSOL over the same time periods.

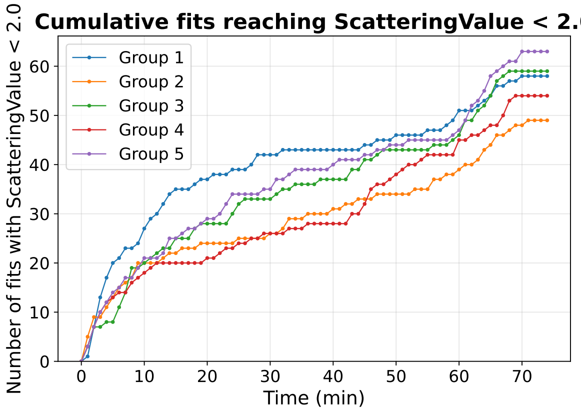


(a) FoXS $\chi^2 < 2$

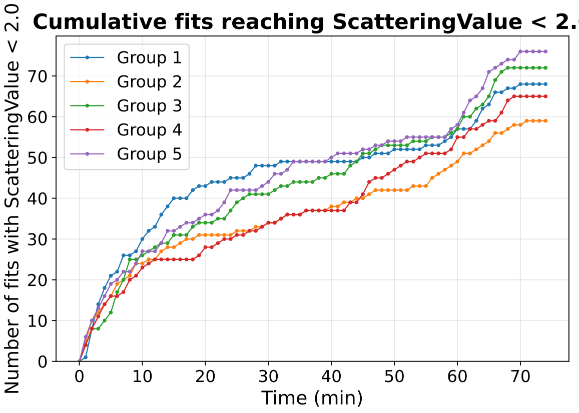


(b) CRYSOL $\chi^2 < 2$, full runtime

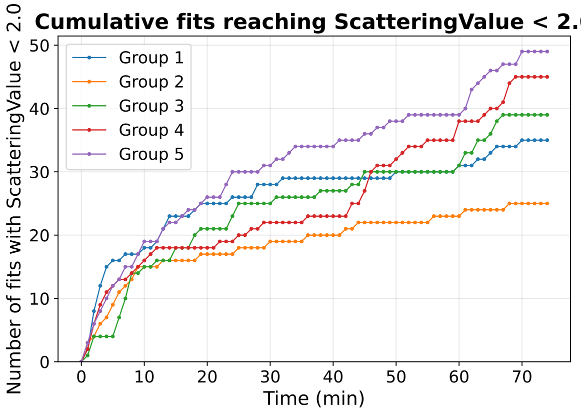


(c) FoXS $\chi^2 < 2$

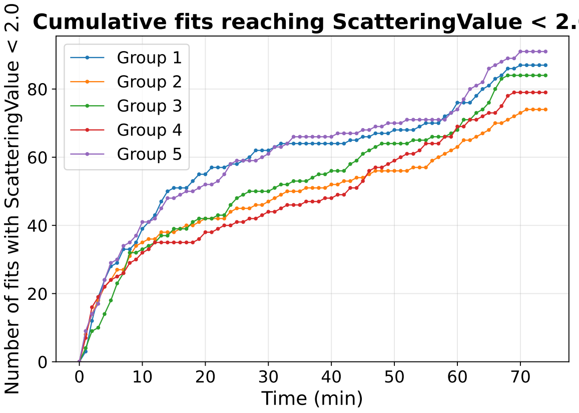


(d) CRYSOL $\chi^2 < 2$, full runtime

Figure S15: **Quantification of successful ($\chi^2 < 2$) ChiLob 7/4 models produced at runtime** Number of all-atom IGg2 predictions obtained, up to a given time, with a fit quality $\chi^2 < 2$ as assessed by FoXS and CRYSOL on the full data range $q \in [0, 0.37]$ and $q \in [0, 0.1]$, the range over which the most substantial improvement from the original homology model was obtained. The five plots in each panel correspond to the 5 batch runs of twenty refinements. Panel (a) show the FoXS assessments on $q \in [0, 0.37]$, panel (b) CRYSOL on $q \in [0, 0.37]$, panel (c) FoXS assessments on $q \in [0, 0.1]$, panel (b) CRYSOL on $q \in [0, 0.1]$.

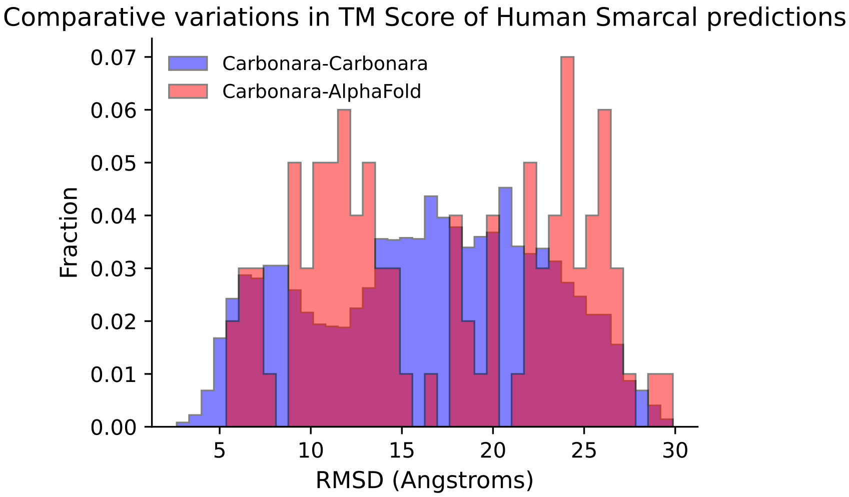


(a)

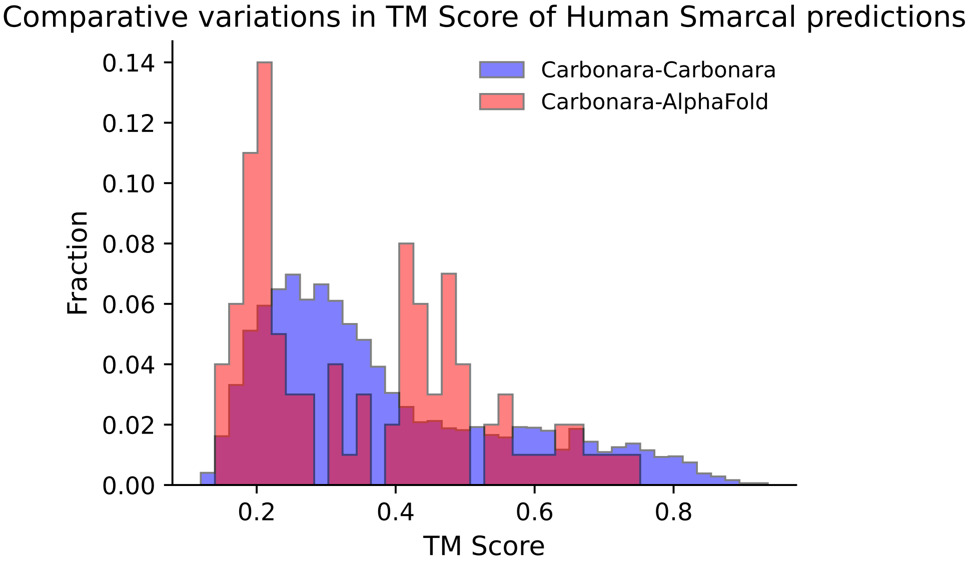


(b)

Figure S16: **Structural variation of SMARCAL1$^{CD}$ Carbonara predictions**. Panel (a) shows the RMSD scores for prediction-prediction and prediction-AlphaFold comparisons and panel (b) the TM scores of models for $q_c^{max} = 0.2$.

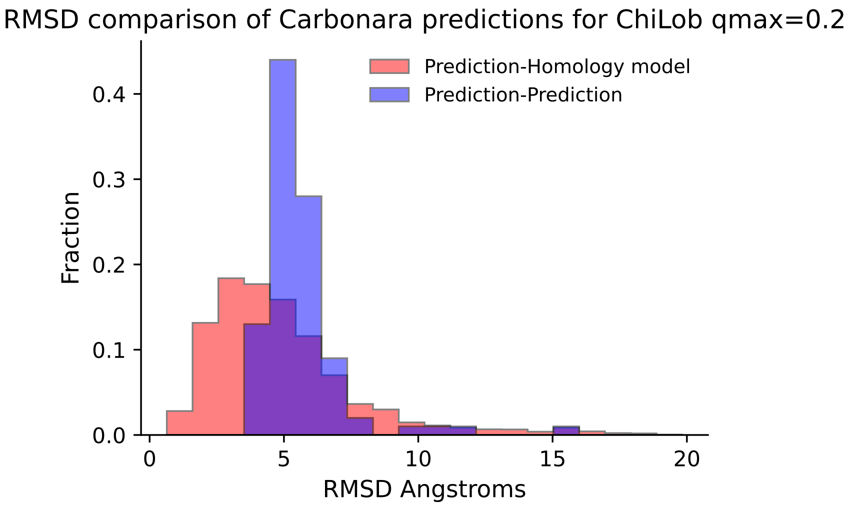


(a)

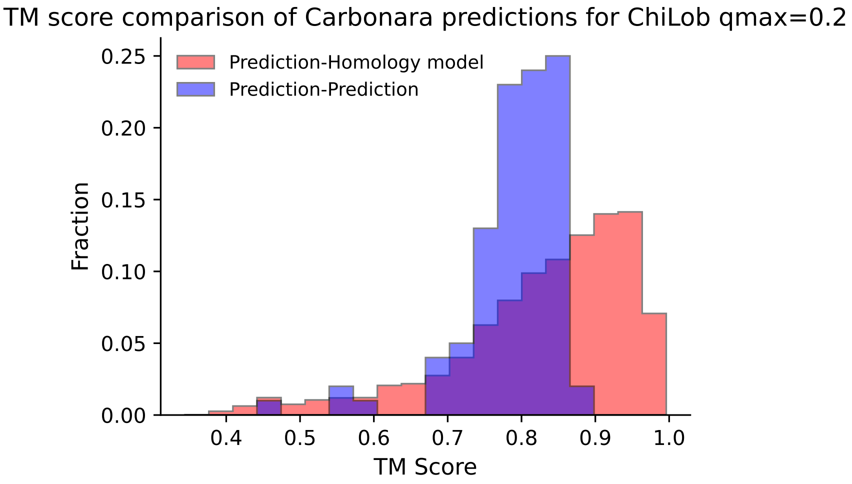


(b)

Figure S17: **Structural variation of ChiLob 7/4 Carbonara predictions**. Histograms comparing the structural variation of Carbonara predictions for $q_c^{max} = 0.2$. Panel (a) shows the RMSD scores for prediction-prediction and prediction-AlphaFold comparisons, panel (b) the TM scores.

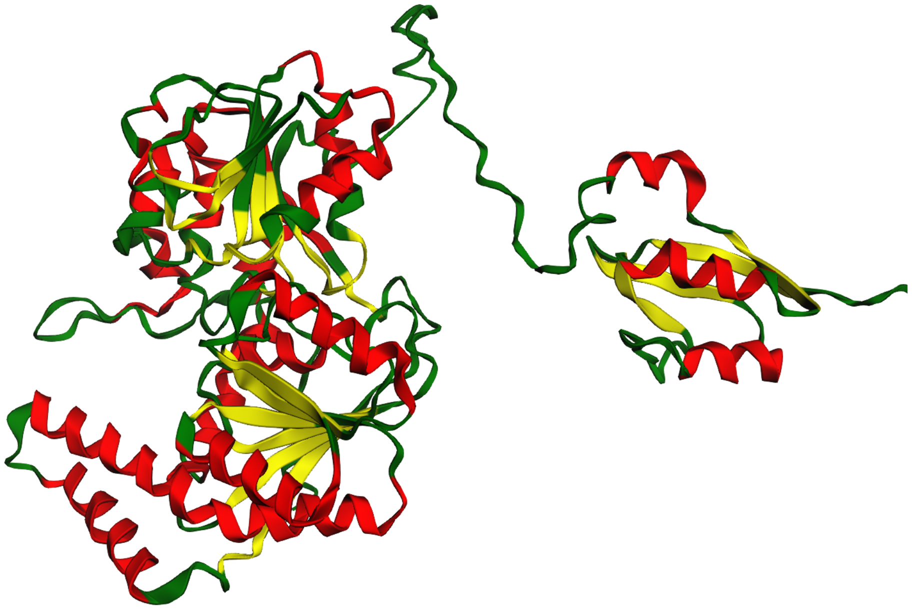

(a) CORAL model

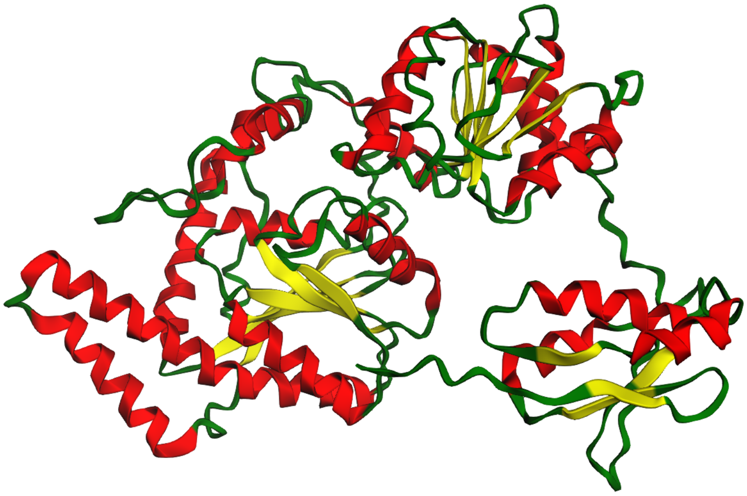

(b) Carbonara model

Figure S18: **Representative SMARCAL1$^{CD}$ models from CORAL and Carbonara**. The Carbonara model is more compact than the CORAL one. The CORAL model is produced via rigid body motions around a dummy linker section, an approach that can lead to artificially extended conformations.

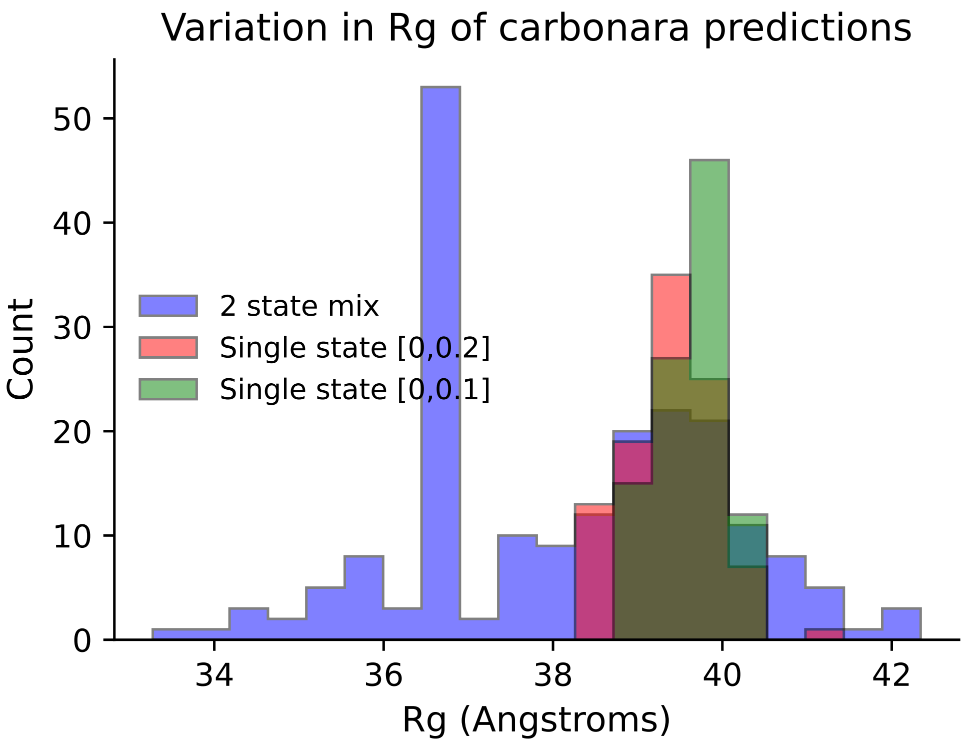


(a)

Figure S19: **Radius of Gyration ($R_g$) of Carbonara IgG2 predictions**. We analyse batch runs completed with a mixture prediction of two states with $q^c_{max} = 0.2$ and for comparison single state prediction sets with $q^c_{max} = 0.2$ and $q^c_{max} = 0.1$.

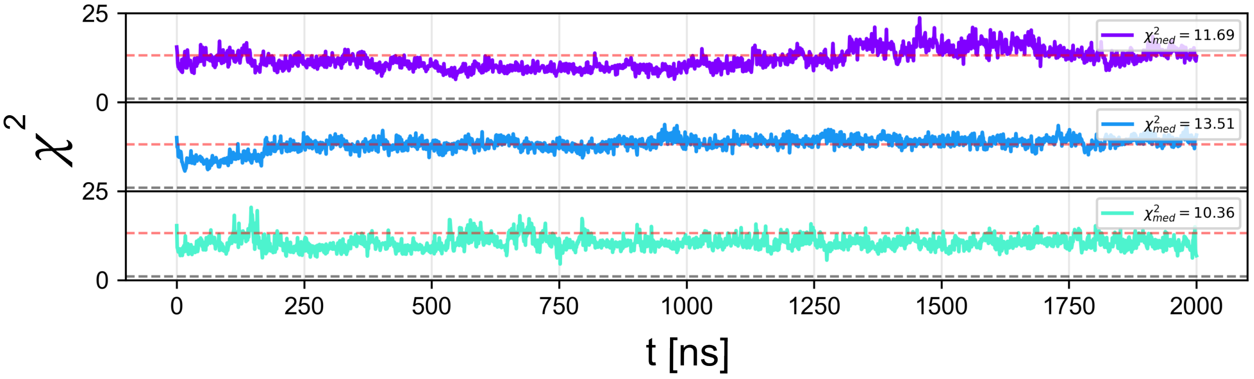


Figure S20: $\chi^2$ **over each 2 $\mu$s simulation of the AlphaFold SMARCAL1$^{CD}$ structure**. Median $\chi^2$ values shown in top right of each plot. Red dashed line indicates the average $\chi^2$ value over all simulations seeded from the AlphaFold model.

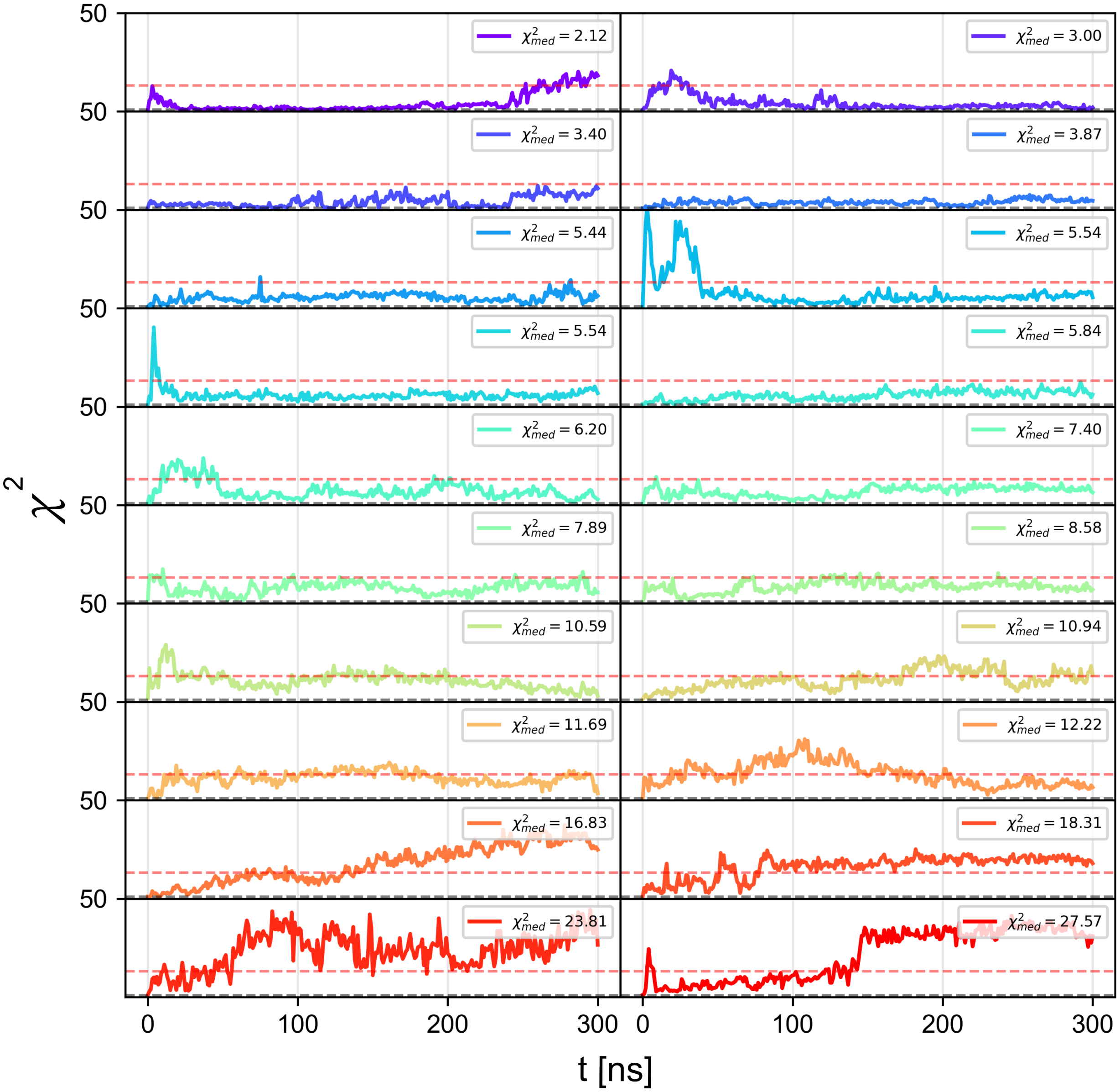


Figure S21: $\chi^2$ **over each 300 ns simulation of each of the 20 Carbonara refined SMARCAL1$^{CD}$ structures**. Structures sorted by median $\chi^2$ values, shown in top right of each plot. Red dashed line indicates the average $\chi^2$ value over all simulations seeded from the AlphaFold model.

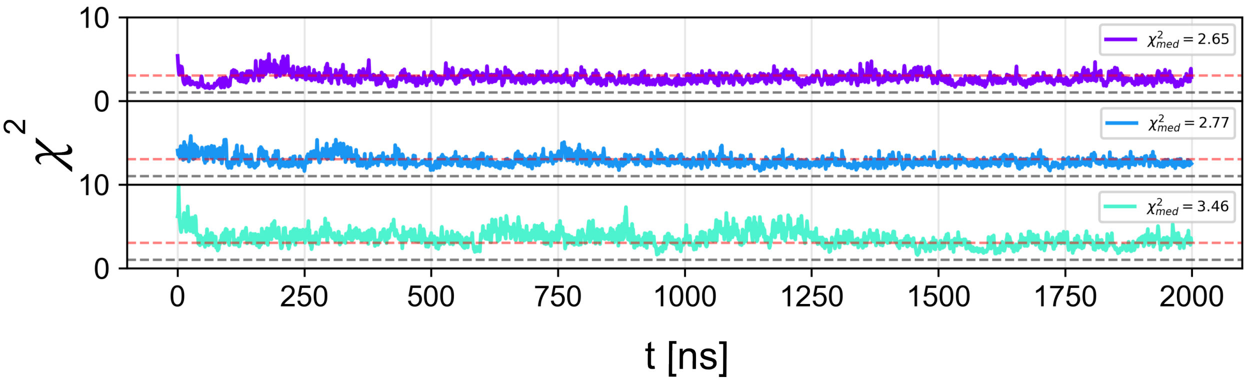


Figure S22: $\chi^2$ **over each 2000 ns simulation of the Crystal IgG2 structure**. Median $\chi^2$ values shown in top right of each plot. Red dashed line indicates the average $\chi^2$ value over all Crystal seeded IgG2 simulations

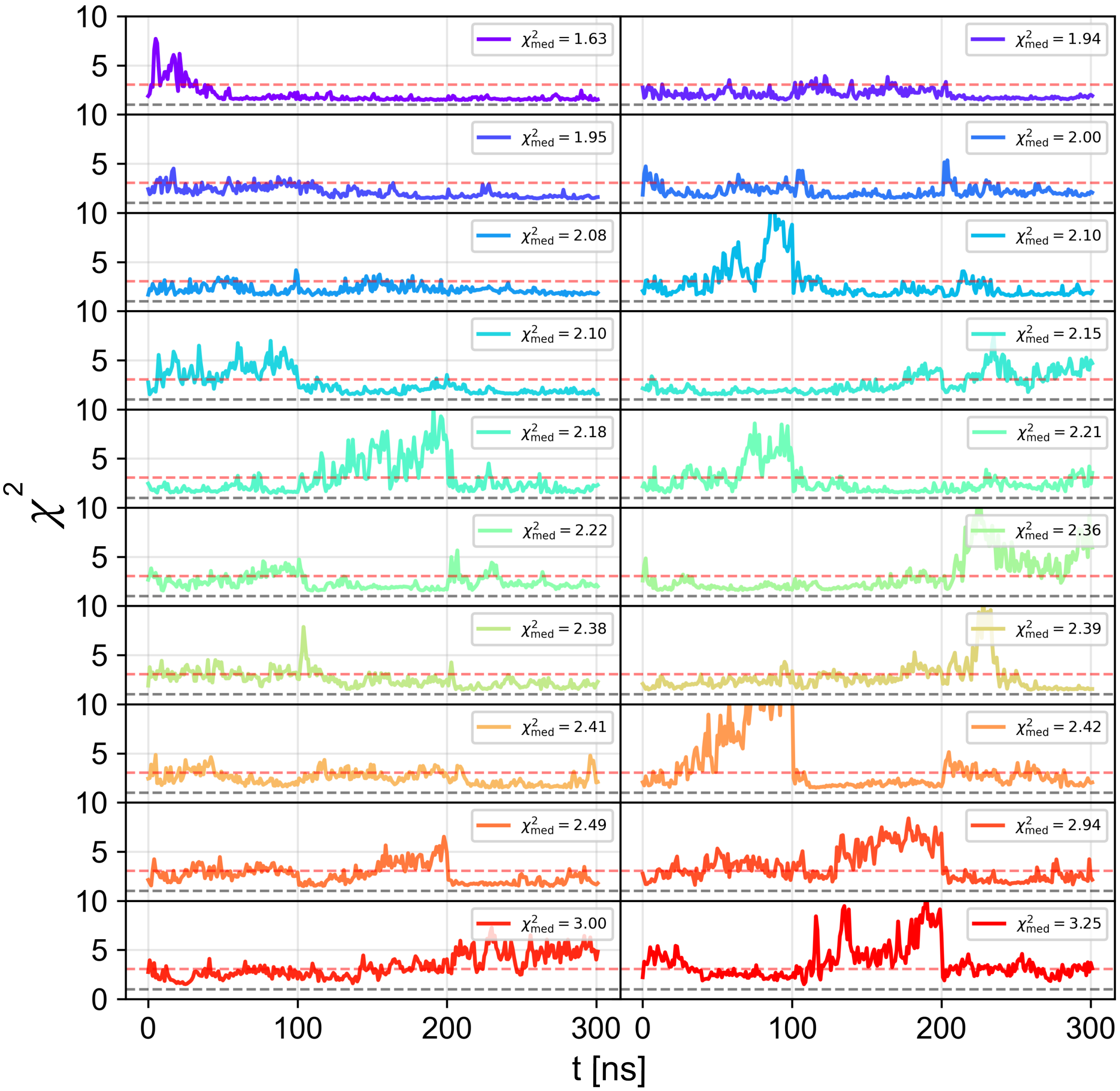


Figure S23: $\chi^2$ **over each 300 ns simulation of each of the 20 Carbonara refined IgG2 structures**. Structures sorted by median $\chi^2$ values, shown in top right of each plot. Red dashed line indicates the average $\chi^2$ value over all simulations seeded from the IgG2 crystal structure.

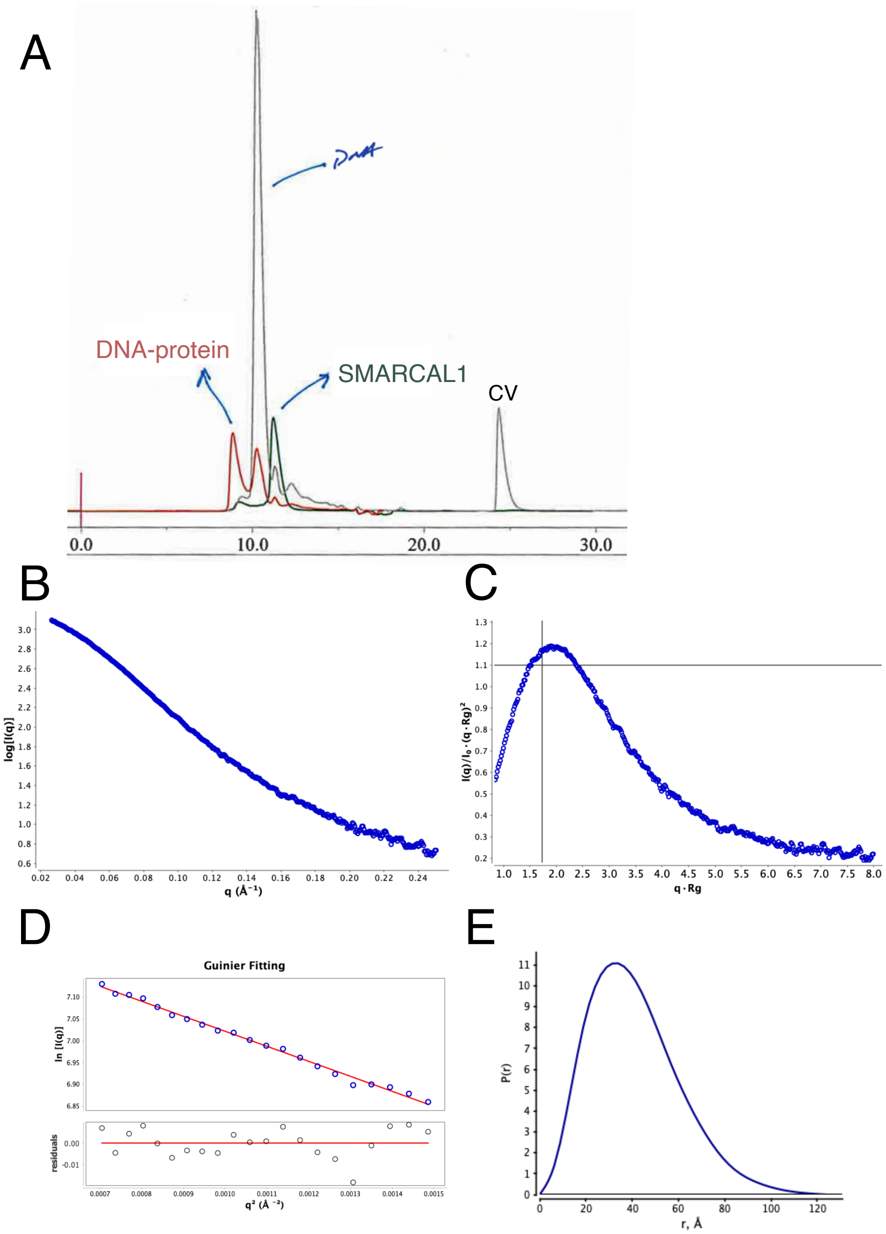


Figure S24: SMARCAL1$^{CD}$ purification and SAXS analysis. A) SMARCAL1$^{CD}$ purified on Shodex KW-402 column as described in,[28] green trace. Single, symmetric elution peak corresponding to SMARCAL1$^{CD}$ was fractioned at peak elution ( $60uL$) and taken for SAXS measurements. SAXS data was collected as a 2/3rds dilution series over 4 successive dilutions. B) Merged SAXS curve of SMARCAL1$^{CD}$ producing an $R_g$ of 34.76 Å Porod Volume of 130,000 Å$^3$ and a volume-of-correlation molecular mass of 67 kDa (actual 65 kDa). C) Normalized Kratky plot suggesting a compact, non-globular particle. D) Guinier region with corresponding residuals. E) Real-space, distance distribution function suggesting maximum dimension of 118 Å$^{\cdot}$

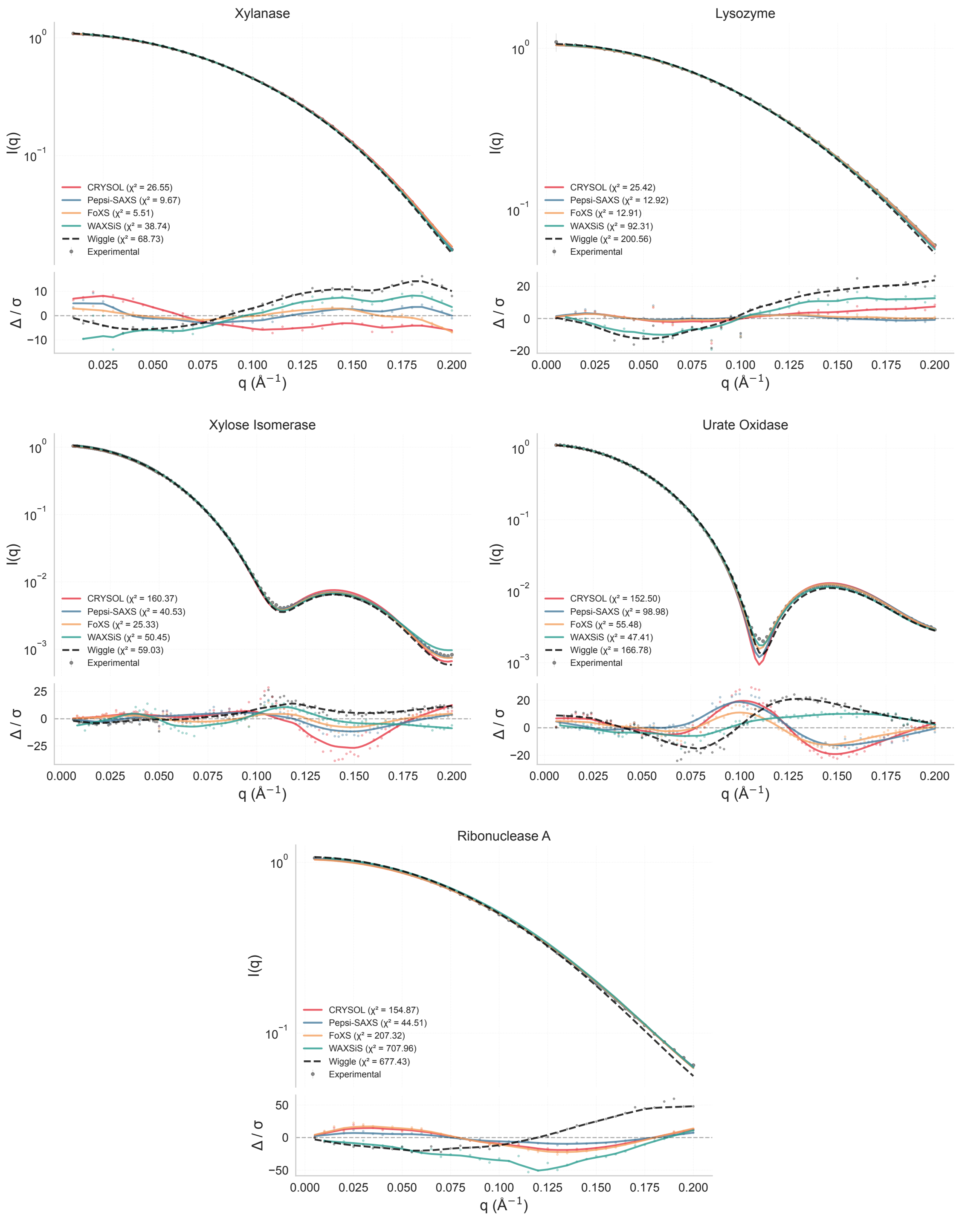


Figure S25: **Comparison of SAXS prediction methods on experimental benchmark data.** Scattering intensity profiles I(q) (top panels) and error-weighted residuals (bottom panels) for five proteins from the round-robin consensus dataset.[29] Experimental data (gray points) are compared with predictions from crystal structures by CRYSOL (red), Pepsi-SAXS (blue), FoXS (green), WAXSiS (orange), and Wiggle (black dashed). All methods capture the overall profile shapes and characteristic features. The large error-weighted residuals reflect the exceptionally small experimental uncertainties in the consensus profiles. Predictions from CRYSOL, Pepsi-SAXS, FoXS, and WAXSiS as reported in the original study, Wiggle predictions calculated using identical crystal structures and q-range.
